# ChiS is a noncanonical DNA-binding hybrid sensor kinase that directly regulates the chitin utilization program in *Vibrio cholerae*

**DOI:** 10.1101/2020.01.10.902320

**Authors:** Catherine A. Klancher, Shouji Yamamoto, Triana N. Dalia, Ankur B. Dalia

## Abstract

<u>T</u>wo-<u>c</u>omponent signal transduction <u>s</u>ystems (TCSs) represent a major mechanism that bacteria use to sense and respond to their environment. Prototypical TCSs are composed of a membrane-embedded histidine kinase (HK), which senses an environmental stimulus and subsequently phosphorylates a cognate partner protein called a response regulator (RR) that regulates gene expression in a phosphorylation-dependent manner. *Vibrio cholerae* uses the hybrid HK ChiS to activate the expression of the chitin utilization program, which is critical for the survival of this facultative pathogen in its aquatic reservoir. A cognate RR for ChiS has not been identified and the mechanism of ChiS-dependent signal transduction remains unclear. Here, we show that ChiS is a noncanonical membrane-embedded one-component system that can both sense chitin and directly regulate gene expression via a cryptic DNA binding domain. Unlike prototypical TCSs, we find that ChiS DNA binding is diminished, rather than stimulated, by phosphorylation. Finally, we provide evidence that ChiS likely activates gene expression by directly recruiting RNA polymerase. Together, this work addresses the mechanism of action for a major transcription factor in *V. cholerae* and highlights the versatility of signal transduction systems in bacterial species.

**Significance Statement:** From bacteria to humans, the ability to properly respond to environmental cues is critical for survival. The cholera pathogen *Vibrio cholerae* uses one protein, ChiS, to sense chitin in its environmental reservoir to regulate the expression of genes that are critical for the survival and evolution of this pathogen in this niche. Here, we study how the chitin sensor ChiS works, and discover that it regulates gene expression in an unexpected and unorthodox manner. Thus, this study uncovers how the major regulator ChiS works in this important human pathogen and highlights the versatile mechanisms that living systems use to respond to their environment.

## Introduction

*Vibrio cholerae*, the bacterium responsible for the diarrheal disease cholera, is naturally found in the marine environment where it forms biofilms on the chitinous shells of crustacean zooplankton (1). Chitin is a major nutrient source for *V. cholerae* in this niche. In addition to serving as a nutrient source, chitin is also used as a cue to induce horizontal gene transfer by natural transformation. Furthermore, the formation of chitin biofilms promotes the waterborne transmission of cholera in endemic areas (2, 3). Thus, *V. cholerae*-chitin interactions are critical for the survival, evolution, and transmission of this pathogen.

*V. cholerae* senses chitin via the hybrid sensor kinase ChiS to activate the expression of the chitin utilization program (4, 5). Hybrid sensor kinases are a member of the two-component system (TCS) family of proteins. Prototypical TCSs consist of a membrane-embedded histidine kinase (HK) and a cytoplasmic partner protein called a response regulator (RR) (6). In response to an environmental stimulus, the HK autophosphorylates a conserved histidine. This phosphate is then transferred to a conserved aspartate on the receiver (Rec) domain of its cognate RR. The output activity of the RR, which is DNA-binding in prototypical systems, is enhanced upon phosphorylation. This leads to altered gene expression is response to the upstream cue sensed by the HK. Hybrid sensor kinases, like ChiS, contain additional domains (Rec and/or histidine phosphotransfer domains) that increase the number of steps in the phosphorelay that leads to phosphorylation of their cognate RR. ChiS contains both HK and Rec domains, including their conserved phosphorylation sites (H469 and D772, respectively) (**Fig. S1**). Though it was discovered ∼15 years ago that chitin induces the chitin utilization program of *Vibrio cholerae* through the hybrid HK ChiS, the mechanism of action for this regulator has remained unclear. Here, we show that ChiS does not have a cognate RR, but rather acts as a one-component system that can both sense chitin and directly regulate gene expression from the membrane.

## Results

### Phosphorylation of the ChiS receiver domain inhibits Pchb activation

ChiS is required for activation of the chitin utilization program (5). To study ChiS activity, most studies employ the chitobiose utilization operon (*chb*), which is highly induced in the presence of chitin oligosaccharides and required for the uptake and catabolism of the chitin disaccharide (4, 7-9). In the absence of chitin, the periplasmic <u>c</u>hitin <u>b</u>inding <u>p</u>rotein (CBP) represses ChiS. Thus, in a *cbp*^+^ strain, ChiS is repressed and unable to mediate activation of P_*chb*_ in the absence of chitin when assessed using a P_*chb*_-GFP transcriptional reporter (**Fig. 1a**) or by directly assessing transcript abundance via qRT-PCR (**Fig. 1b**). ChiS can be activated to induce P_*chb*_ expression genetically by deleting *cbp* or by culturing cells in the presence of soluble chitin oligosaccharides (4, 7) (**Fig. 1a-b**).

**Figure 1.**
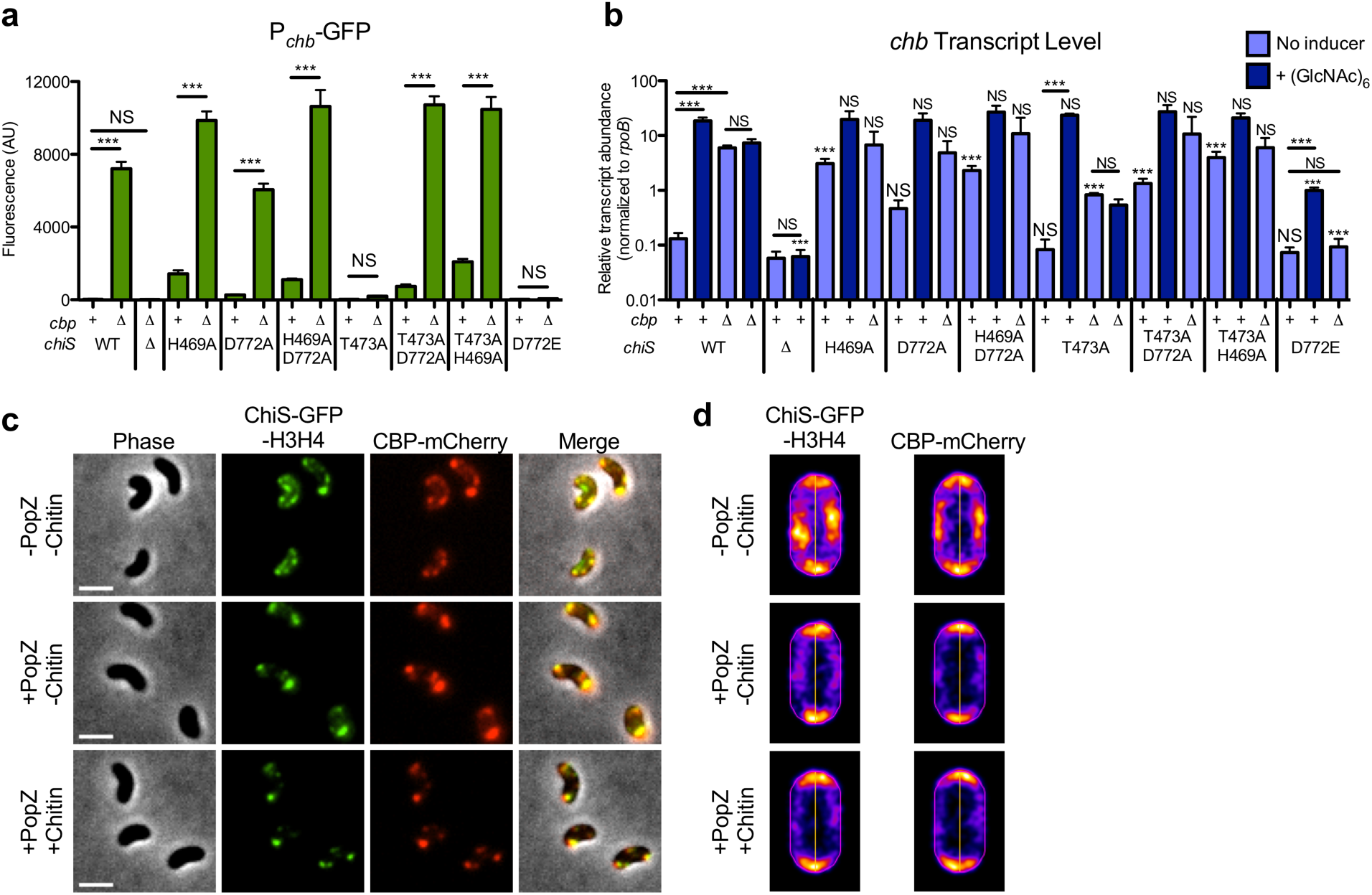
ChiS activity is regulated by CBP and the phosphorylation status of its receiver domain. Point mutants of ChiS in the conserved sites of phosphorylation (H469, D772) or a residue required for phosphatase activity (T473) were assessed for activation of P_*chb*_ using (**a**) a P_*chb*_-GFP transcriptional reporter using deletion of *cbp* as an inducer or (**b**) by assessing *chb* transcript abundance by qRT-PCR using deletion of *cbp* or chitin hexasaccharide as an inducer. In **a**, strains were grown in LB and either have *cbp* intact (+) or deleted (Δ) as indicated. In **b**, strains were grown in LB only (no inducer) or in LB medium supplemented with chitin hexasaccharide (+ (GlcNAc)_6_); and, strains either have CBP intact (+) or deleted (Δ) as indicated. Statistical markers indicated directly above bars indicate comparisons to the equivalent WT condition (*e.g.* the “*cbp*^+^ +(GlcNAc)_6_” bars for all mutant strains are compared to the “WT *cbp*^+^ +(GlcNAc)_6_” condition). (**c-d**) POLAR assay with a *V. cholerae* strain containing P_*tac*_-ChiS-GFP-H3H4 (H3H4 is a PopZ interaction domain), CBP-mCherry, and P_*BAD*_-PopZ. Scale bars = 2 µm. All cells were cultured in LB supplemented with 10 µM IPTG to induce ChiS-GFP-H3H4 expression. PopZ expression was induced by growing cells with 0.05% arabinose where indicated (+PopZ). Chitin hexasaccharide (0.5%) was added to cells for 30 minutes before imaging where indicated (+Chitin). (**c**) Representative images and (**d**) heat maps indicating the localization of fluorescence in the indicated channels. Heat maps were generated from analysis of at least 500 cells. For a complete kinetic analysis of CBP-mCherry localization in the presence of chitin hexasaccharide, see **Fig. S8**. Data in **a** and **b** are from at least three independent biological replicates and shown as the mean ± SD. Statistical comparisons in **a** and **b** were made by one-way ANOVA with Tukey’s post-test. NS, not significant. ***, p < 0.001. For a detailed list of all statistical comparisons see **Table S1**.

To regulate gene expression in response to an environmental stimulus, HKs initiate a phosphorylation cascade via kinase activity to signal to their cognate RR. Thus, we tested whether the kinase activity and/or phosphorylation of ChiS are required for P_*chb*_ activation. Mutation of either or both phosphorylation sites in ChiS (ChiS^H469A^, ChiS^D772A^, and ChiS^H469A D772A^) still allowed for P_*chb*_ induction when *cbp* was deleted (**Fig. 1a-b**), consistent with prior results (8). This suggests that the kinase activity and phosphorylation of ChiS are both dispensable for ChiS activation. One possible explanation for these results is that ChiS naturally lacks kinase activity. To test this, we purified the cytoplasmic domain of ChiS and found that ChiS was, in fact, capable of autophosphorylation *in vitro*, and that this activity was dependent on the conserved histidine in the HK domain (**Fig. S2**). This demonstrates that ChiS is capable of kinase activity; however, this activity is dispensable for P_*chb*_ induction.

In addition to autokinase and phosphotransfer activity, HKs also harbor phosphatase activity. In some HKs, a conserved threonine is critical for phosphatase activity, but is dispensable for kinase activity (10, 11). This residue is conserved in ChiS (T473A) (**Fig. S1**). We found that for ChiS^T473A^, there was a loss of P_*chb*_ induction when *cbp* is deleted (**Fig. 1a-b**). Complementation in the ChiS^T473A^ background with ChiS^WT^ restored activation of P_*chb*_-GFP (**Fig. S3**). These results suggest that ChiS phosphatase activity is critical for P_*chb*_ activation. To verify that ChiS^T473A^ retains its kinase activity as expected, we characterized this protein using *in vitro* assays. Indeed, the purified cytoplasmic domain of ChiS^T473A^ exhibited kinase activity *in vitro*, albeit at a reduced level compared to the parent (**Fig. S2**). This mutation may reduce kinase activity because this residue is in the same motif as the conserved histidine that is critical for kinase activity. Demonstrating ChiS phosphatase activity *in vitro* is challenging; because there are two sites of phosphorylation on the same protein, it is difficult to discern the phosphorylation status of each independent residue. Thus, we took a genetic approach to test whether ChiS^T473A^ was a poor activator of P_*chb*_ due to a loss of phosphatase activity.

If ChiS^T473A^ is phosphatase inactive, this would result in constitutive phosphorylation of the conserved aspartate in the ChiS Rec domain, which may inactivate ChiS and result in decreased P_*chb*_ expression. Therefore, we hypothesized that preventing phosphorylation of the ChiS Rec domain should recover P_*chb*_ activation, even in the ChiS phosphatase inactive background. To test this, we prevented phosphorylation of the ChiS Rec domain in the ChiS^T473A^ background by either mutating the conserved aspartate that is phosphorylated (ChiS^T473A D772A^) or by preventing ChiS kinase activity (ChiS^T473A H469A^). Both mutants rescued P_*chb*_ induction when *cbp* was deleted (**Fig. 1a-b**), suggesting that the ChiS^T473A^ mutation prevents activation of P_*chb*_ due to constitutive phosphorylation of the ChiS Rec domain. To further test whether phosphorylation of the Rec domain prevents activation of P_*chb*_, we generated a phosphomimetic allele by mutating the conserved aspartate to a glutamate (ChiS^D772E^) (12). We found that ChiS^D772E^ did not induce P_*chb*_ when *cbp* was deleted, further demonstrating that phosphorylation of the ChiS Rec domain prevents ChiS activity (**Fig. 1a**). Complementation in the ChiS^D772E^ background with ChiS^WT^ restored P_*chb*_ activation (**Fig. S3**).

In the above experiments, ChiS was activated genetically by deleting *cbp*. We next wanted to assess ChiS activity in a physiologically relevant context by using the natural inducer for this system, chitin oligosaccharides. To do so, we took two approaches: 1) we induced strains with chitin oligosaccharides and determined *chb* transcript abundance by qRT-PCR and 2) we assessed growth of strains on chitobiose. As expected, ChiS^WT^ induced P_*chb*_ in the presence of chitin oligosaccharides (**Fig. 1b**) and supported growth on chitobiose (**Fig. S4**). The Δ*chiS* strain neither activated *chb* expression when induced with chitin oligosaccharides nor grew on chitobiose, further verifying the role of ChiS as an essential activator of P_*chb*_ (**Fig. 1b, S4**). All ChiS point mutants where P_*chb*_ induction was observed in a Δ*cbp* background were also able to activate *chb* transcription when chitin oligosaccharides were used as an inducer and, accordingly, these strains grew like the parent (ChiS^WT^) on chitobiose (**Fig. 1b, S4**). ChiS^D772E^ supported a minor increase in P_*chb*_ expression, though not to the level of ChiS^WT^ (**Fig. 1b, S4**); consistent with this, ChiS^D772E^ grew poorly on chitobiose compared to the parent. All strains tested grew like the parent when tryptone or glucose was used as the carbon source (**Fig. S5**).

There was a discrepancy, however, when analyzing the ChiS^T473A^ phosphatase mutant. With ChiS^T473A^, we found that chitin oligosaccharides induced *chb* expression and this strain grew relatively well on chitobiose (**Fig. 1b & S4**). This result is contrary to the absence of P_*chb*_ induction observed for ChiS^T473A^ when *cbp* is deleted (**Fig. 1a-b**). Importantly, P_*chb*_ induction was restored when ChiS^T473A^ Δ*cbp* was complemented via ectopic expression of ChiS^WT^ (**Fig. S3**), suggesting that the lack of activation observed in ChiS^T473A^ Δ*cbp* was not simply due to a second site mutation. These data were the first to suggest that activation of ChiS is different when induced genetically via deletion of *cbp* vs naturally via chitin oligosaccharides. Also, because P_*chb*_ was not induced by chitin oligosaccharides in the ChiS^T473A^ Δ*cbp* background, this suggested that the presence of both CBP and chitin were required for activation (**Fig. 1b**). To explore the underlying mechanism, we next investigated the role of CBP in regulating ChiS activity.

### CBP directly interacts with ChiS in the presence and absence of chitin to regulate ChiS activity

It is hypothesized that CBP directly interacts with the ChiS periplasmic domain to repress ChiS activity. Upon CBP binding to chitin oligosaccharides, it is believed that this repression is relieved, allowing ChiS to activate the chitin utilization program (4). To test this model, we first sought to determine whether ChiS and CBP directly interact with one another. To do so, we took advantage of a recently developed cytological assay called the PopZ-Linked Apical Recruitment (POLAR) assay, which is well-suited to study interactions between cell envelope proteins (13). POLAR interrogates protein-protein interactions based on co-localization of fluorescently-tagged “bait” and “prey” proteins. Evidence for direct interaction between proteins is provided by re-localizing the bait protein to the cell poles and by assessing whether the localization pattern of the prey is similarly altered. Re-localization to the cell poles is accomplished by tagging the bait with a PopZ interaction domain (called an H3H4 domain) and through ectopic expression of PopZ, which naturally localizes to the cell pole (14, 15).

For POLAR, we generated a ChiS allele with a GFP-H3H4 tag at its C-terminus (ChiS-GFP-H3H4). This protein was non-functional for P_*chb*_ induction; however, the predicted periplasmic domain of this allele is intact (**Fig. S6a**). We also generated a CBP-mCherry fusion at the native locus. CBP-mCherry was fully functional for repression of ChiS activity in the absence of chitin (**Fig. S6b**). It also supported P_*chb*_ induction in the presence of chitin oligosaccharides and growth on chitobiose, albeit at slightly reduced levels compared to the untagged CBP^WT^ parent (**Fig. S6c-e**). In a strain containing both of these fluorescent fusions, we observed that ChiS-GFP-H3H4 and CBP-mCherry co-localized as puncta at the cell periphery in the absence of PopZ (**Fig. 1c-d, top row**). Upon ectopic expression of PopZ, ChiS-GFP-H3H4 was relocalized to the cell poles, and we observed a concomitant relocalization of CBP-mCherry to the cell poles (**Fig. 1c-d, middle row**). By contrast, in a strain lacking ectopic ChiS-GFP-H3H4, CBP-mCherry was diffusely localized in the periplasm whether PopZ was expressed or not (**Fig. S7**). Together, these data strongly suggest that ChiS and CBP directly interact.

Next, we wanted to investigate what happens to the CBP-ChiS complex in the presence of chitin oligosaccharides. If CBP is purely inhibitory to ChiS, we hypothesized that the presence of chitin oligosaccharides would result in CBP dissociation from ChiS, and CBP would become diffusely localized in the periplasm. However, when cells were incubated with chitin oligosaccharides, we found that CBP-ChiS interactions were not disrupted (**Fig. 1c-d, bottom row**), even after prolonged incubation (**Fig. S8**). These data suggest that CBP may not dissociate from ChiS in the presence of chitin, but instead remains bound.

Above, we show that ChiS^T473A^ can be activated by chitin oligosaccharides and that this activation requires the presence of CBP (**Fig. 1b, S4**). By contrast, ChiS^T473A^ cannot be activated by deletion of *cbp* (**Fig. 1a-b**). Additionally, the POLAR assay suggests that CBP may not dissociate from ChiS in the presence of chitin oligosaccharides (**Fig. 1c-d**). Together, we believe these data suggest that CBP regulates ChiS activity in two ways. First, CBP binding to ChiS in the absence of chitin represses ChiS activity; possibly by altering the conformation of ChiS to an inactive state and by favoring phosphorylation of the ChiS Rec domain. In support of the latter point, mutations to ChiS that prevent or poorly phosphorylate the highly conserved histidine (ChiS^H469A^, ChiS^H469A D772A^, ChiS^T473A D772A^, and ChiS^T473A H469A^) induce P*chb* expression to a small degree, even without induction by chitin or deletion of *cbp* (**Fig. 1a-b**). Second, we propose that chitin-bound CBP represses ChiS kinase activity; this favors dephosphorylation of the ChiS Rec domain, thereby activating ChiS. The suppression of ChiS kinase activity is not essential when ChiS^WT^ is induced by deletion of *cbp* because ChiS^WT^ contains sufficient phosphatase activity to overcome its kinase activity when CBP is absent. However, repression of ChiS kinase activity by chitin-bound CBP is essential in the ChiS^T473A^ background because this allele lacks phosphatase activity. This model helps reconcile the absence of P_*chb*_ induction observed for ChiS^T473A^ via deletion of *cbp*, but the robust induction observed when chitin oligosaccharides are used (**Fig. 1a-b**).

Moving forward, we were interested in studying the mechanism underlying ChiS-dependent activation of P_*chb*_. Chitin oligosaccharides are prohibitively expensive for studying ChiS activity. We show above, however, that deletion of *cbp* serves as a reliable genetic method to activate ChiS (other than in ChiS^T473A^). Thus, deletion of *cbp* was used to induce ChiS activity throughout the remainder of the study.

### ChiS is a one-component membrane-bound phosphorylation-dependent DNA binding protein

ChiS is a hybrid HK and is presumed to regulate gene expression in conjunction with a cognate RR tentatively named ChiR (5). However, no candidate for ChiR has ever been identified despite numerous attempts from our group and others (4, 5, 7-9, 16). Additionally, we have observed that ChiS is active when its kinase activity is ablated, making it unlikely that this HK passes a phosphate to a cognate RR. This led us to hypothesize that ChiS directly regulates gene expression in response to chitin.

To test this, we assessed whether ChiS was sufficient to activate a P_*chb*_-*lacZ* reporter in the heterologous host *Escherichia coli*, which lacks a ChiS homolog. Similar to what we observe in *V. cholerae*, ChiS induced P_*chb*_ expression in *E. coli* and this activity was diminished in the phosphomimetic ChiS^D772E^ background (**Fig. 2a**). Together, these data suggest that ChiS directly activates P_*chb*_.

**Figure 2.**
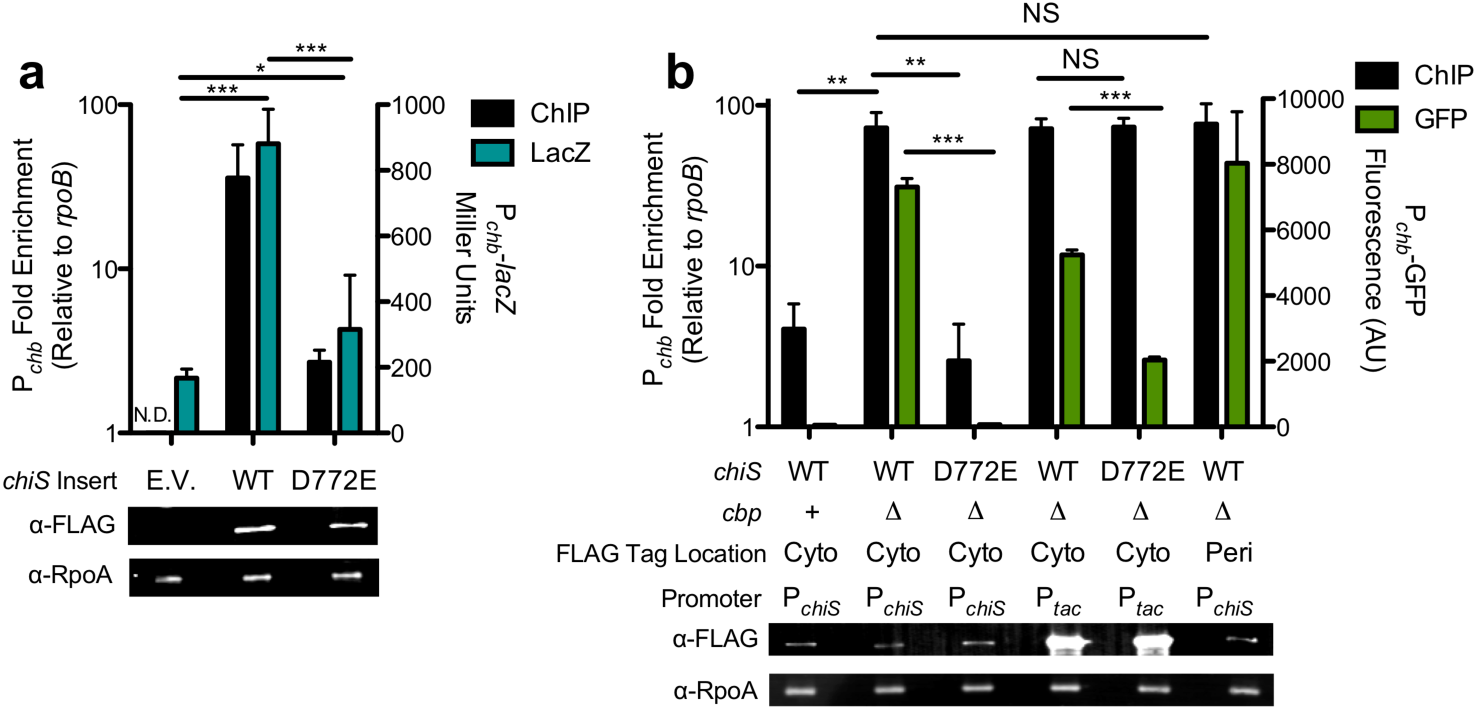
ChiS is a one-component membrane-bound phosphorylation-dependent DNA binding protein. (**a**) Miller assay and chromatin immunoprecipitation (ChIP) performed in *E. coli* MG1655 containing a chromosomally-encoded P_*chb*_-*lacZ* reporter and pMMB empty vector (E.V.) or vectors containing the ChiS allele indicated. Untagged ChiS alleles were used for Miller assays (blue bars), while ChiS-FLAG alleles were used for ChIP assays (black bars). P_*chb*_ enrichment was assessed relative to *rpoB* for ChIP assays. For the empty vector strain, ChIP enrichment was not determined (N.D.). (**b**) *V. cholerae* strains expressing ChiS-FLAG alleles under native or P_*tac*_ overexpression conditions as indicated were assessed for P_*chb*_ enrichment relative to *rpoB* via ChIP (black bars), activation of a P_*chb*_-GFP reporter (green bars), and expression of either ChiS (anti-FLAG) or RpoA (anti-RpoA; loading control) via Western blot analysis. ChiS alleles had a FLAG tag localized to either the cytoplasmic domain (cyto) or the periplasmic (peri) domain of ChiS; for ChiS membrane topology, see **Fig S10a**. All data are from at least three independent biological replicates and shown as the mean ± SD. Statistical comparisons were made by one-way ANOVA with Tukey’s post-test. NS, not significant. *, p < 0.05. **, p < 0.01. ***, p < 0.001.

We hypothesized that ChiS directly regulates P_*chb*_ by binding to the promoter. To test this, we generated a functional internally FLAG-tagged ChiS (**Fig. S9**) and performed ChIP-qPCR to see if this protein bound to the *chb* promoter *in vivo*. When ChiS is active (ChiS-FLAG^WT^ Δ*cbp*), P_*chb*_ was enriched ∼70-fold, suggesting that ChiS directly binds P_*chb*_ *in vivo* (**Fig. 2b**). Enrichment of P_*chb*_ was also observed in the heterologous host *E. coli* (**Fig. 2a**). When ChiS activity is repressed due to the presence of CBP in *V. cholerae*, we find that ChiS binding to P_*chb*_ is significantly reduced, suggesting that CBP may antagonize ChiS by preventing its DNA binding activity (**Fig. 2b**).

The DNA-binding activity of prototypical RRs is controlled by the phosphorylation status of the conserved aspartate in their Rec domain. To determine whether the phosphorylation status of ChiS Rec plays a role in regulating DNA-binding, ChIP assays were performed with the phosphomimetic ChiS-FLAG^D772E^. We observed that ChIP enrichment and P_*chb*_ expression were significantly decreased (**Fig. 2b**), suggesting that phosphorylation of the Rec domain decreases the affinity of ChiS for DNA, which results in loss of *chb* expression.

The phosphorylation state of RRs, however, generally only alters their affinity for DNA and not their absolute ability to bind DNA. Consistent with this, overexpression of ChiS-FLAG^D772E^ resulted in ChIP enrichment similar to the ChiS parent (**Fig. 2b**). Additionally, overexpression of ChiS-FLAG^D772E^ partially restored P_*chb*_ expression (**Fig. 2b**), suggesting that reduced activation by this phosphomimetic allele is largely attributed to its reduced affinity for DNA.

ChiS has two predicted transmembrane (TM) domains. Based on the domain architecture, we predicted that the sequence between the two TM domains would be periplasmic, and the sequence following the second TM would be cytoplasmic (predicted topology in **Fig. S10a**). We tested the membrane topology of ChiS using *lacZ* and *phoA* fusions, which rely on the observation that LacZ is only functional in the cytoplasm and PhoA is only functional in the periplasm (17-19). The LacZ fusions tested exhibited high activity only when linked to a residue before the first TM (ChiS^1-2^), or past the second TM (ChiS^1-452^ and ChiS^1-1129^; **Fig. S10b**). Conversely, the only PhoA fusion that was functional was linked to a residue between the first and second TMs (ChiS^1-52^; **Fig. S10b**). These data were consistent with the predicted membrane topology (**Fig. S10a**).

Typically, DNA-binding transcription factors are soluble cytoplasmic proteins. It is possible that ChiS is proteolytically processed to release a cytoplasmic DNA-binding portion of the protein, or it is possible that ChiS directly binds to DNA from the membrane. If ChiS is post-translationally processed, we hypothesized that P_*chb*_ should only be enriched during ChIP experiments when the FLAG tag in ChiS is cytoplasmically-localized, and not when the FLAG tag is periplasmically-localized because the latter should be separated from the cytoplasmic DNA-binding domain following proteolytic cleavage. To test this, we generated two functional alleles of ChiS where the FLAG tag was inserted in the cytoplasmic domain (after residue E566; see **Fig. S10a**) or the periplasmic domain (after residue T287; see **Fig. S10a**). We observed similar levels of P_*chb*_ enrichment in both strains (**Fig. 2b**), indicating that ChiS likely binds P_*chb*_ from the membrane *in vivo*.

### Specific binding of ChiS to P_chb_ is required for activation of the chb operon

Thus far, we have shown that ChiS binds to P_*chb*_ *in vivo* via ChIP assays. This assay relies on crosslinking to stabilize protein-DNA interactions. So, it remains possible that the enrichment we observed was not due to ChiS directly binding P_*chb*_, but instead was the result of ChiS being in a complex with a crosslinked partner protein (like an RR) that binds to DNA. To further test whether ChiS binds DNA directly or indirectly, we purified a fragment of ChiS encompassing the Rec domain and C-terminus (ChiS^Rec-C^) and tested its ability to bind P_*chb*_ via electrophoretic mobility shift assays (EMSAs) *in vitro*. We found that purified ChiS^Rec-C^ binds to a P_*chb*_ DNA probe. This interaction could be competed with unlabeled P_*chb*_, but not an unrelated promoter, demonstrating that this interaction is both direct and specific (**Fig. S11**). Next, we sought to narrow down the location of the ChiS binding site(s) within the *chb* promoter. Through EMSA analysis of P_*chb*_ promoter fragments (**Fig. S12**), we identified two distinct 13bp sequences that represented two putative ChiS binding sites (CBSs).

To test these putative CBSs, a 60bp probe that spans both sites (**Fig. S12**) was used for EMSA analysis (**Fig. 3a**). When ChiS was incubated with the WT probe, we observed three shifts. Two of these shifts are likely due to the two CBSs in this probe, while the third shift may result from DNA bending as described for other transcriptional regulators (20, 21). Mutation of either CBS in isolation resulted in loss of one shift, while mutation of both CBSs completely prevented ChiS binding (**Fig. 3a**). Additionally, ChiS did not bind to a full length P_*chb*_ probe when both CBSs were mutated, confirming that these are the only two CBSs within this promoter (**Fig. S11**). Furthermore, use of the CBS 1 & 2 mutated P_*chb*_ probe as a cold competitor did not prevent ChiS binding to WT P_*chb*_ (**Fig. S11**). Finally, the CBSs are conserved within P_*chb*_ in diverse *Vibrio* species, further suggesting that ChiS binds specific sequences in P_*chb*_ (**Fig. S13a**).

**Figure 3.**
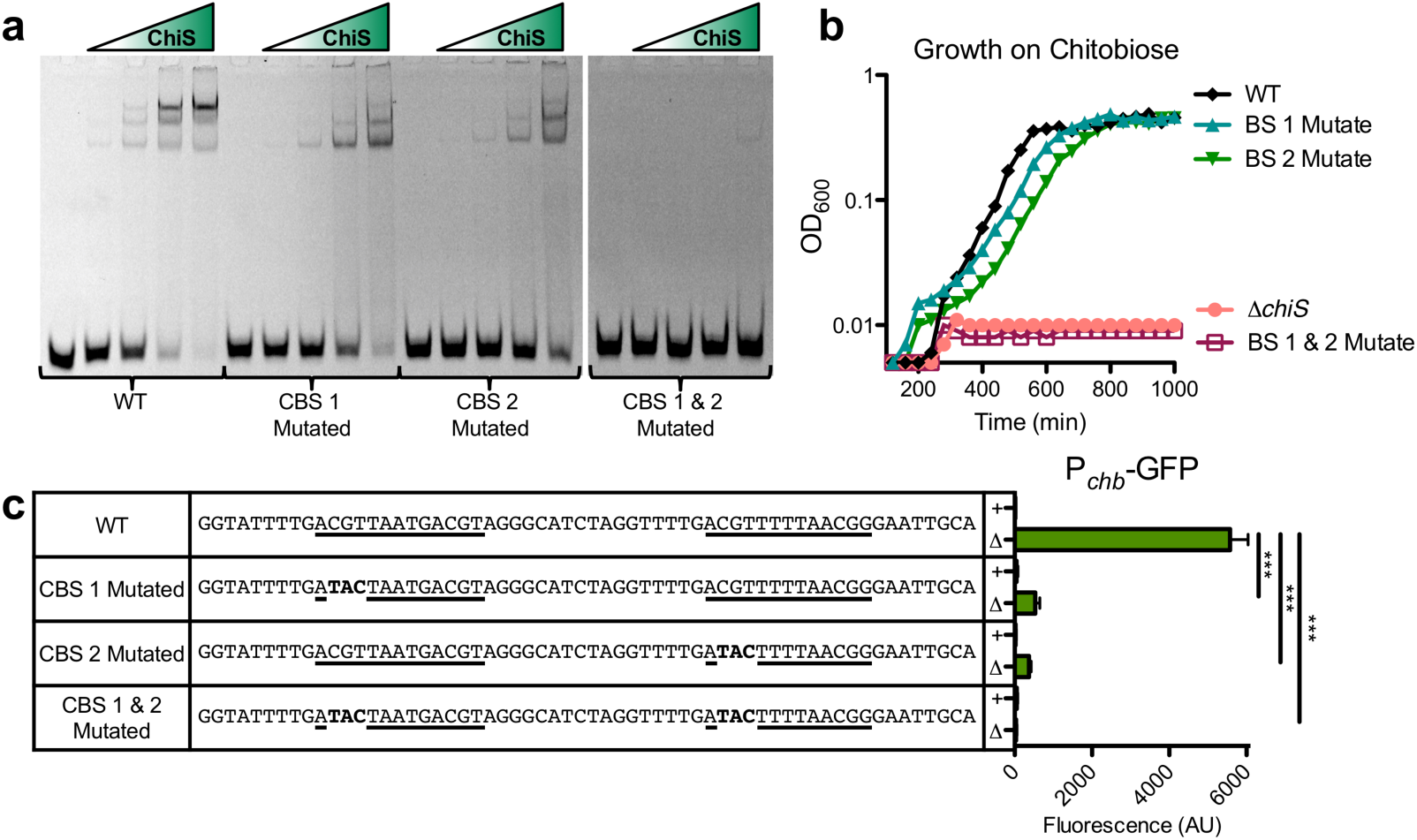
Specific binding of ChiS to P_chb_ is required for activation of the chb operon. (**a**) Electrophoretic mobility shift assays (EMSAs) were carried out using a purified portion of the ChiS cytoplasmic domain (ChiS^Rec-C^, residues 725-1129 – see **Fig. S10a**) and a Cy5-labeled 60 bp fragment of the P_*chb*_ promoter that is either intact or mutated as indicated (exact sequences in **c**). This 60 bp fragment encompasses both ChiS binding sites (CBSs) (see **Fig. S12**). (**b**) Strains containing the indicated mutations in the native P_*chb*_ promoter were assessed for growth on chitobiose. Data in **a-b** are representative of at least 2 independent experiments. (**c**) Expression of a P_*chb*_-GFP reporter containing the indicated mutations to the ChiS binding sites was assessed in strains where *cbp* is intact (+) or deleted (Δ) as indicated. Data are the result of at least three independent biological replicates and shown as the mean ± SD. Statistical comparisons were made by one-way ANOVA with Tukey’s post-test. ***, p < 0.001.

To characterize the role of the CBSs in P_*chb*_ activation and *V. cholerae* physiology, we mutated these sites in both the P_*chb*_-GFP reporter and at the native locus. Mutation of either CBS significantly decreased P_*chb*_ expression (**Fig. 3c**) and correspondingly slightly delayed growth on chitobiose (**Fig. 3b**). However, mutation of both CBSs completely prevents P_*chb*_ activation and growth on chitobiose (**Fig. 3b-c**). Collectively, these data indicate that ChiS binding to the *chb* promoter is required for activation.

Our *in vivo* and *in vitro* data showing that ChiS directly binds P_*chb*_ was unexpected because ChiS lacks a canonical DNA binding domain based on primary sequence (BLAST (22)) or structural (Phyre2 (23)) homology predictions. We hypothesized that the C-terminal 197 amino acids of ChiS (residues 937-1129; see **Fig. S10**), which have no predicted homology to other domains and are conserved among ChiS homologs (**Fig. S13b**), encoded a noncanonical DNA binding domain. To test this, we performed ChIP experiments with N-terminal truncations of ChiS, which revealed that the C-terminal 106 amino acids of ChiS were sufficient to bind P_*chb*_ *in vivo* (**Fig. S14a**). Furthermore, deletion of these residues from full length ChiS prevented P_*chb*_ enrichment (**Fig. S14a**), indicating that this domain is necessary and sufficient for DNA binding. These experiments also revealed that while cytoplasmic fragments of ChiS were sufficient to bind P_*chb*_ *in vivo*, they did not activate P_*chb*_ expression (**Fig. S14b)** and poorly facilitated growth on chitobiose (**Fig. S14d**).

### ChiS may activate P_chb_ by direct recruitment of the alpha subunit of RNAP

Finally, we sought to define the mechanism by which ChiS activates gene expression. Some activators promote transcription by directly recruiting RNA Polymerase (RNAP). Previous work with activators like CRP, shows that the absolute distance between the activator binding site and promoter is not critical for activation, but the phasing of DNA between these sites is critical to ensure that the transcription factor recruits RNAP in the correct orientation (24, 25). One turn of B DNA is 10bp; thus, insertions or deletions of 10bp should maintain helical phasing, while insertions or deletions of 5bp should ablate phase-dependent interactions.

To assess if CBSs needed to be in phase with the *chb* promoter to mediate activation of P_*chb*_, we inserted 5 or 10 bp between the CBSs and the -35 signal (**Fig. 4a, Region A**). Insertion of 5bp, which would put the CBSs and -35 out of phase, resulted in loss of P_*chb*_ expression; insertion of 10bp, which maintains helical phasing, allowed for partial activation (**Fig. 4a-b**). A distance-dependent reduction in transcriptional activation despite maintenance of phase-dependent interactions is consistent with prior results in other regulators (25). These results suggest that CBSs must be in phase with the -35 to activate P_*chb*_.

**Figure 4.**
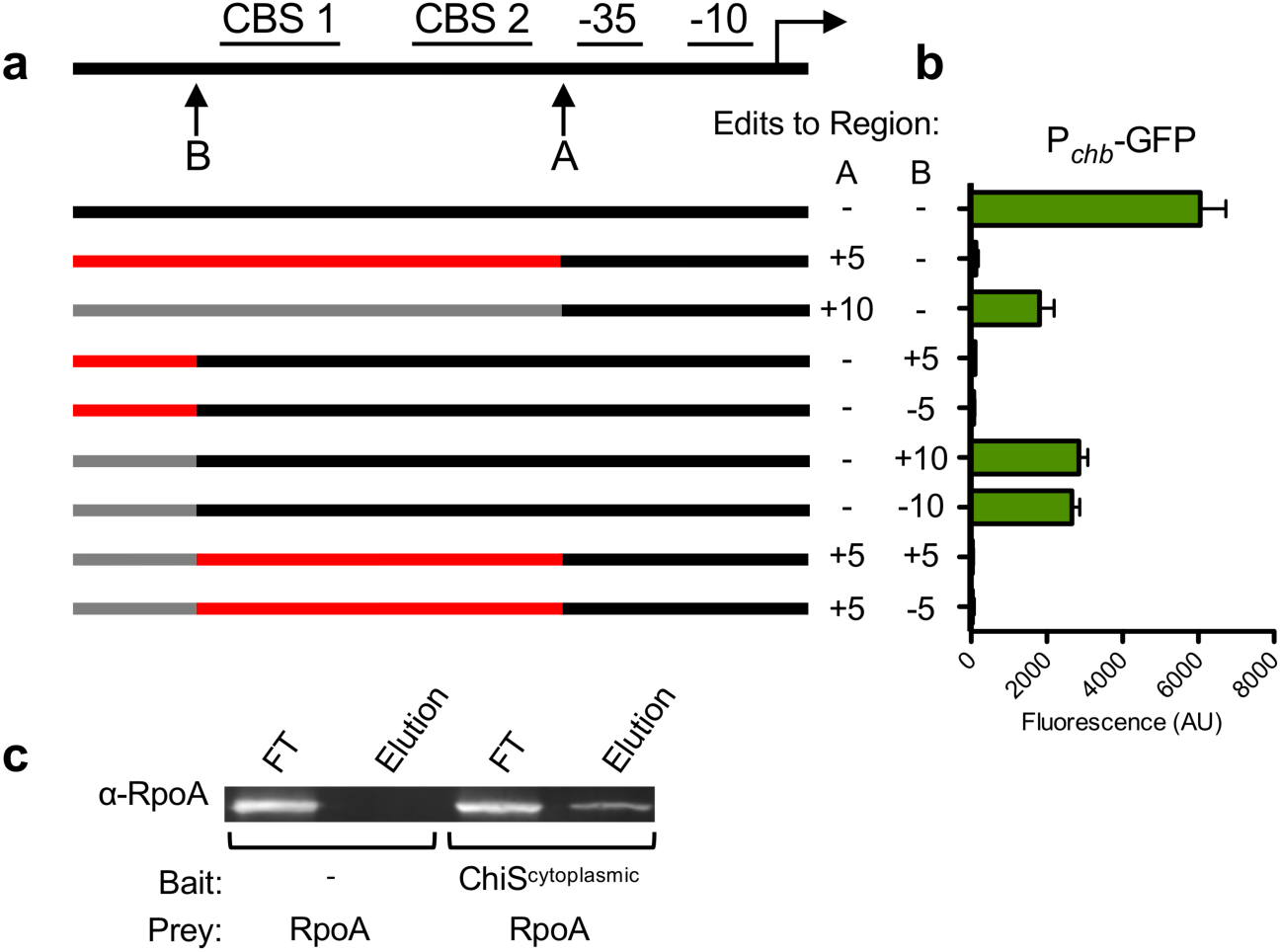
ChiS may activate P_chb_ by direct recruitment of the alpha subunit of RNAP. (**a**) Schematic of the approach to test phase-dependent activation of P_*chb*_. Indels of 5 or 10bp were introduced at region A and B to alter the phasing of the elements within P_*chb*_ relative to the -35 signal. For each mutant generated, the schematic highlights the phasing of promoter elements with respect to the -35. WT phasing is indicated in black, if the sequence is mutated to be out-of-phase it is indicated in red, and if the sequence is mutated to maintain helical phasing it is indicated in gray. (**b**) P_*chb*_-GFP expression was determined for strains harboring the indicated phase-mutated P_*chb*_-GFP alleles. Data are the result of at least three independent biological replicates and shown as the mean ± SD. (**c**) Protein pulldown assays were performed using purified ChiS (MBP-tagged ChiS^cytoplasmic^) as bait and purified RpoA as prey as indicated. The presence of the RpoA prey was assessed in the flow through (FT) and elution by Western blot analysis. Data are representative of two independent experiments.

In addition to ChiS, other DNA binding proteins, like SlmA, bind upstream of the CBSs and contribute to P_*chb*_ activation (7). To assess if upstream factors exhibit helical phase-dependence for P_*chb*_ activation, we inserted or deleted 5 or 10 bp upstream of the CBSs (**Fig. 4a, Region B**). We found that indels of 5bp (loss of helical phasing) resulted in loss of P_*chb*_ expression, while indels of 10bp (helical phasing maintained) allowed for partial activation (**Fig. 4a-b**). This suggests that factors upstream of the CBSs must be in phase for proper activation of P_*chb*_.

Collectively, these data support either of the following models for proper P_*chb*_ activation: 1) CBSs and other upstream activator binding sites must all be in phase with the -35, or 2) only upstream activator binding sites need to be in phase with the -35 while the CBSs do not. To distinguish between these two models, we introduced mutations to isolate the phasing of the CBSs. When 5bp is inserted at Region A, both the CBSs and the upstream activator binding sites are out of phase. We restored helical phasing for the upstream activator binding sites, but not the CBSs, by inserting or deleting 5bp from Region B. We observed that P_*chb*_ activation is not restored when phasing is restored for the upstream activator binding sites, which suggests that both the CBSs and upstream activator sites must be in phase with the -35 signal for activation of P_*chb*_. The spacing between the CBSs and the -35 signal in P_*chb*_ is conserved across *Vibrio* species (**Fig. S13a**), further indicating an important role for DNA phasing in activation of *chb* expression.

Helical phase dependence between the CBSs and the -35 supports a model where ChiS directly recruits RNAP. Transcriptional activators that recruit RNAP often interact with the α-subunit, and so we hypothesized that ChiS may directly interact with RpoA. To test this, we assessed direct binding between purified ChiS^cytoplasmic^ and RpoA *in vitro* using a protein pulldown assay. In pulldown reactions where RpoA was the prey, RpoA only came out in the elution when ChiS^cytoplasmic^ was used as the bait, suggesting that ChiS and α-subunit interact (**Fig. 4**). Reciprocal pulldowns using RpoA as the bait further confirmed this interaction (**Fig. S15**). Despite interacting with RpoA *in vitro*, ChiS^cytoplasmic^ does not activate gene expression *in vivo* (**Fig. S9b-c**). One possibility for this observation is that the interaction between full length ChiS and RpoA *in vivo* is much stronger than the interaction observed *in vitro* with ChiS^cytoplasmic^. This could be because the cytoplasmic fragment of ChiS is not as rigidly locked in the active form compared to full length membrane-bound ChiS. Alternatively, it is possible that the interaction between ChiS^cytoplasmic^ and RpoA *in vivo* are in the wrong conformation to support transcriptional activation while full-length ChiS may bind RpoA in the correct conformation to allow for transcriptional activation. Regardless, these results suggest that ChiS can directly bind to RNAP.

## Discussion

TCSs represent a diverse family of proteins that allow bacteria to sense and respond to their environment. Here, we describe a protein that, based on homology, falls into the two-component family, but uses a mechanism that is contrary to the canon for these systems. ChiS is noncanonical in two ways (**Fig. 5**). First, phosphorylation of the Rec domain diminishes its DNA-binding activity while dephosphorylation enhances it. To our knowledge, this is the first example of such regulation among TCS family proteins. Second, ChiS is able to bind P_*chb*_ from the membrane to regulate gene expression. This result was particularly surprising because ChiS lacks a canonical DNA-binding domain.

**Figure 5.**
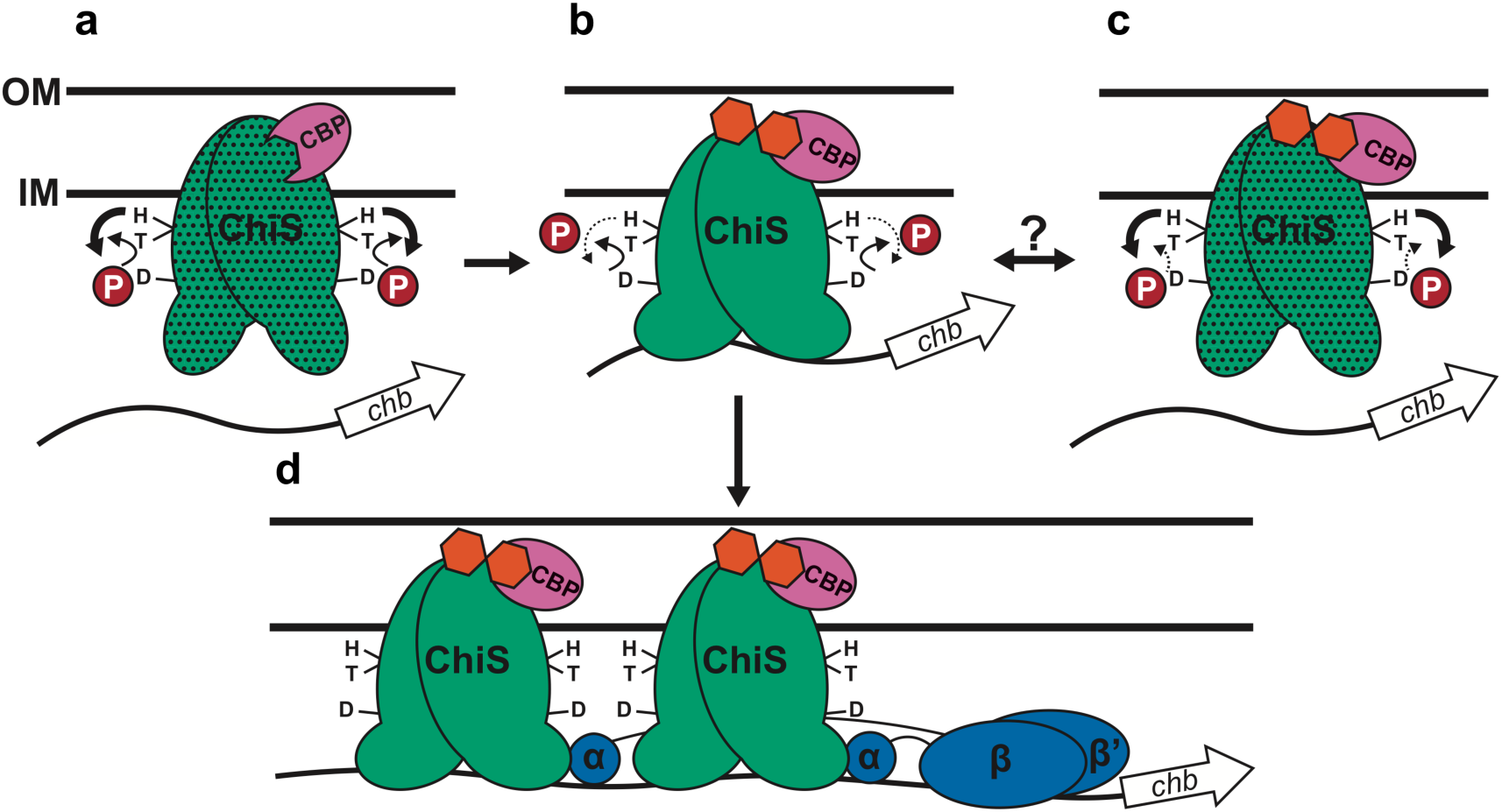
Proposed model for ChiS-mediated activation of P_chb_. (**a**) In the absence of chitin, CBP (pink) interacts with the ChiS (green) periplasmic domain, which prevents activation of P_*chb*_ by (1) promoting ChiS kinase activity and (2) by keeping ChiS in a conformationally inactive state (stippled). (**b**) In the presence of chitin (orange hexagons), chitin-bound CBP remains bound to ChiS and likely (1) diminishes ChiS kinase activity and (2) places ChiS into a conformationally active state. This ultimately allows the ChiS C-terminus to bind to the *chb* promoter. (**c**) Additional currently unknown regulatory cues may affect ChiS activity by either enhancing its kinase activity or diminishing its phosphatase activity. (**d**) Dephosphorylated ChiS (as in **b**) can bind P_*chb*_ and promote transcription by recruiting RNA polymerase through a direct interaction with the alpha subunit.

It is tempting to speculate that the ChiS C-terminus represents a novel class of DNA-binding domain. Homology searches (BLAST) revealed that this DNA-binding domain is exclusively found at the C-terminus of other proteins (**Fig. S16**), which often contain an N-terminal sensing domain (i.e. HK/Rec or PAS (6, 26, 27)). This analysis also revealed that this domain is largely restricted to gamma- and deltaproteobacteria, with a few examples among alphaproteobacteria. This suggests that this DNA-binding domain may not be unique to *V. cholerae* ChiS, but may be a conserved output domain for diverse signal transduction proteins in proteobacteria.

Here, we have also clarified the mechanism of CBP regulation of ChiS. First, CBP regulates ChiS differentially depending on the presence or absence of chitin in the periplasm. In the absence of chitin, CBP inhibits ChiS by preventing its DNA binding activity (**Fig. 5a**). The mechanism of inhibition by CBP is likely due to a combination of 1) regulating ChiS enzymatic function (i.e. kinase/phosphatase activity) and 2) inducing an inactive conformational change in ChiS. In support of the first point, we see that ChiS activity is partially derepressed in mutants where the conserved His and/or Asp sites are mutated even when *cbp* is intact (**Fig. 1a-b**). Because CBP is still able to repress ChiS activity in the unphosphorylated backgrounds (ChiS^H469A^, ChiS^D772A^, ChiS^H469A D772A^, and ChiS^T473A H469A^; **Fig. 1a-b**), CBP also represses ChiS independent of regulating its kinase/phosphatase activity, possibly by inducing an inactive conformation in ChiS. Thus, deletion of *cbp* can relieve the repressive conformational change and serves as one mechanism to activate ChiS. We show that in the presence of chitin, CBP maintains its interaction with ChiS (**Fig. 5b-d**). Furthermore, our data suggest that chitin-bound CBP likely inhibits ChiS kinase activity (**Fig. 5b**) because chitin oligosaccharides and CBP are both required for P_*chb*_ induction in the ChiS^T473A^ phosphatase mutant background (**Fig. 1a-b**). The regulation of histidine kinases by ligand-binding periplasmic proteins is a well-known phenomenon. For example, the AI-2 autoinducer sensing system LuxPQ is made up of a periplasmic AI-2 binding protein (LuxP) and a membrane embedded hybrid histidine kinase (LuxQ). In the absence of *luxP* or at low AI-2 concentrations, LuxQ acts as a kinase (28). Upon AI-2-bound LuxP binding to the periplasmic domain of LuxQ, the enzymatic activity of LuxQ shifts to phosphatase activity.

ChiS activity is regulated by the physiology of the cell at multiple levels. It is regulated by the presence of chitin oligosaccharides as described above. ChiS activity is also negatively regulated by carbon catabolite repression (CCR) (9), suggesting that the chitin utilization program is suppressed in the presence of preferred carbon sources. The mechanism underlying CCR-dependent inhibition of ChiS, however, remains unclear and will be the focus of future work. It is also possible that additional cues regulate ChiS activity. For example, it remains unclear what role ChiS kinase/phosphatase activity plays in regulating this system because CBP-dependent repression ensures that ChiS activity is still largely suppressed in the absence of chitin. It is possible that additional currently unknown factors regulate ChiS kinase/phosphatase activity (**Fig. 5c**). ChiS has a C-terminal PAS domain (**Fig. S10a**) which may modulate ChiS enzymatic activities through small ligand binding or mediating protein-protein interactions as described for other HKs (29, 30). The complex regulation of ChiS activity may serve to integrate additional cues that ultimately control expression of the chitin utilization program.

Hybrid HKs that putatively bind DNA are uncommon, but not unprecedented. *Bacteroides thetaiotaomicron* encodes 33 hybrid HKs, 32 of which contain AraC-type DNA binding domains within their C-terminus (31). However, these proteins have merely been identified by bioinformatic analyses and, to our knowledge, the molecular mechanism of DNA-binding HKs has not previously been tested in any system. Here, we show that ChiS is a one-component hybrid HK that directly binds DNA from the membrane to regulate gene expression. Aside from TCSs, membrane-embedded DNA-binding transcriptional regulators are conserved in bacterial species, with ChiS representing the fifth such member within *V. cholerae* (alongside ToxR, TcpP, CadC, and TfoS (8, 16, 32-35)). We show that ectopic expression of just the cytoplasmic domain of ChiS is sufficient to bind DNA *in vivo*, but does not activate gene expression (**Fig. S14**). Thus, it is tempting to speculate that the capacity of membrane-embedded regulators to recruit target promoters to the cell periphery is critical for their ability to activate transcription (36). Also, the mechanism by which membrane-embedded regulators access their target sites in the genome while being sequestered in the membrane remains poorly understood (37). ChiS may serve as a valuable model system to dissect the molecular mechanism of membrane-embedded regulators, which will be the focus of future work.

## Methods

### Bacterial strains and culture conditions

All *V. cholerae* strains used in this study are derived from the El Tor strain E7946 (38). The *E. coli* strains used to test ChiS activity in a heterologous host were derived from MG1655 (39), while the strains used to test ChiS membrane topology were derived from *E. coli* DH1 (40). *V. cholerae* and *E. coli* strains were grown in LB medium and on LB agar supplemented when necessary with carbenicillin (20 or 100 μg/mL), kanamycin (50 μg/mL), spectinomycin (200 μg/mL), trimethoprim (10 μg/mL), and/or chloramphenicol (2 μg/mL).

### Generating mutant strains

*V. cholerae* mutant constructs were generated using splicing-by-overlap extension exactly as previously described (41). See **Table S3** for all of the primers used to generate mutant constructs in this study. Mutant *V. cholerae* strains were generated by chitin-dependent natural transformation and cotransformation exactly as previously described (42, 43). The P_*chb*_-*lacZ* reporter was introduced into *E. coli* MG1655 using λ Red recombineering (44). *E. coli* MG1655 harboring pKD46 was subcultured in SOB medium supplemented with 100 μg/mL carbenicillin and 0.02% arabinose at 30°C until an OD_600_ of 0.6 was reached. Cells were washed with ice cold 10% glycerol, electroporated with 200 ng Kan^R^-P_*chb*_ PCR product (amplified using primers described in **Table S3**), outgrown in SOC medium for 1 hour at 37°C, then plated on selective media. Plasmids were mated into *E. coli* and *V. cholerae* strains using *E. coli* S17 (45). Mutant strains were confirmed by PCR and/or sequencing. See **Table S2** for a detailed list of mutant strains used in this study.

### Plasmid construction

Plasmids for protein purification and the Miller assays to test ChiS activity in **Fig. 2** were constructed using the FastCloning method (46). Vectors and inserts were amplified using the primers listed in **Table S3**. Plasmids used for the Miller assays to define ChiS membrane topology in **Fig. S10** were made by cloning amplified inserts into the SalI and XbaI sites of pMW-SY*lacZ* or pMW-SY*phoA*.

### Measuring GFP and mCherry reporter fluorescence

GFP and mCherry fluorescence was determined essentially as previously described (7, 47). Briefly, single colonies were picked and grown in LB broth at 30°C for 18 hours. Cells were then washed and resuspended to an OD_600_ of 1.0 in instant ocean medium (7 g/L; Aquarium Systems). Then, fluorescence was determined using a BioTek H1M plate reader with excitation set to 500 nm and emission set to 540 nm for GFP and excitation set to 580 nm and emission set to 610 nm for mCherry.

### Quantitative reverse transcriptase PCR (qRT PCR)

From overnight cultures, strains were subcultured into LB medium and grown to an OD_600_ of ∼2.5. Cells were then induced with sterile water (as the no inducer condition) or chitin hexasaccharide (as the +(GlcNAc)_6_ condition; commercially available from Carbosynth) to a final concentration of 0.05% and grown for an additional 1 hour. RNA was purified, reverse transcribed, and then detected via qPCR exactly as previously described (7). The primers used for qRT-PCR experiments are found in **Table S3**.

### Microscopy data collection and analysis

Cultures were grown overnight in LB medium supplemented with inducers (10 μM IPTG and/or 0.05% arabinose) where indicated. Cells were then washed in instant ocean medium, resuspended to an OD_600_ of 0.25, and placed on a coverslip under an 0.2% gelzan pad made in instant ocean medium. Cells were imaged on an inverted Nikon Ti-2 microscope with a Plan Apo 60x objective lens, FITC and mCherry filter cubes, a Hamamatsu ORCAFlash 4.0 camera, and Nikon NIS Elements imaging software. Heat maps of ChiS-GFP-H3H4 and CBP-mCherry were generated on Fiji (48) using the MicrobeJ plugin (49).

### Growth curves

Growth was kinetically monitored at 30°C with shaking using a BioTek H1M plate reader with absorbance set to 600 nm. Growth was tested using M9 minimal medium supplemented with the indicated carbon source to a final concentration of 0.2%. For the growth curves carried out in **Fig. S14**, the inocula were grown overnight in 10 μM IPTG and growth reactions were also supplemented with 10 μM IPTG to induce ectopic expression of the ChiS alleles indicated. *V. cholerae* can grow on chitobiose through both the activity of the chitobiose ABC transporter encoded within the *chb* operon and through a GlcNAc PTS transporter (7, 50). ChiS is only required for regulation of the chitobiose ABC transporter (7). Thus, to study the effect of ChiS on regulating chitobiose utilization, the PTS GlcNAc transporter VC0995 was inactivated in all strains used to test growth on chitobiose as previously described (7).

### Miller assay

To assess ChiS activity in MG1655 strains, overnight cultures were inoculated from single colonies and grown at 30°C for 18 hours in LB broth supplemented with carbenicillin (100 µg/mL), kanamycin (50 µg/mL), and IPTG (10 µM). LacZ activity was then determined exactly as previously described (51).

For membrane topology experiments, overnight cultures of strains grown at 37°C in LB broth supplemented with ampicillin (100 µg/mL) were subcultured 1:50 into fresh LB broth containing ampicillin (100 µg/mL) and grown with shaking at 37°C for 2.5 hours to an OD_600_ of ∼0.7-1.0. Cells were then harvested, and alkaline phosphatase (PhoA) and beta-galactosidase (LacZ) activities of strains (as appropriate) were determined as previously described (52, 53).

### Chromatin immunoprecipitation (ChIP) assay

Overnight cultures of strains for ChIP were diluted to an OD_600_ of 0.08 and then grown for 6 hours at 30°C with rolling. For strains with natively expressed ChiS, cells were grown in plain LB broth. For *V. cholerae* strains containing P_*tac*_ constructs, cells were grown in LB broth supplemented with IPTG (10 μM). For *E. coli* strains, cells were grown in LB broth supplemented with kanamycin (50 µg/mL), carbenicillin (100 µg/mL), and IPTG (10 µM). Cultures were crosslinked with paraformaldehyde (1% final concentration) for 20 minutes at room temperature, quenched with Tris for 10 minutes at room temperature, washed twice with TBS (25 mM Tris HCl pH 7.5 and 125 mM NaCl), and then cell pellets were stored at -80°C overnight. Cell pellets were resuspended to an OD_600_ of 50 in Lysis Buffer (1x FastBreak cell lysis reagent (Promega), 50 μg/mL lysozyme, 1% Triton X-100, 1 mM PMSF, and 1x protease inhibitor cocktail; 100x inhibitor cocktail contained the following: 0.07 mg/mL phosphoramidon (Santa Cruz), 0.006 mg/mL bestatin (MPbiomedicals/Fisher Scientific), 1.67 mg/mL AEBSF (DOT Scientific), 0.07 mg/mL pepstatin A (Gold Bio), 0.07 mg/mL E64 (Gold Bio)). Cells were lysed with a QSonica Q55 tip probe sonicator, resulting in a DNA shear size of ∼500 bp. Lysates were clarified by centrifugation and then diluted 5-fold in IP Buffer (50 mM HEPES NaOH pH 7.5, 150 mM NaCl, 1 mM EDTA, and 1% Triton X-100). Diluted lysates were applied to Anti-FLAG M2 Magnetic Beads (Sigma) equilibrated in IP buffer. Lysates were incubated with beads at room temperature with end-over-end mixing for 2 hours. Beads were then washed twice with IP Buffer, once with Wash Buffer 1 (50 mM HEPES NaOH pH 7.5, 1 mM EDTA, 1% Triton X-100, 500 mM NaCl, and 0.1% SDS), once with Wash Buffer 2 (10 mM Tris HCl pH 8.0, 250 mM LiCl, 1 mM EDTA, 0.5% NP-40, and 1% Triton X-100), and finally once with TE (10 mM tris pH 8.0 and 1 mM EDTA). Bound protein-DNA complexes were eluted off the beads by incubation with Elution Buffer (50 mM Tris HCl pH 7.5, 10 mM EDTA, and 1% SDS) at 65°C for 30 minutes. Samples were digested with 20 μg Proteinase K in Elution Buffer for 2 hours at 42°C, then crosslinks were reversed by incubating samples at 65°C for 6 hours.

DNA samples were cleaned up and used as template for quantitative PCR (qPCR) using iTaq Universal SYBR Green Supermix (Bio-Rad) and primers specific for the genes indicated (primers are listed in **Table S3**) on a Step-One qPCR system. Standard curves of genomic DNA were included in each experiment and were used to determine the abundance of each amplicon in the input (derived from the lysate prior to ChIP) and output (derived from the samples after ChIP). Primers to amplify *rpoB* served as a baseline control in this assay because ChiS does not bind this locus. Data are reported as ‘Fold Enrichment’, which is defined as the ratio of P_*chb*_/*rpoB* DNA found in the output divided by the ratio of P_*chb*_/*rpoB* DNA found in the input.

### Protein purification

For purification of ChiS^Rec-C^ (which contained an N-terminal 6x-histidine tag), *E. coli* BL21 harboring the vector of interest was grown shaking at 37°C in LB supplemented with 100 μg/mL carbenicillin until an OD_600_ of 0.6 was reached. Protein expression was induced by the addition of IPTG to a final concentration of 100 μM, and cells were grown shaking at 30°C for an additional 4 hours. Cell pellets were stored at -80°C overnight. Pellets were resuspended in Buffer A (50 mM Tris HCl pH 7.5, 500 mM NaCl, 1 mM DTT, 20 mM imidazole, and 10% glycerol) supplemented with 2 mg/mL DNaseI, 1 mM PMSF, and 1 mg/mL lysozyme, then incubated rocking at room temperature for 20 minutes. Cells were lysed by French Press, clarified by centrifugation, and then applied to a HisTrap HP Nickel column (GE Healthcare). The column was washed with Buffer A, and then the protein was eluted using a gradient with Buffer B (50 mM Tris HCl pH 7.5, 500 mM NaCl, 1 mM DTT, 500 mM imidazole, and 10% glycerol). Fractions were stored at -80°C in single use aliquots, which were subsequently used for EMSAs.

For purification of MBP-ChiS^cytoplasmic^ isoforms, *E. coli* BL21 harboring the vectors of interest were grown at 37°C with shaking in LB supplemented with 100 μg/mL carbenicillin to an OD_600_ of 0.6. Protein expression was induced by the addition of IPTG to a final concentration of 1 mM and cells were grown shaking at 30°C for an additional 4 hours. Cell pellets were stored at -80°C overnight. Pellets were resuspended in Column Buffer (20 mM Tris HCl pH 8.0, 200 mM NaCl, 1 mM EDTA, and 1 mM DTT) supplemented with 2 mg/L DNaseI, 1 mM PMSF, and 1 mg/mL lysozyme. Lysis reactions were incubated rocking at 4°C for 20 minutes, then cells were lysed by sonication. Lysates were clarified by centrifugation and then applied to amylose resin (NEB). Resin was washed with 10x column volumes of Column Buffer, and then protein was eluted with Elution Buffer (20 mM Tris HCl pH 8.0, 200 mM NaCl, 1mM EDTA, 1 mM DTT, and 10 mM maltose). Protein was exchanged into Storage Buffer (20 mM tris HCl pH 8.0, 25 mM KCl, 1 mM DTT, 10% glycerol) using PD-10 desalting columns (GE Healthcare). Buffer exchanged protein was stored in aliquots at -80°C.

The α-subunit of RNA Polymerase from *Vibrio harveyi* was purified exactly as previously described (54).

### Electrophoretic mobility shift assays (EMSAs)

Binding reactions contained 10 mM Tris HCl pH 7.5, 1 mM EDTA, 10 mM KCl, 1 mM DTT, 50 μg/mL BSA, 0.1 mg/mL salmon sperm DNA, 5% glycerol, 1 nM of a Cy5 labeled DNA probe, and ChiS^Rec-C^ at the indicated concentrations (diluted in 10 mM Tris pH 7.5, 10 mM KCl, 1 mM DTT, and 5% glycerol). Reactions were incubated at room temperature for 20 minutes in the dark, then electrophoretically separated on polyacrylamide gels in 0.5x Tris Borate EDTA (TBE) buffer at 4°C. Gels were imaged for Cy5 fluorescence on a Typhoon-9210 instrument.

Short DNA probes (30-60bp) were made by end-labeling one primer of a complementary pair (primers listed in **Table S3**) using 20 μM Cy5-dCTP and Terminal deoxynucleotidyl Transferase (TdT; Promega). Complementary primers (one labeled with Cy5 and the other unlabeled) were annealed by slow cooling at equimolar concentrations in annealing buffer (10 mM Tris pH 7.5 and 50 mM NaCl). P_*chb*_ and P_VCA0053_ probes were made by Phusion PCR, where Cy5-dCTP was included in the reaction at a level that would result in incorporation of 1–2 Cy5 labeled nucleotides in the final probe as previously described (41).

### Protein pulldown

Purified 6xHis-α-subunit of RNA Polymerase from *V. harveyi* and purified MBP-ChiS^cytoplasmic^ were incubated in Pulldown Buffer (50 mM HEPES pH 7.5, 200 mM NaCl, 0.1% Triton X-100, 1 mM DTT, and 0.1 mg/mL salmon sperm DNA) at room temperature for 30 minutes. Each protein was present at a final concentration of 2 μM. For reactions using 6xHis-α-subunit as the bait, buffers were supplemented with 13 mM imidazole. For reactions with no bait, the storage buffer for each protein was added at an equal volume as the bait protein. Protein incubation reactions were applied to amylose resin or cobalt resin (pre-equilibrated with Pulldown Buffer) and incubated rocking at room temperature for 30 minutes. After pulldown, the supernatant (FT) was reserved, then beads were washed 5 times with Pulldown Buffer. Bait protein was eluted with 500 mM imidazole (using 6x His-α-subunit as the bait) or 10 mM maltose (using MBP-ChiS^cytoplasmic^ as bait). FT and Elution samples were then subject to Western blot analysis.

### Western blot analysis

For *in vivo* sample Western blots, strains were grown as described for ChIP assays, pelleted, resuspended, and boiled in 1x SDS PAGE sample buffer (110 mM Tris pH 6.8, 12.5% glycerol, 0.6% SDS, 0.01% Bromophenol Blue, and 2.5% β-mercaptoethanol). For *in vitro* sample Western blots, protein samples were mixed with an equal volume of 2x SDS PAGE sample buffer. Proteins were separated by SDS polyacrylamide gel electrophoresis, then transferred to a PVDF membrane, and probed with rabbit polyconal α-FLAG (Sigma), rabbit polyclonal α-MBP (Sigma), or mouse monoclonal α-RpoA (Biolegend) primary antibodies. Blots were then incubated with α-rabbit or α-mouse secondary antibodies conjugated to IRdye 800CW (Licor) as appropriate and imaged using an Odyssey classic LI-COR imaging system.

### In vitro kinase assay

Purified MBP-ChiS^cytoplasmic^ alleles were diluted to 5 μM in kinase buffer (50 mM Tris HCl pH 8.0, 50 mM KCl, 5 mM MgCl2, 1 mM DTT), then kinase reactions were initiated with the addition of 200 μM ATP and 0.125 μCi/μL ATP [γ-^32^P]. Reactions were incubated at room temperature for 30 minutes, then separated by SDS polyacrylamide gel electrophoresis. Gels were dried, exposed to a Phosphoimager Screen overnight, and then imaged on a Typhoon-9210 instrument.

### Statistics

Statistical differences were assessed by one-way ANOVA followed by a multiple comparisons Tukey’s post-test using GraphPad Prism software. Statistical analyses were performed on the log-transformed data for qRT-PCR experiments. All means and statistical comparisons can be found in **Table S1.**

## Acknowledgements

We would like to thank Mark Goulian, David Grainger, Clay Fuqua, Daniel Kearns, Julia van Kessel, Malcolm Winkler, and Matthew Neiditch for helpful discussions. We also thank Clay Fuqua for sharing POLAR strains. We thank Alyssa Ball and Ryan Chaparian for advice on experimental set up and for supplying reagents for protein pulldown and ChIP experiments. This work was supported by grants R35GM128674 and AI118863 from the National Institutes of Health to ABD and grant no. 19K07551 from JSPS (the Japan Society for the Promotion of Sciences) KAKENHI to SY.

## Supplemental Information for

**Figure S1.**
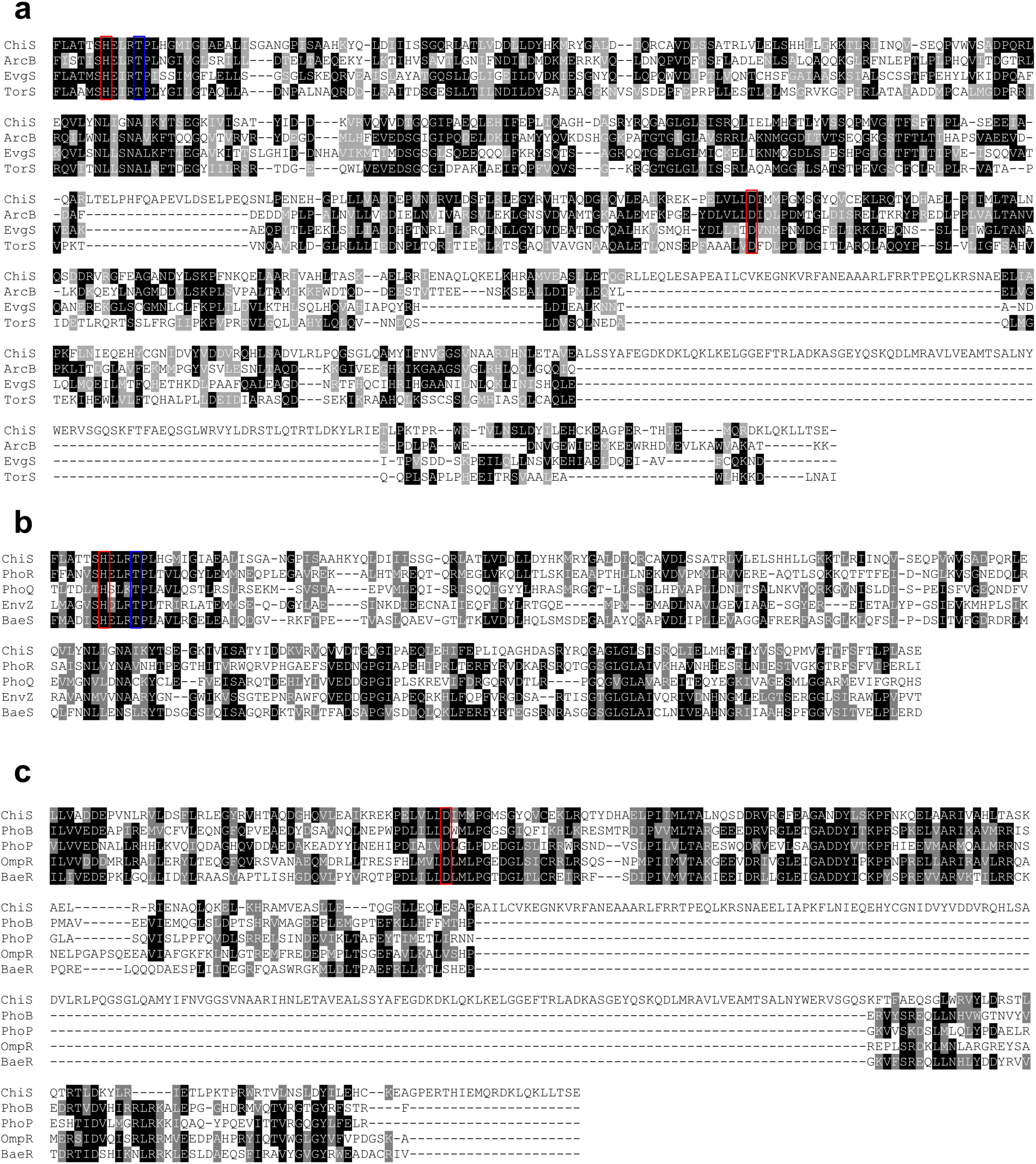
ChiS is a hybrid histidine kinase. Alignment of ChiS with (**a**) other hybrid sensor kinases (ArcB, EvgS, and TorS) starting at the H box (55), (**b**) histidine kinases (PhoR, PhoQ, EnvZ, and BaeS) starting at the H box (55), and (**c**) response regulators (PhoB, PhoP, OmpR, and BaeR) starting at the receiver domain. All protein sequences for alignment with *V. cholerae* ChiS are from *E. coli* MG1655. The conserved sites of phosphorylation (red boxes) and the threonine critical for phosphatase activity (blue box) are highlighted. Residues highlighted in black are identical, while those highlighted in gray are similar.

**Figure S2.**
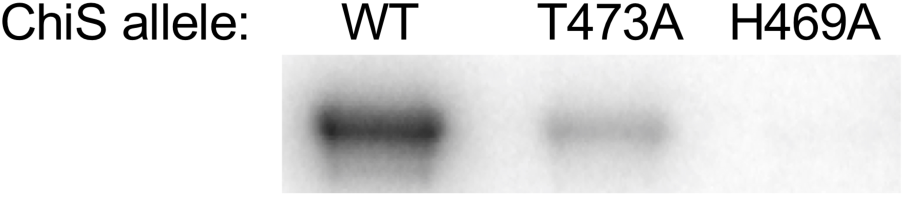
ChiS exhibits kinase activity in vitro. Purified ChiS (MBP-ChiS^cytoplasmic^) containing the indicated mutations were assessed for autokinase activity *in vitro* using ^32^P-labelled ATP. Data are representative of two independent experiments.

**Figure S3.**
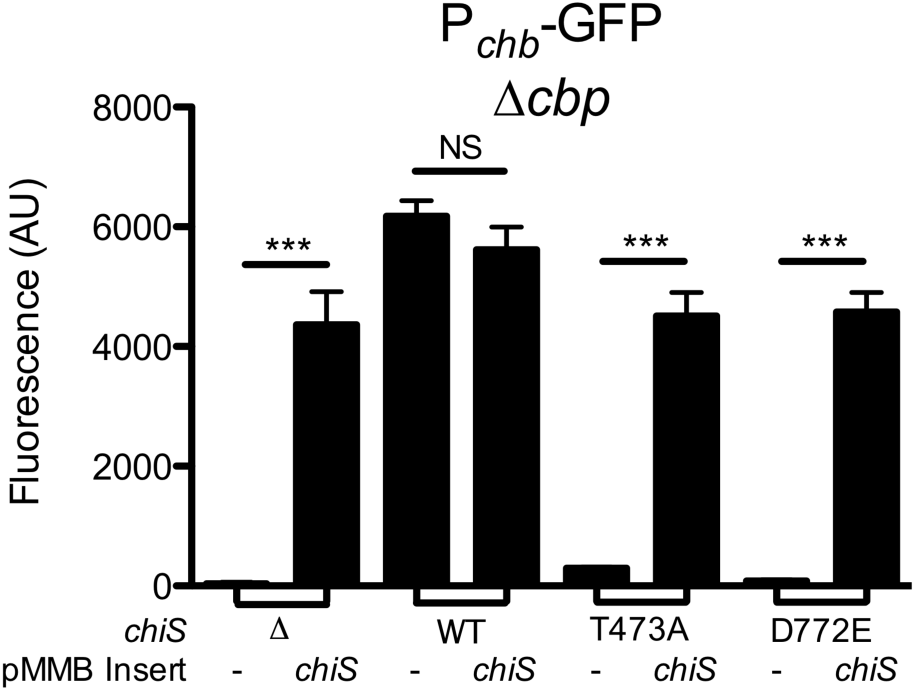
Complementation of ChiS inactive alleles with ChiS^WT^. Strains with P_*chb*_-GFP and Δ*cbp* expressing the indicated ChiS alleles (from **Fig. 1a**) were complemented with the pMMB67EH+riboswitchE vector harboring no insert (-) or ChiS^WT^ (*chiS*). Complemented strains were then assessed for P_*chb*_-GFP expression. Data are from at least three independent biological replicates and shown as the mean ± SD. Statistical comparisons were made by one-way ANOVA with Tukey’s post-test. NS, not significant. ***, p < 0.001.

**Figure S4.**
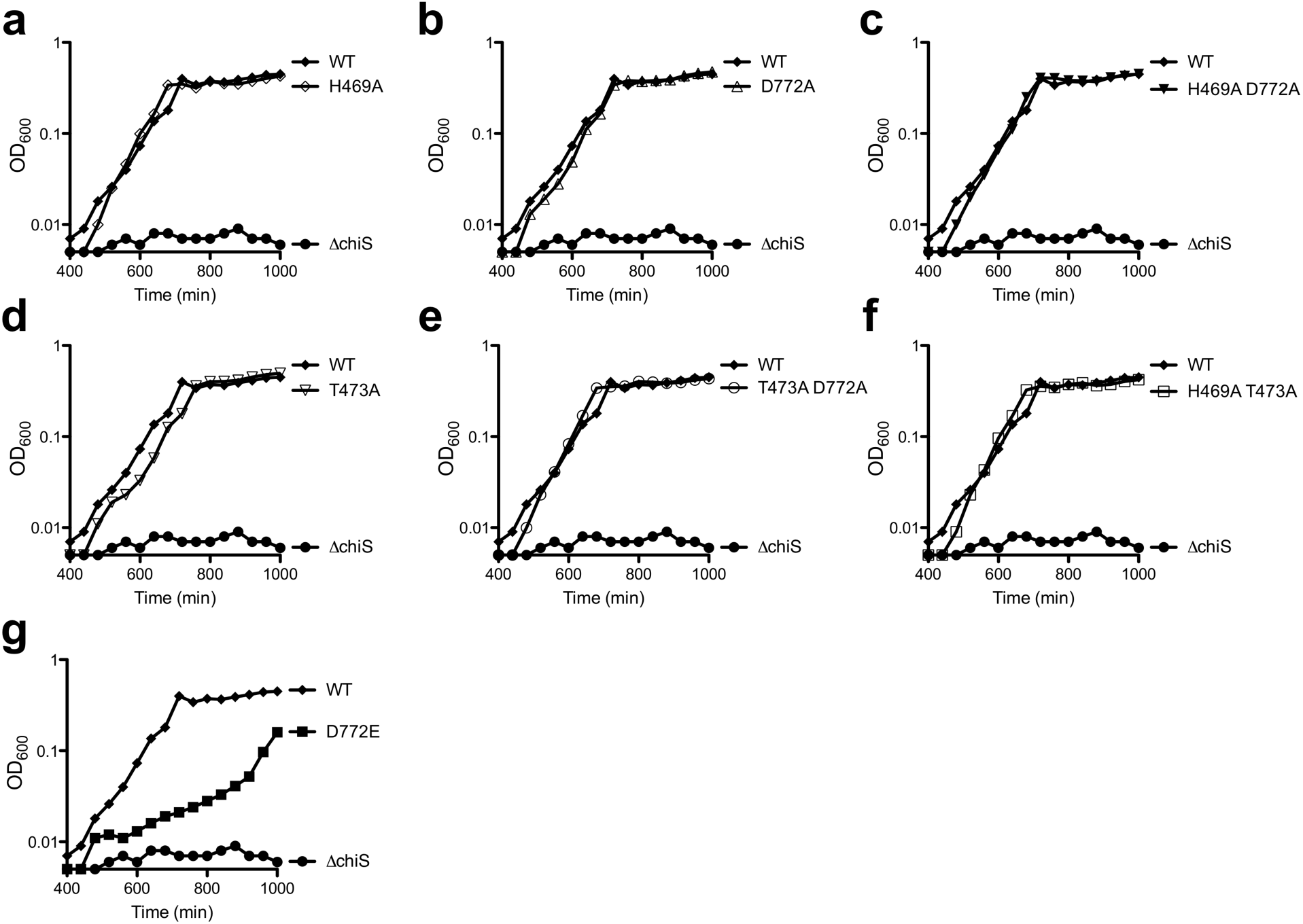
Growth of various ChiS point mutants on chitobiose. Strains harboring the indicated point mutations in ChiS were tested for growth on M9 minimal medium containing chitobiose as the sole carbon source. Data for WT and Δ*chiS* are identical in each panel and are included in for ease of comparison. The lower limit of the Y-axis is the limit of detection for this assay (OD_600_ = 0.005). Data are representative of two independent experiments.

**Figure S5.**
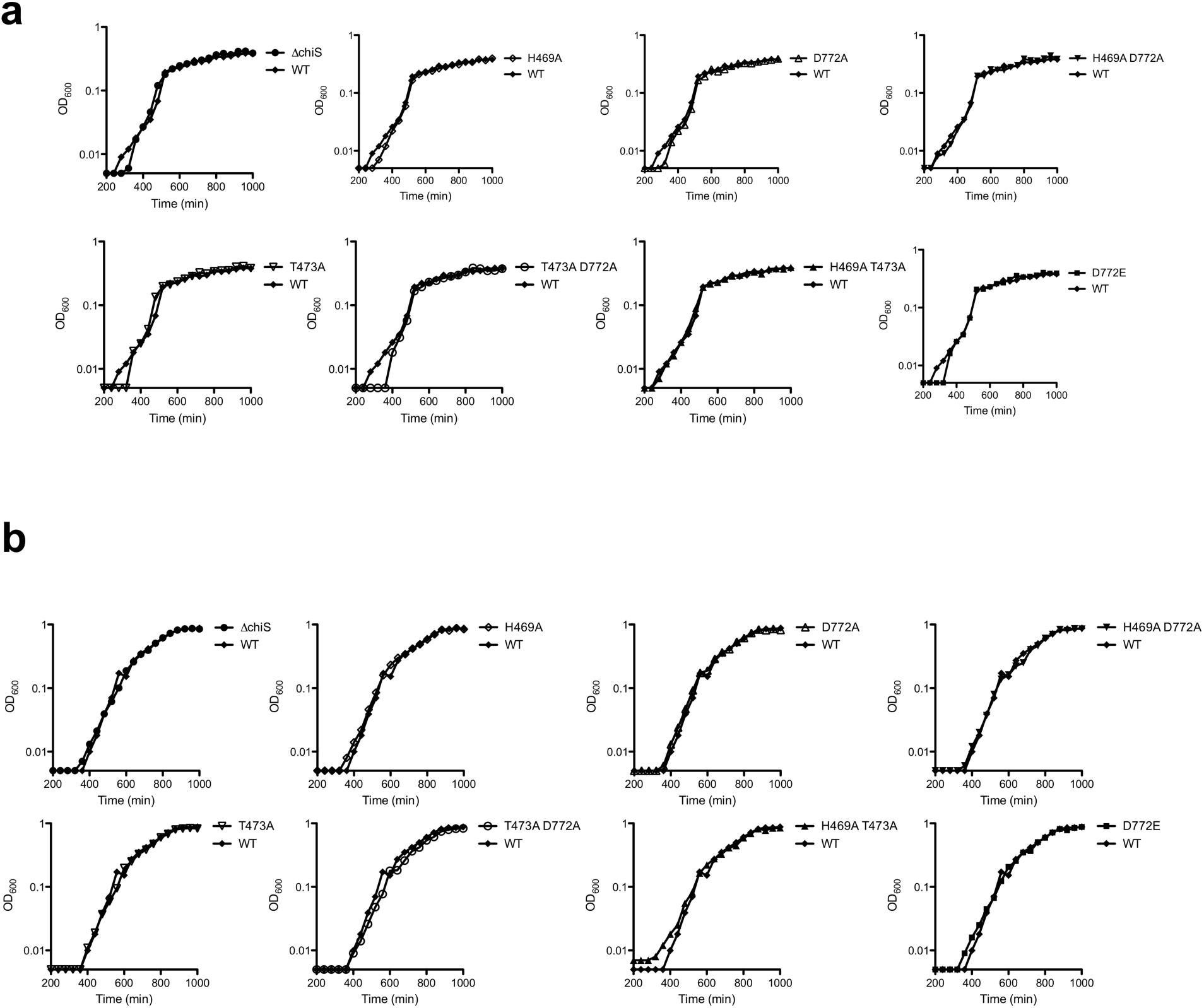
Mutations in ChiS do not alter growth on other carbon sources. Strains harboring the indicated mutations in ChiS were tested for growth on minimal medium containing tryptone (**a**) or glucose (**b**) as the sole carbon source. The lower limit of the Y-axis is the limit of detection for this assay (OD_600_ = 0.005). Data are representative of two independent experiments.

**Figure S6.**
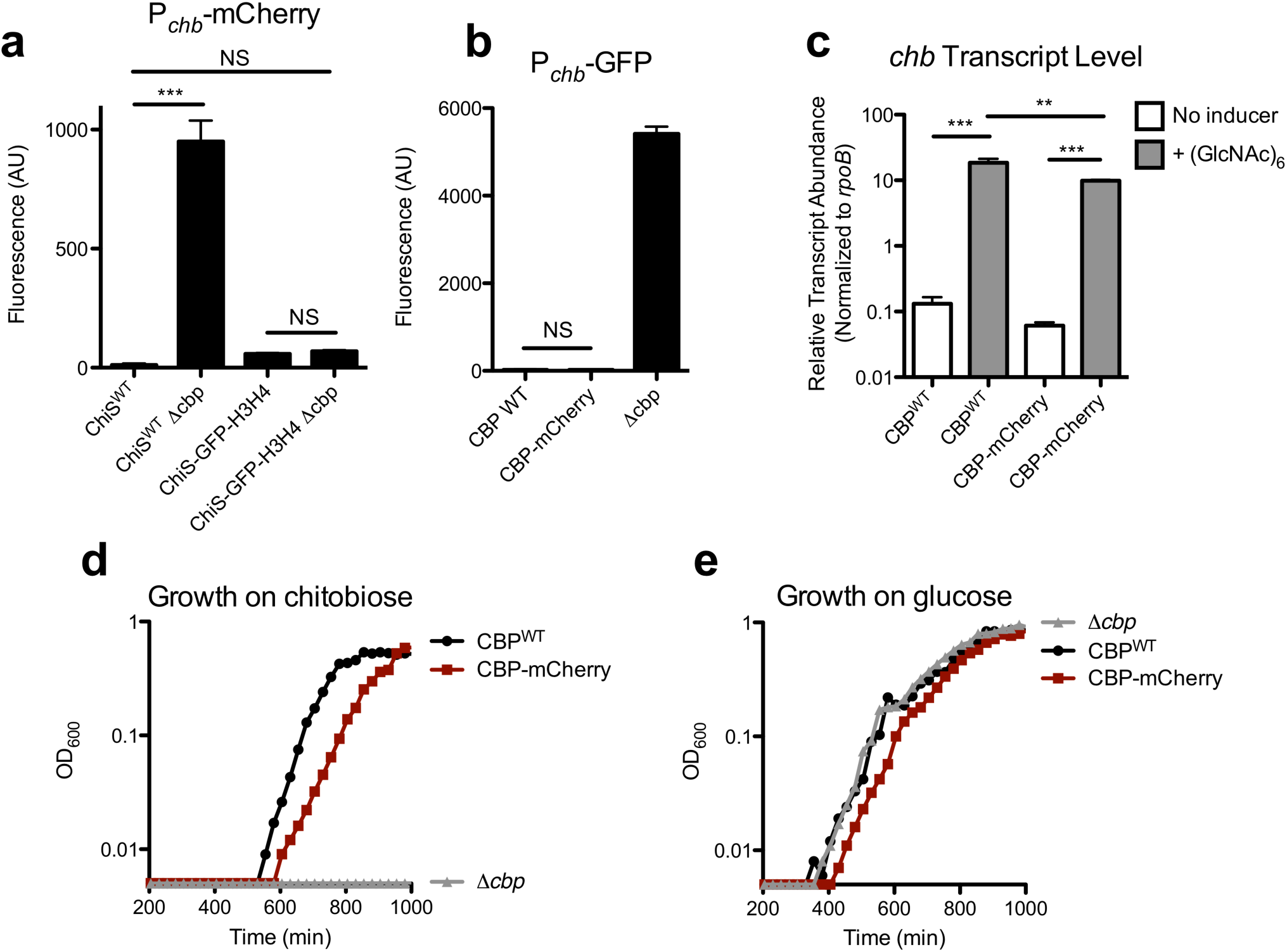
Functionality of fluorescently tagged ChiS and CBP for POLAR assays. (**a**) Strains containing a P_*chb*_-mCherry transcriptional reporter the indicated ChiS alleles and *cbp* mutations were assessed for mCherry fluorescence. (**b**) Strains containing a P_*chb*_-GFP transcriptional reporter and the indicated CBP alleles were assessed for GFP fluorescence. (**c**) Strains expressing the indicated *cbp* alleles were grown in LB medium only (No inducer) or supplemented with chitin hexasaccharide (+ (GlcNAc)_6_) and *chb* transcript abundance was determined by qRT-PCR. Data for CBP^WT^ are identical to the data presented for ChiS^WT^ in **Fig. 1b** and are included here for ease of comparison. (**d-e**) Strains expressing the indicated *cbp* alleles were assessed for growth in M9 supplemented with (**d**) chitobiose or (**e**) glucose as the sole carbon source.

**Figure S7.**
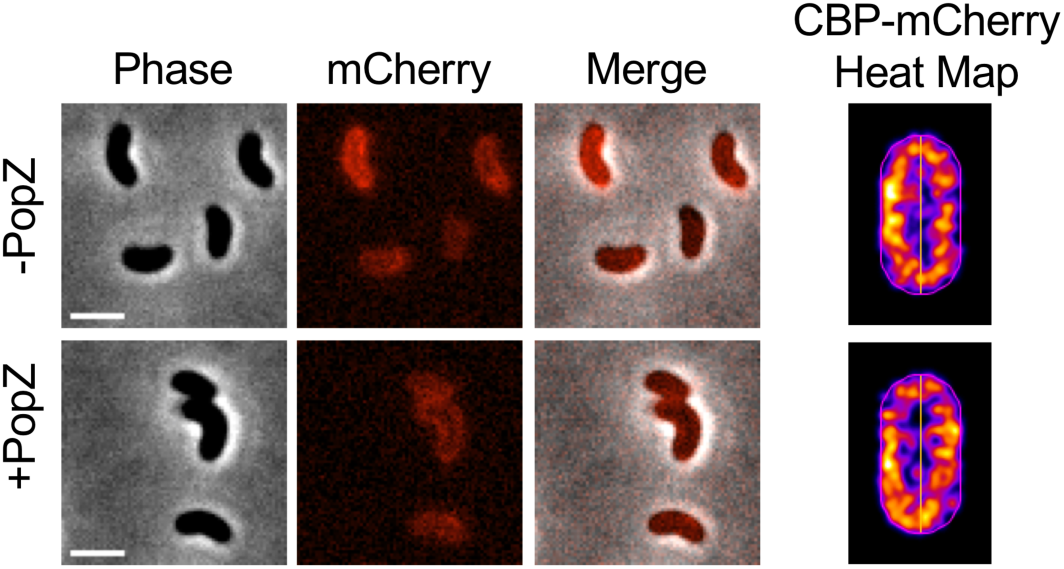
CBP is diffuse in the periplasm in the absence of ectopically expressed ChiS-GFP-H3H4. Strains containing P_*BAD*_-PopZ and CBP-mCherry were grown in LB (-PopZ) or LB supplemented with 0.05% arabinose (+PopZ) then imaged. Representative images are shown (left), and the corresponding heat maps (right) were derived from at least 500 cells.

**Figure S8.**
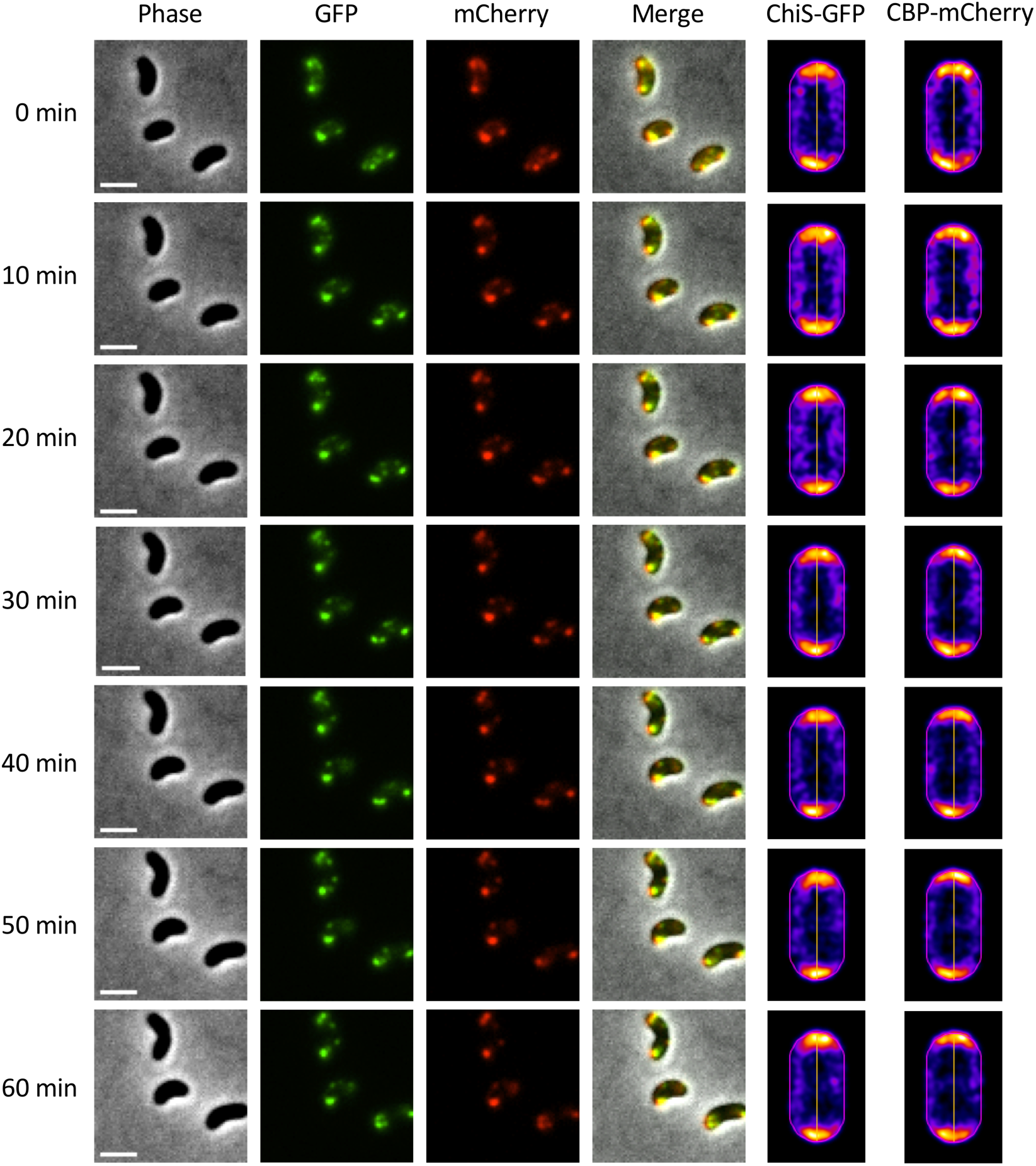
CBP does not dissociate from ChiS in the presence of chitin. Montage of time lapse imaging of a *V. cholerae* strain containing P_*tac*_-ChiS-GFP-H3H4 (H3H4 domain is required for PopZ interaction), CBP-mCherry, and P_*BAD*_-PopZ. The strain was grown with 10 µM IPTG and 0.05% arabinose to induce ChiS-GFP-H3H4 and PopZ, respectively. Chitin hexasaccharide (0.5%) was added as an inducer to cells immediately before imaging and fluorescent localization of ChiS and CBP was kinetically monitored. Representative images (left) and heat maps (right) were generated for each fluorescent channel at the indicated time points. Heat maps are representative of at least 500 cells.

**Figure S9.**
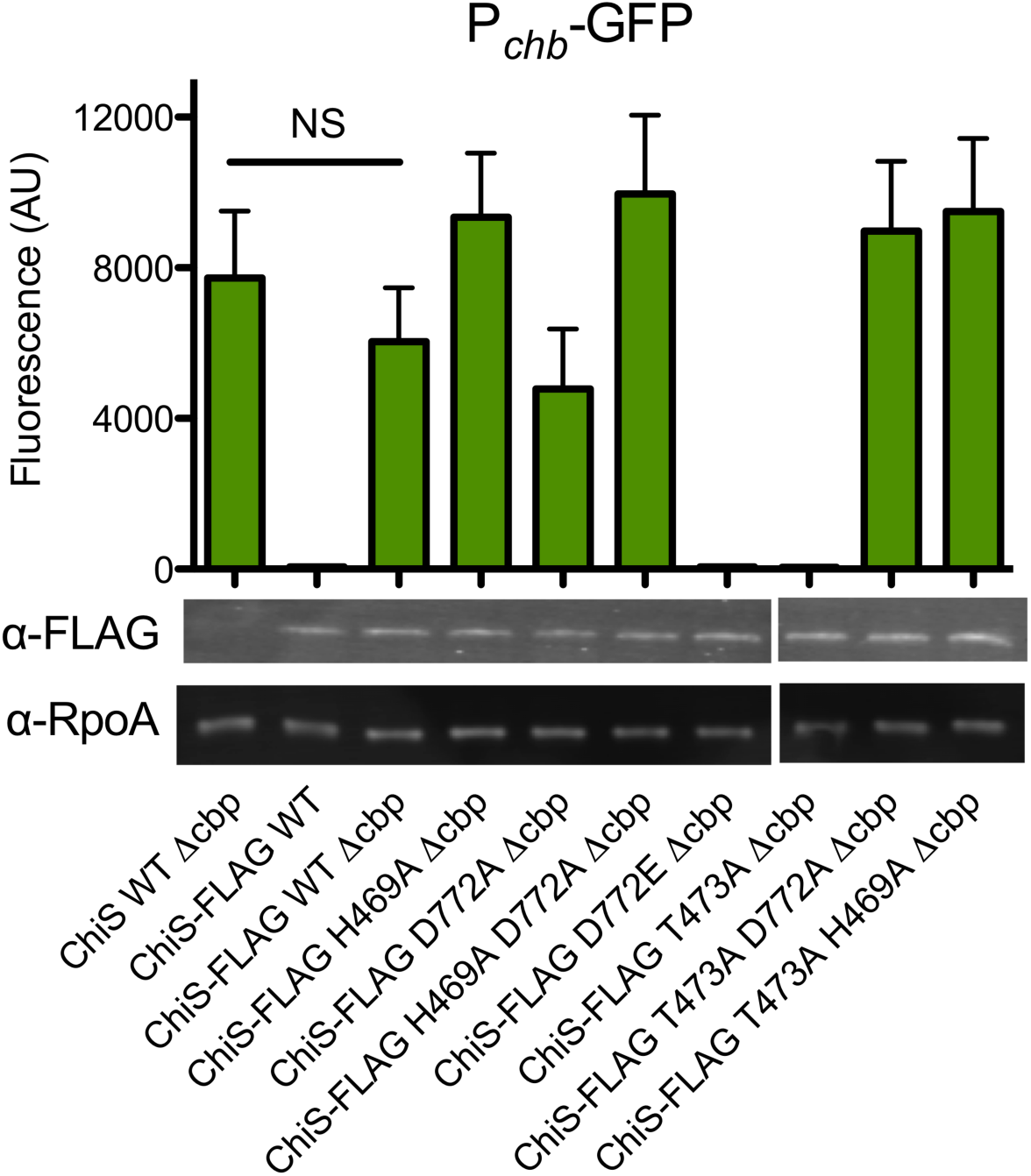
ChiS-FLAG is functional and ChiS expression does not change when residues critical for phosphorylation or phosphatase activity are mutated. FLAG-tagged ChiS alleles were assessed for functionality using a P_*chb*_-GFP reporter and expression of ChiS was assessed by Western blot analysis. P_*chb*_-GFP results confirm that ChiS-FLAG functions similar to untagged ChiS (as shown in **Fig. 1a**). Western blot results demonstrate that mutating residues critical for phosphorylation (H469, D772) or phosphatase activity (T473) do not alter ChiS protein levels. P_*chb*_-GFP data are the result of at least three independent biological replicates and shown as the mean ± SD. Western blot results are representative of two independent experiments. Statistical comparisons were made by one-way ANOVA with Tukey’s post-test. NS, not significant.

**Figure S10.**
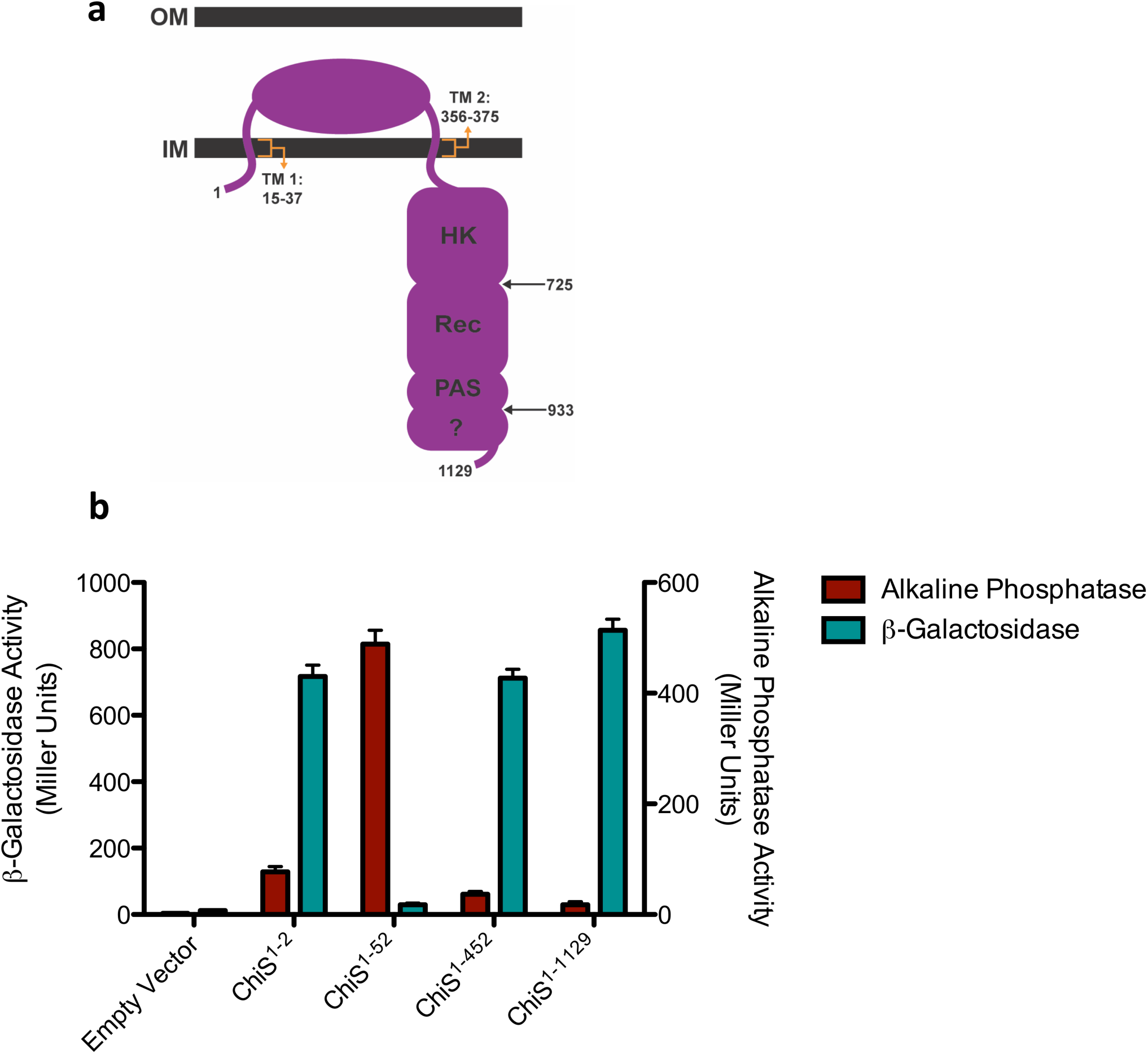
Testing ChiS membrane topology. (**a**) ChiS predicted membrane topology with transmembrane (TM), histidine kinase (HK), receiver (Rec), and Per-Arnt-Sim (PAS) domains labeled with residue numbers. (**b**) The indicated ChiS C-terminal truncations were fused to *lacZ* or *phoA* and then assessed for beta-galactosidase and alkaline phosphatase activity, respectively. All data are the result of at least three independent biological replicates and shown as the mean ± SD.

**Figure S11.**
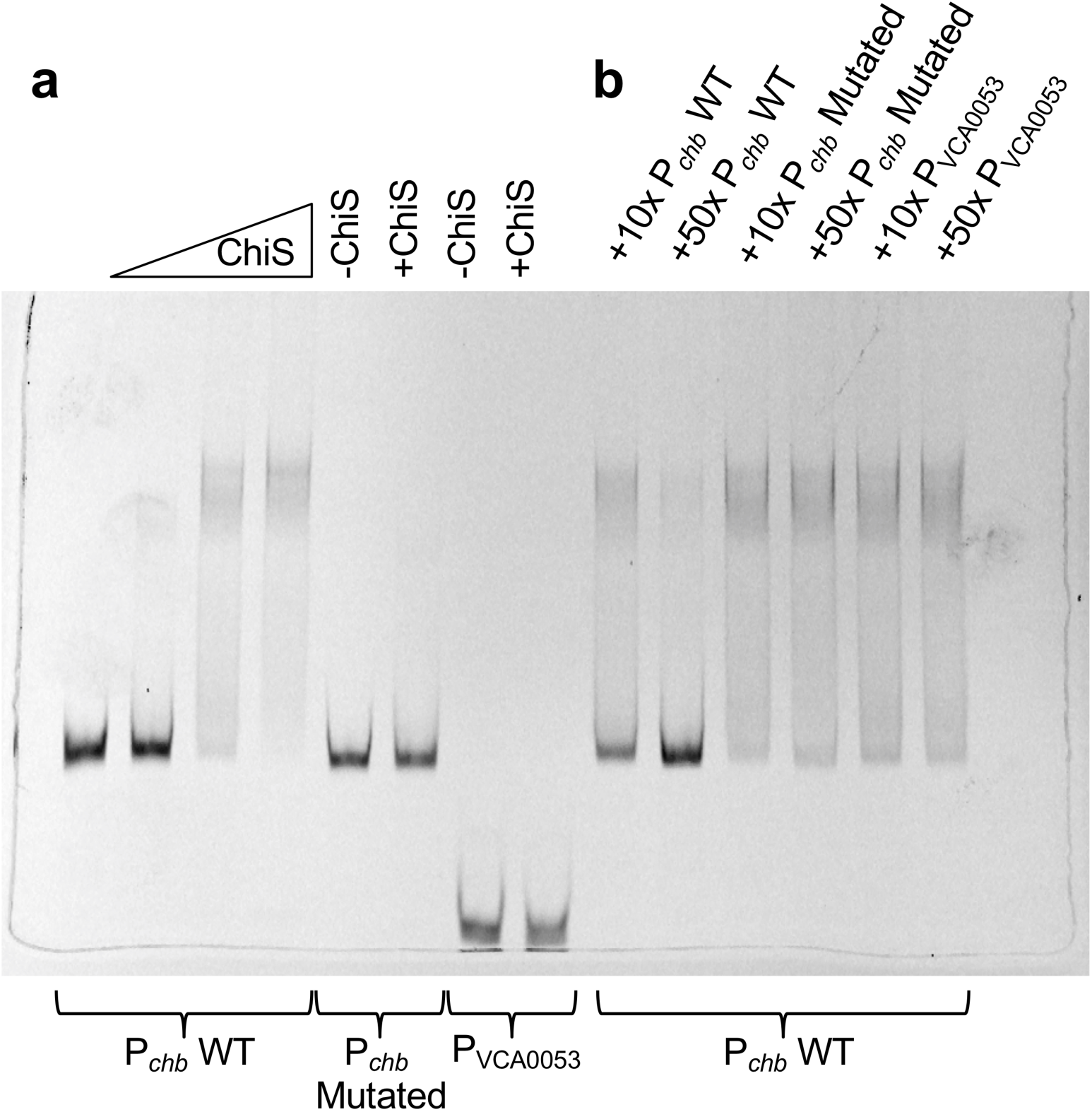
ChiS binds to the P_chb_ promoter. (**a**) EMSA assays were carried out using a purified portion of the ChiS cytoplasmic domain (ChiS^Rec-C^) and the Cy5-labeled DNA probe indicated below the gel. For lanes 1-4, a fixed concentration of the minimal fragment of the P_*chb*_ promoter required for activation (7) (P_*chb*_ WT) was incubated with increasing concentrations of ChiS (from left to right: 0 nM, 50 nM, 100 nM, 200 nM). A probe where the two ChiS binding sites within P_*chb*_ were mutated (P_*chb*_ Mutated – see **Fig. 3c** for details) and a negative control probe that represents a promoter not regulated by ChiS (P_VCA0053_) were incubated with no ChiS (-ChiS) or 200 nM ChiS (+ChiS). (**b**) Cold competitor EMSAs were carried out using reactions where 100 nM ChiS was incubated with a fixed concentration of Cy5 labeled P_*chb*_ WT probe and either a 10x or 50x fold-excess of the unlabeled competitor DNA indicated above the gel. These data indicate that ChiS binding to P_*chb*_ WT can only be competed off by specific DNA (unlabeled P_*chb*_ WT) but not nonspecific DNA (P_VCA0053_) or the probe where both ChiS binding sites in P_*chb*_ are mutated (P_*chb*_ Mutated). Data are representative of two independent experiments.

**Figure S12.**
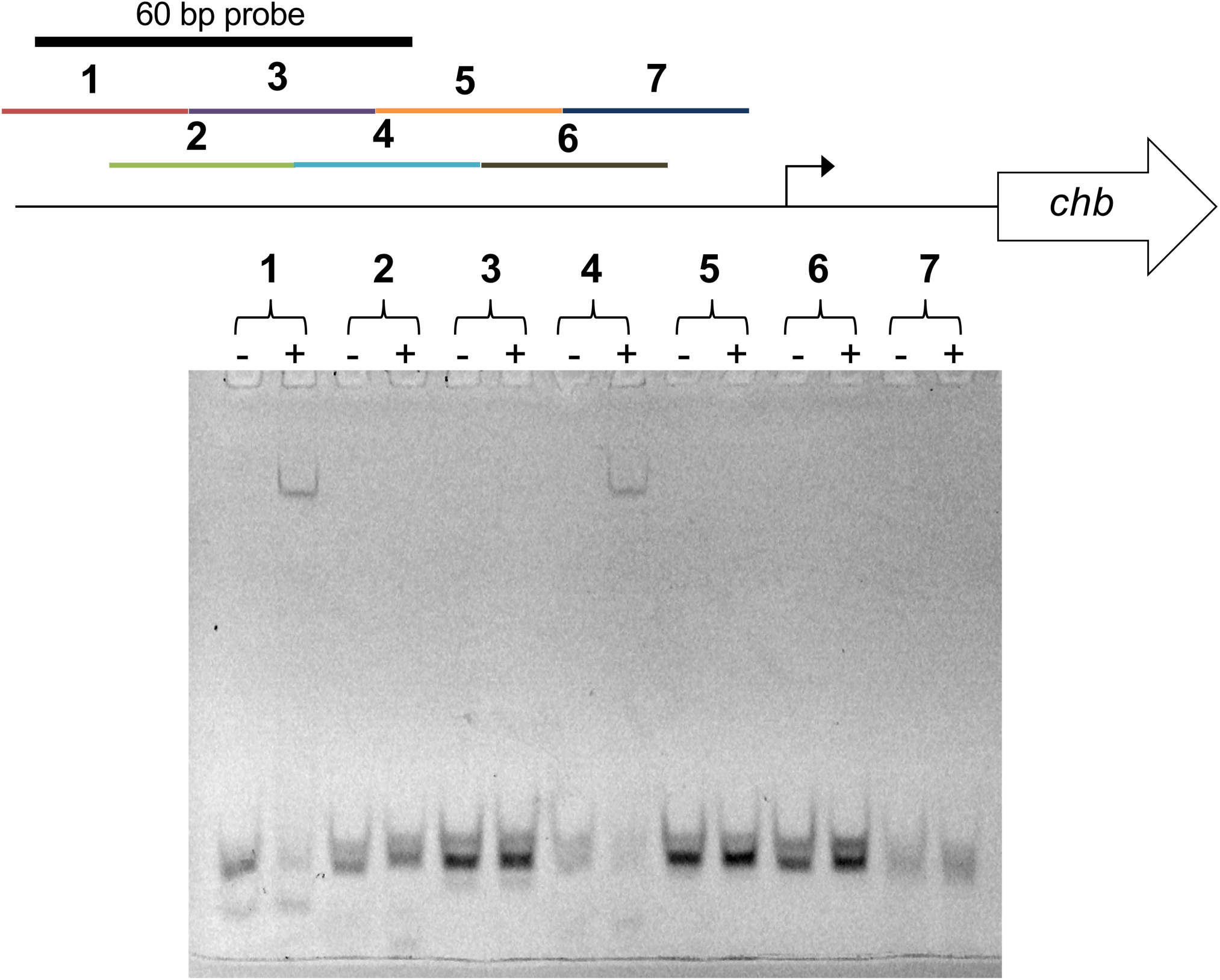
ChiS binds two sites within the P_chb_ promoter. EMSA reactions were carried out using a purified portion of the ChiS cytoplasmic domain (ChiS^Rec-C^) and Cy5 labeled 30 bp overlapping segments of P_*chb*_ (labeled 1-7). Probes were incubated with no ChiS (-) or 200 nM ChiS (+). The 60 bp probe containing both ChiS binding sites used in **Fig. 3a** is also indicated on the schematic. Data are representative of two independent experiments.

**Figure S13.**
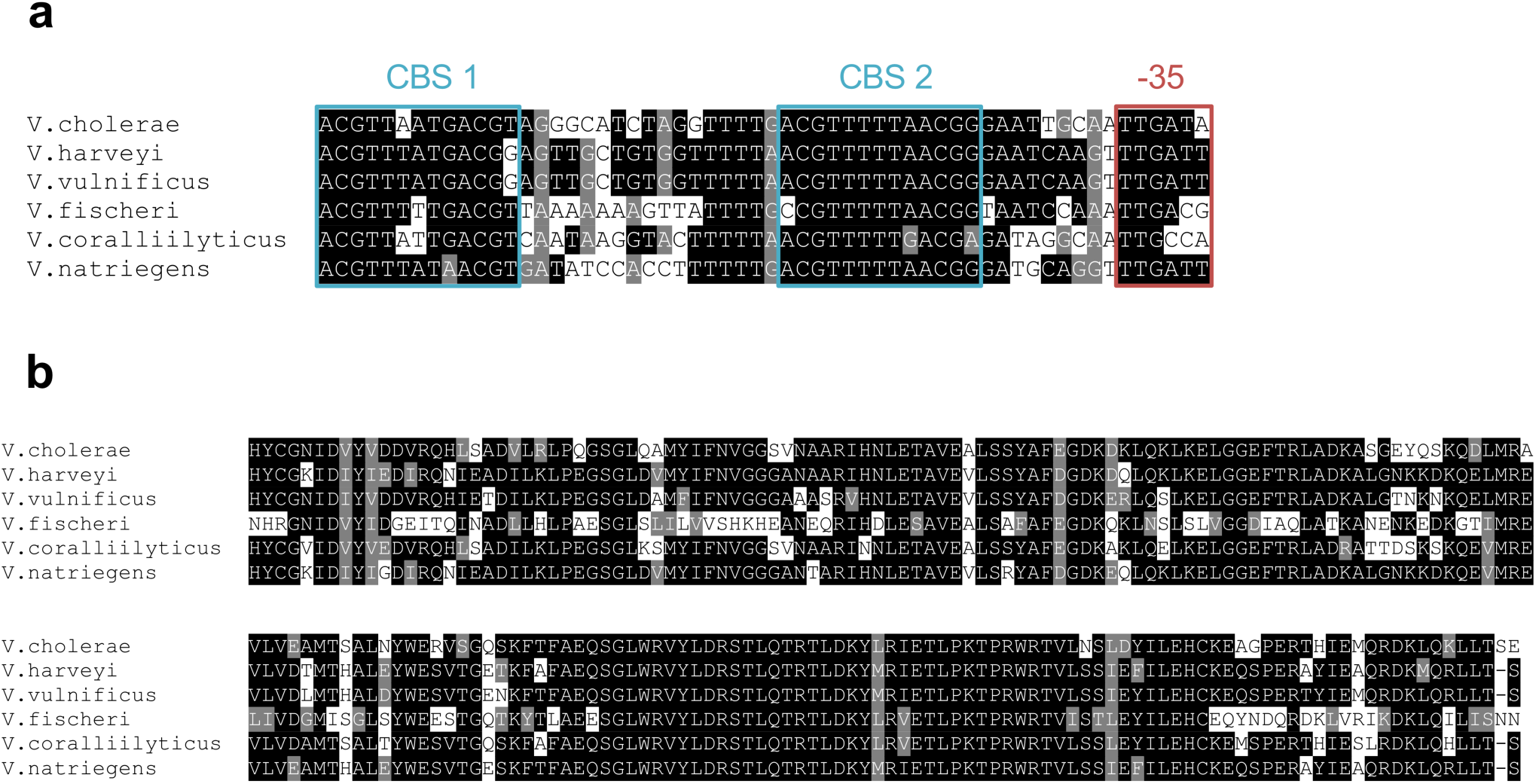
The ChiS binding sites (CBSs) in P_chb_ and the ChiS C-terminus are conserved among Vibrio species. (**a**) A section of the P_*chb*_ DNA sequence (CBS 1 until the -35 signal) from the indicated *Vibrio* species were aligned. CBSs are boxed in blue and the -35 signal is boxed in red. (**b**) The last 197 amino acids of *V. cholerae* ChiS, which contains a cryptic DNA binding domain, was aligned to the C-terminal domains of ChiS homologs from the indicated *Vibrio* species. Residues highlighted in black are identical, those highlighted in gray are similar.

**Figure S14.**
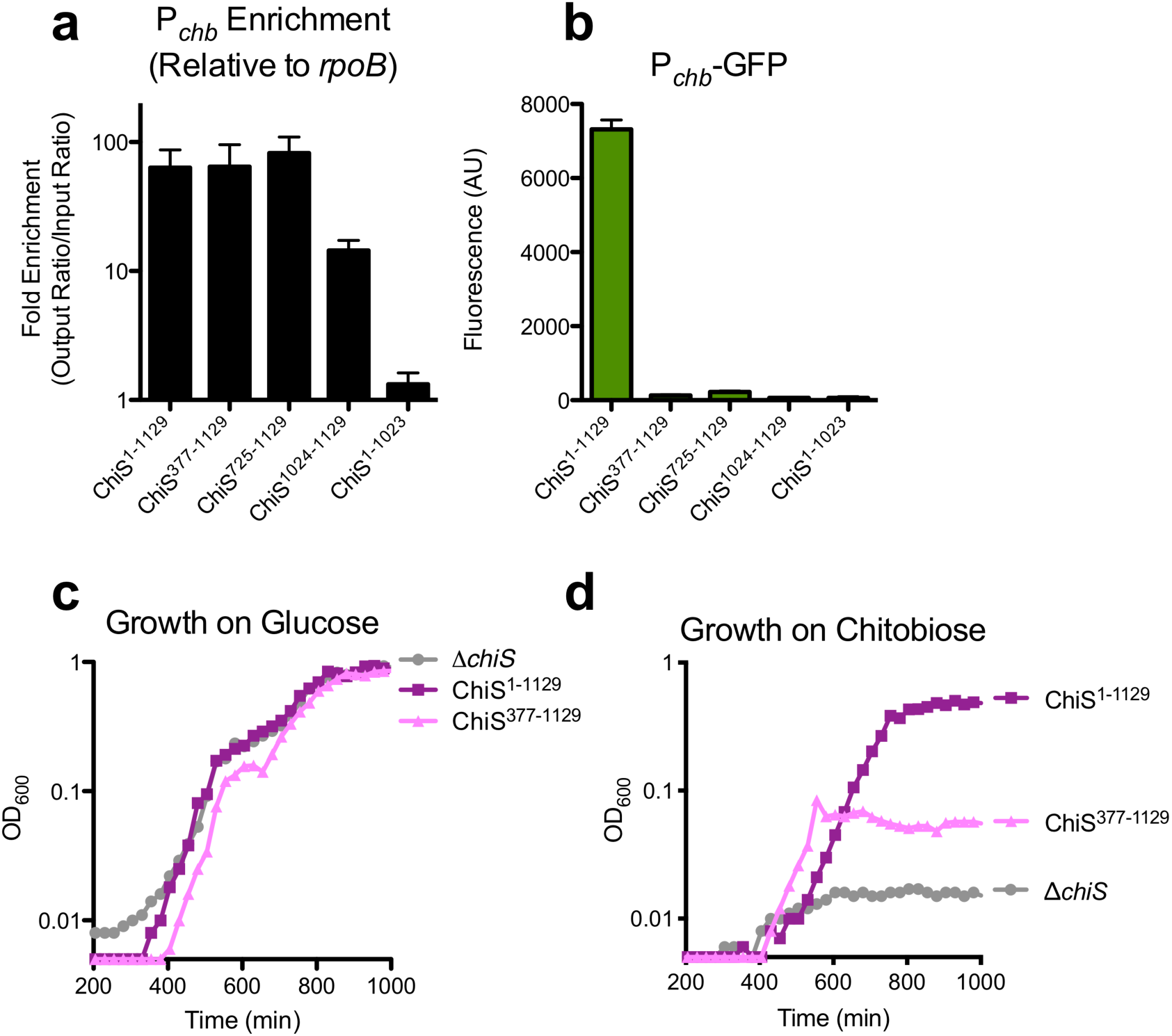
The last 106 amino acids of ChiS is necessary and sufficient to bind P_chb_, but does not activate P_chb_. (**a**) ChIP assays were carried out in *V. cholerae* strains overexpressing N-terminal truncations of ChiS (first 4 bars) or natively expressing ChiS with a C-terminal truncation of the last 106 amino acids (last bar). (**b**) The same strains indicated in **a** were assessed for activation of a P_*chb*_-GFP reporter. (**c** and **d**) Growth curves of *V. cholerae* strains expressing the indicated ChiS alleles. Strains were grown on chitobiose or glucose, as indicated. The lower limit of the Y-axis is the limit of detection for this assay (OD_600_ = 0.005). Data in **a** and **b** are from at least three independent biological replicates and shown as the mean ± SD. The data for ChiS^1-1129^ in **a** and **b** are identical to the data shown for “ChiS WT Δ*cbp*” in **Fig. 2b** and are included here for ease of comparison. Data in **c** and **d** are representative of two independent experiments.

**Figure S15.**
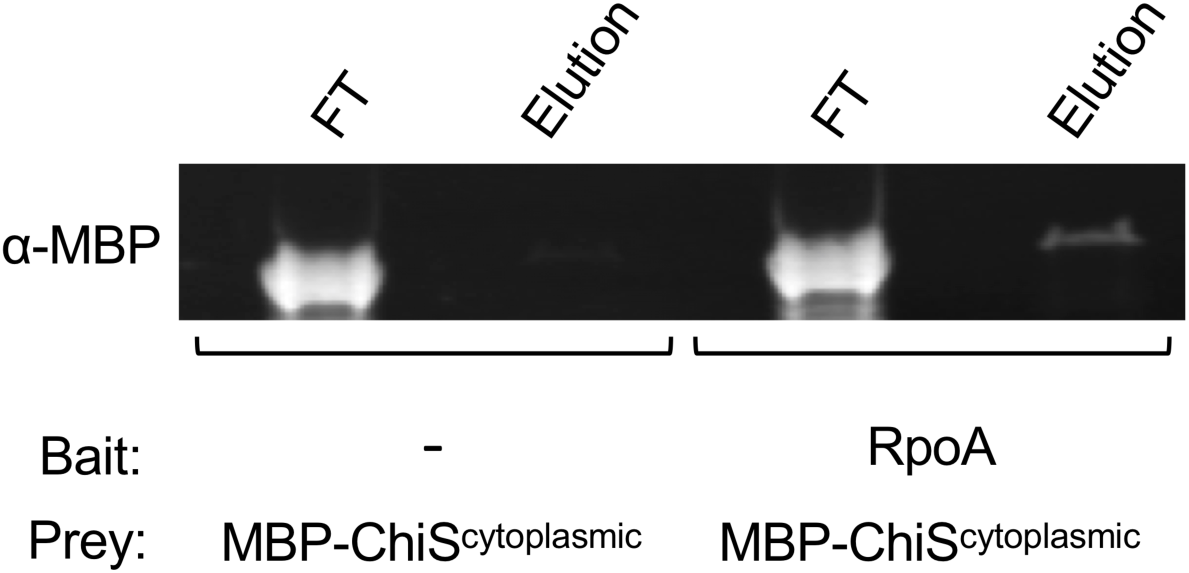
ChiS and RpoA interact in vitro. A protein pulldown using purified RpoA as bait and purified ChiS (MBP-ChiS^cytoplasmic^) as prey was carried out. The presence of the MBP-ChiS^cytoplasmic^ prey was assessed in the flow through (FT) and elution by Western blot analysis. Data are representative of two independent experiments.

**Figure S16.**
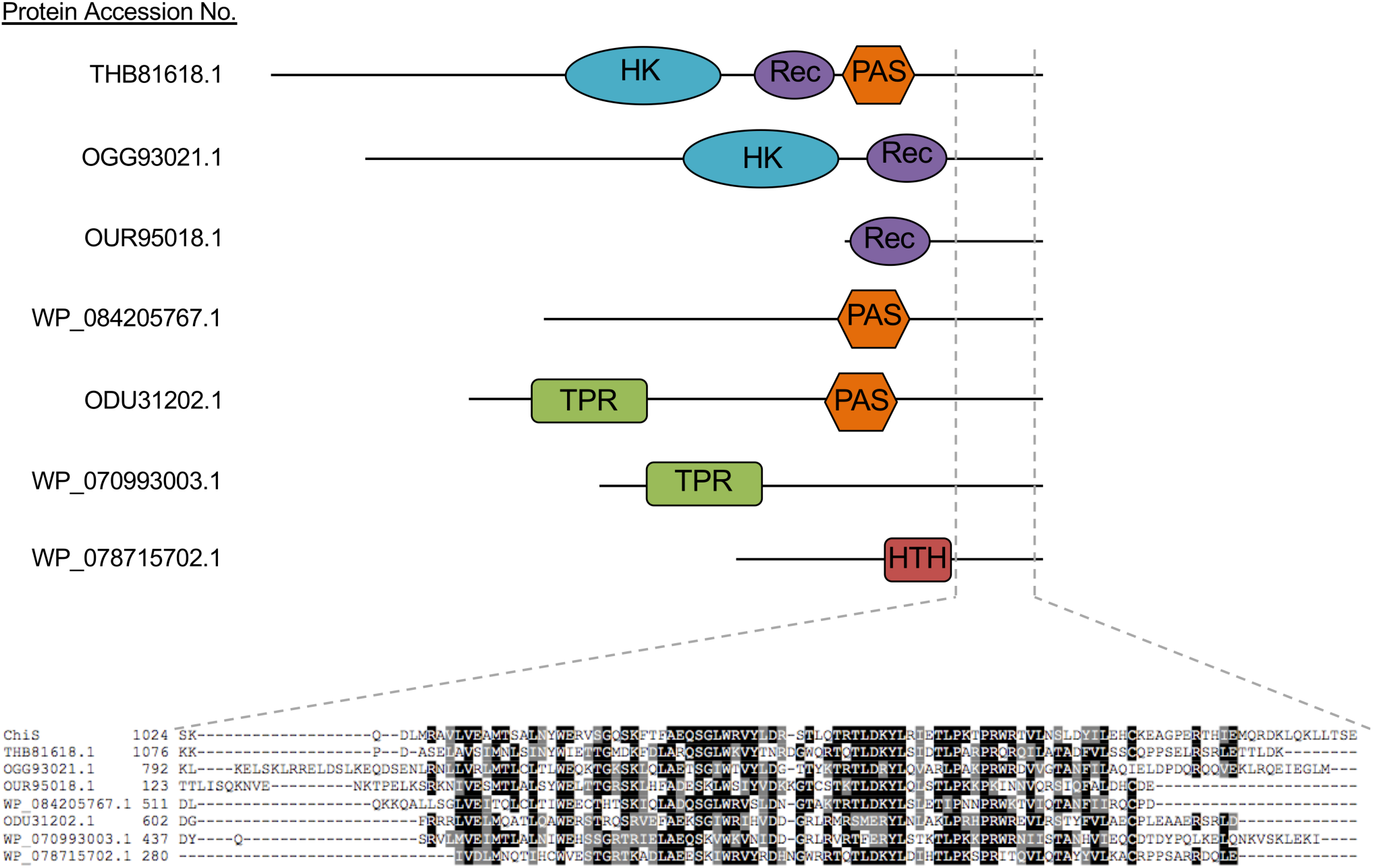
Representative examples of BLAST hits containing putative ChiS-like C-terminal DNA binding domains. Various hits from BLAST hits gave proteins that contain histidine kinase (HK), receiver (Rec), Per-ARNT-Sim (PAS), tetratricopeptide protein repeat (TPR), and helix-turn-helix (HTH) domains. The C-terminal domains of the indicated proteins were aligned against the last 106 amino acids of ChiS. Residues highlighted in black are identical, while those highlighted in gray are similar.

**Table S1.**
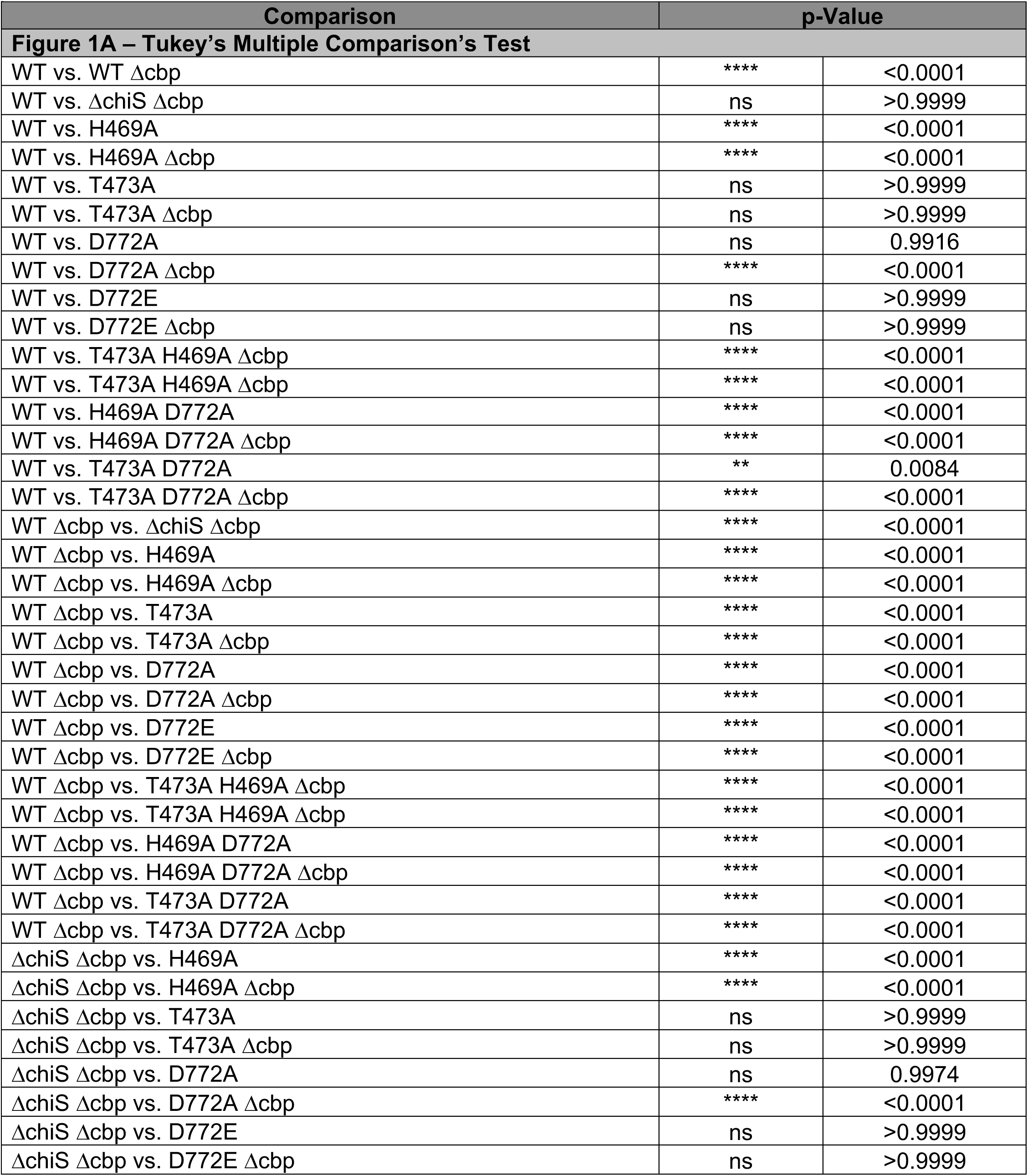
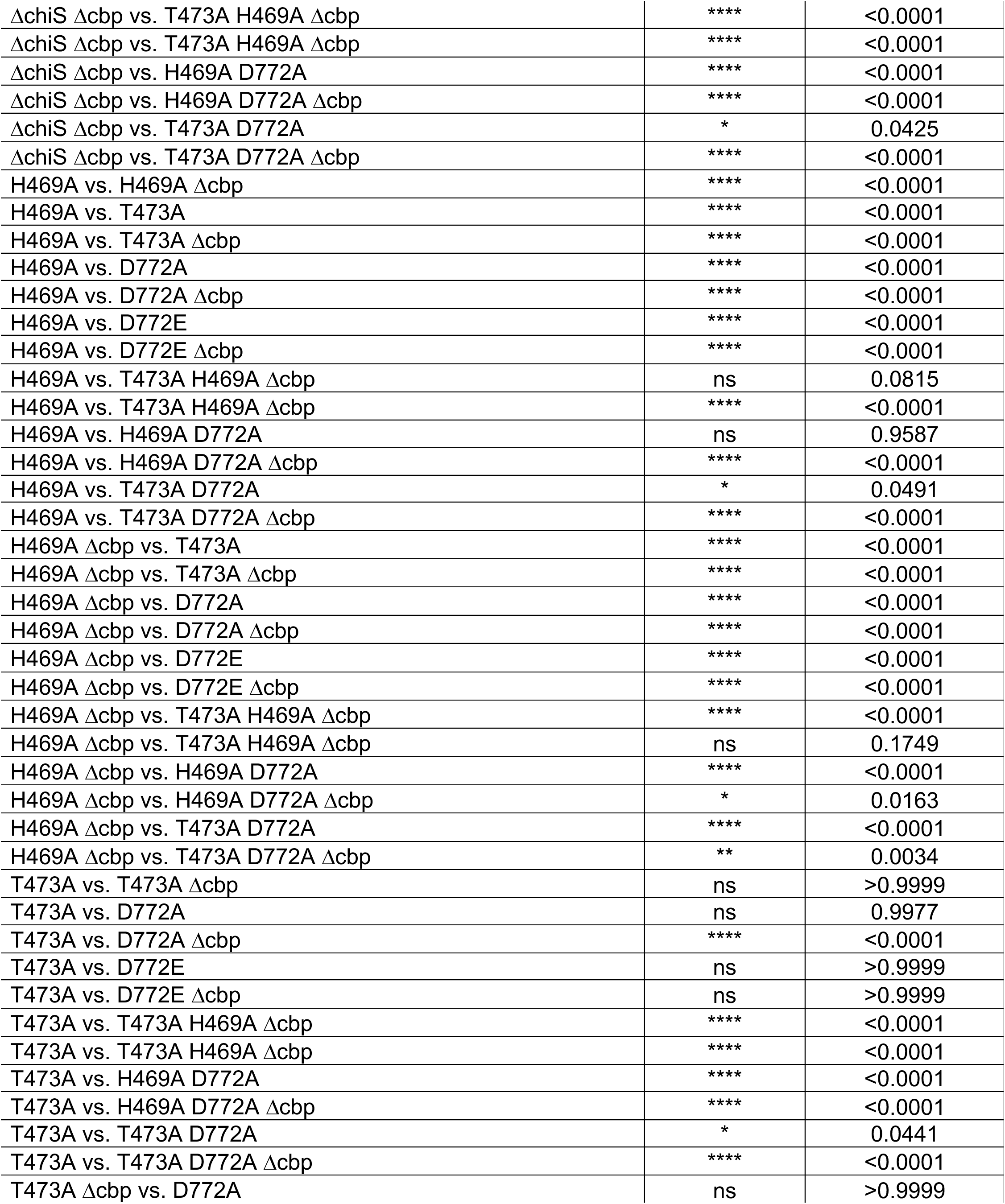
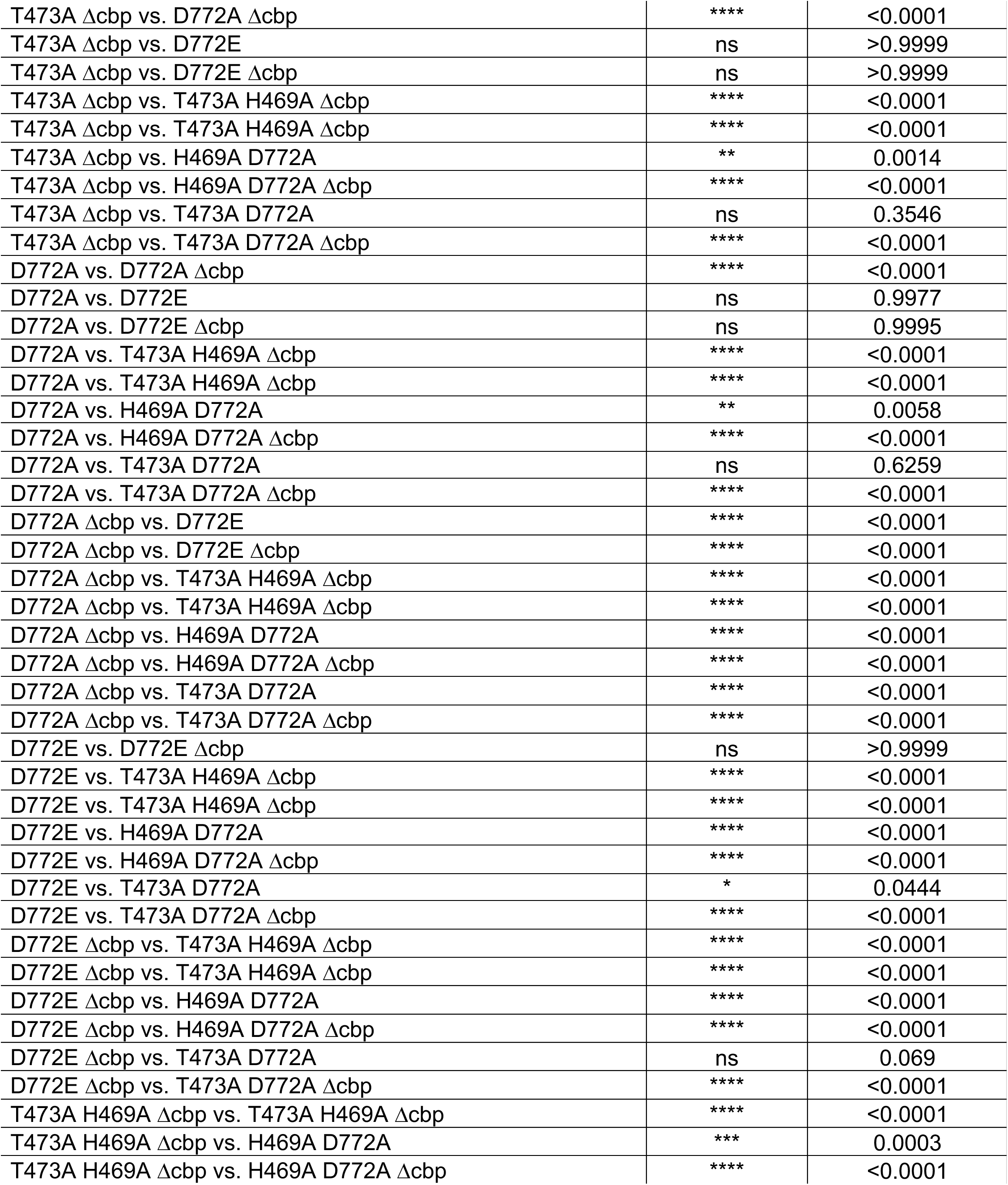
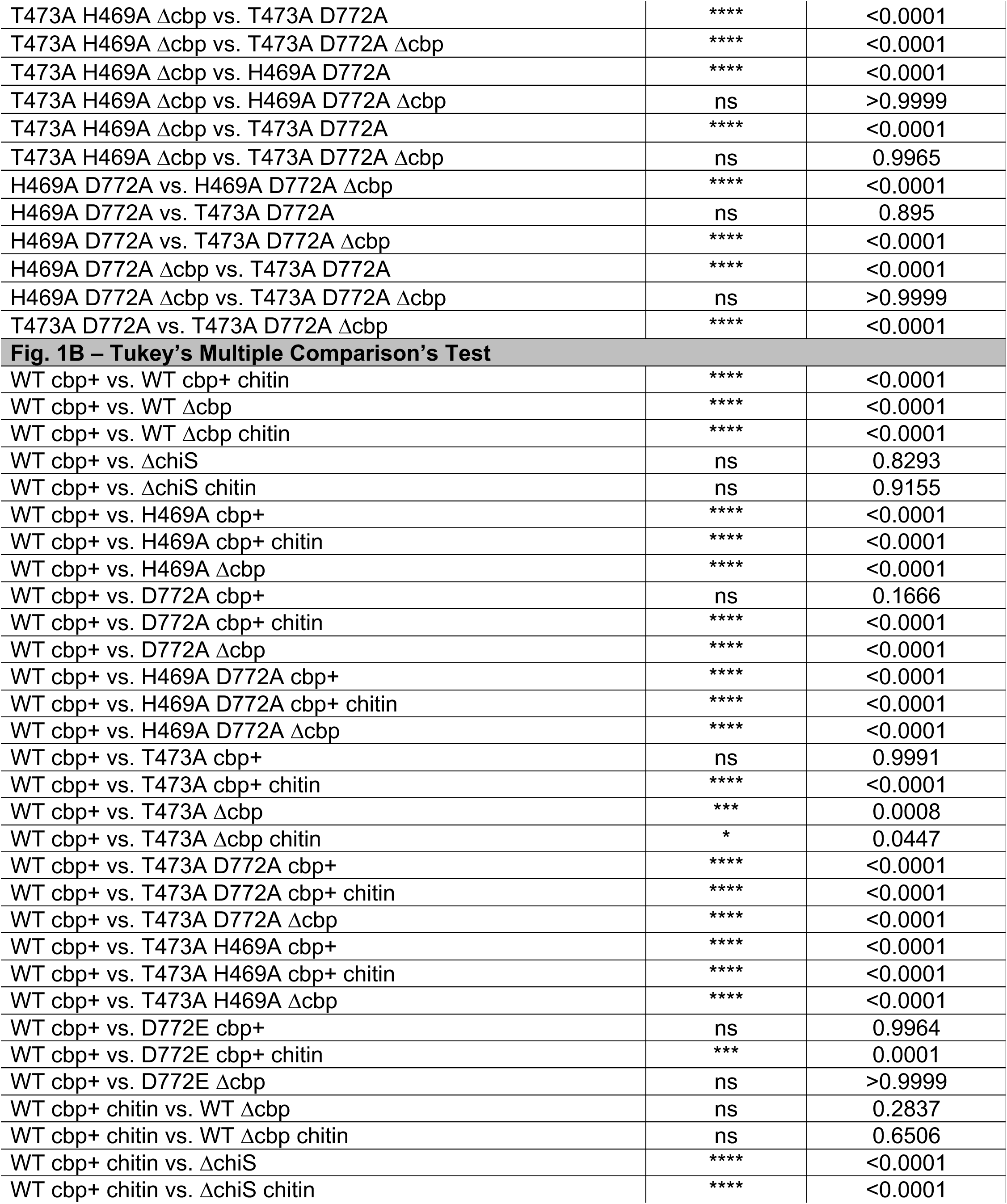
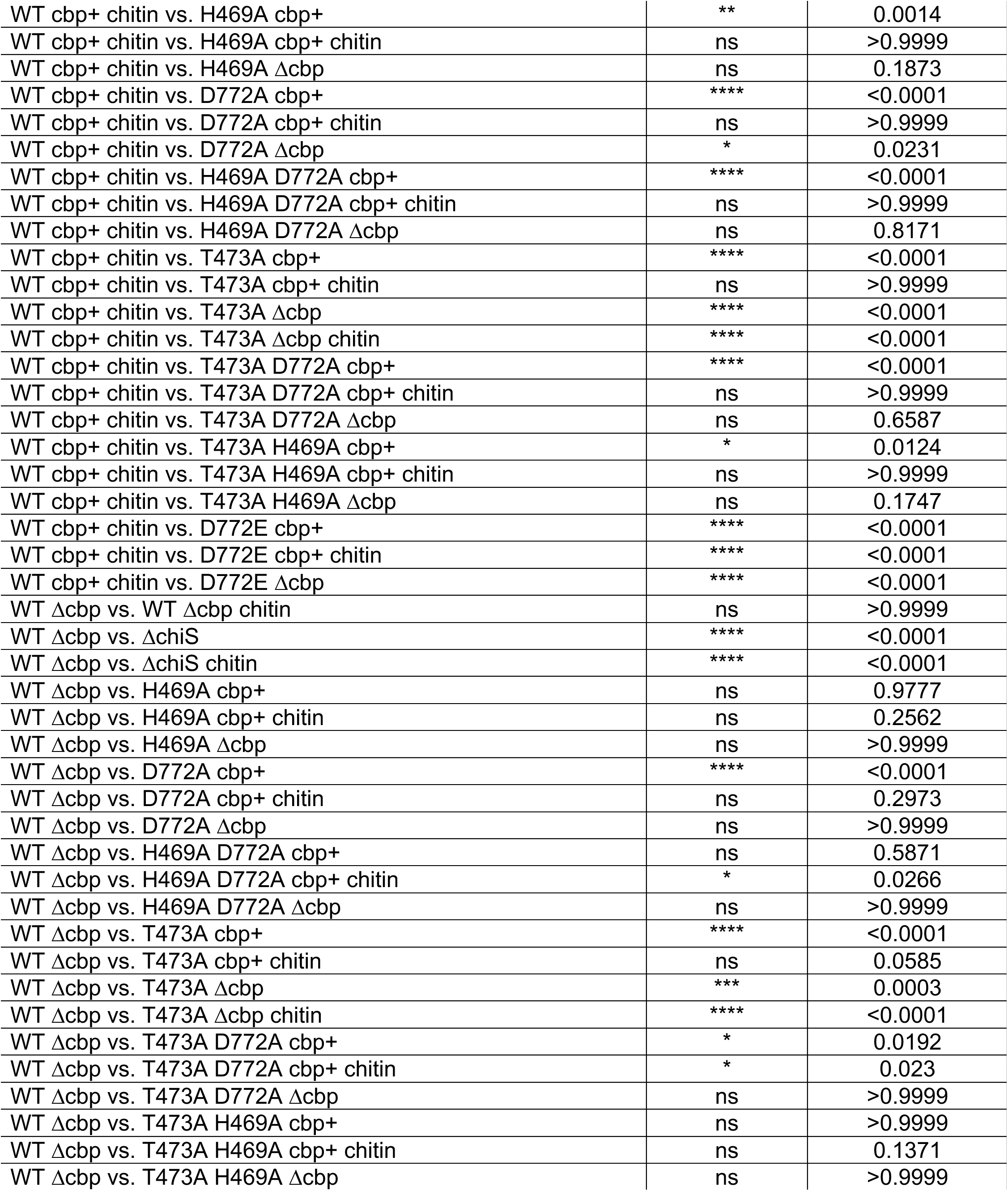
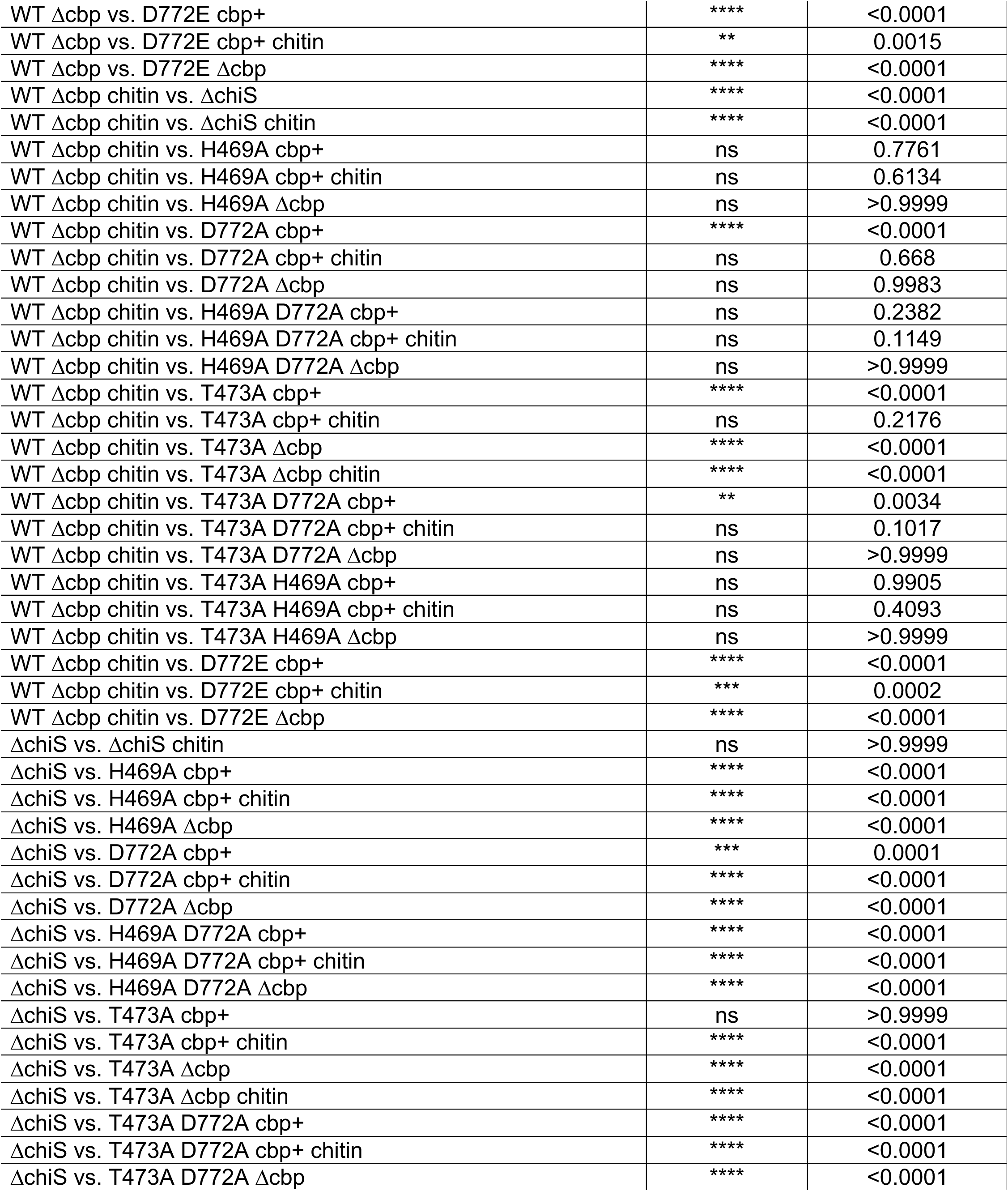
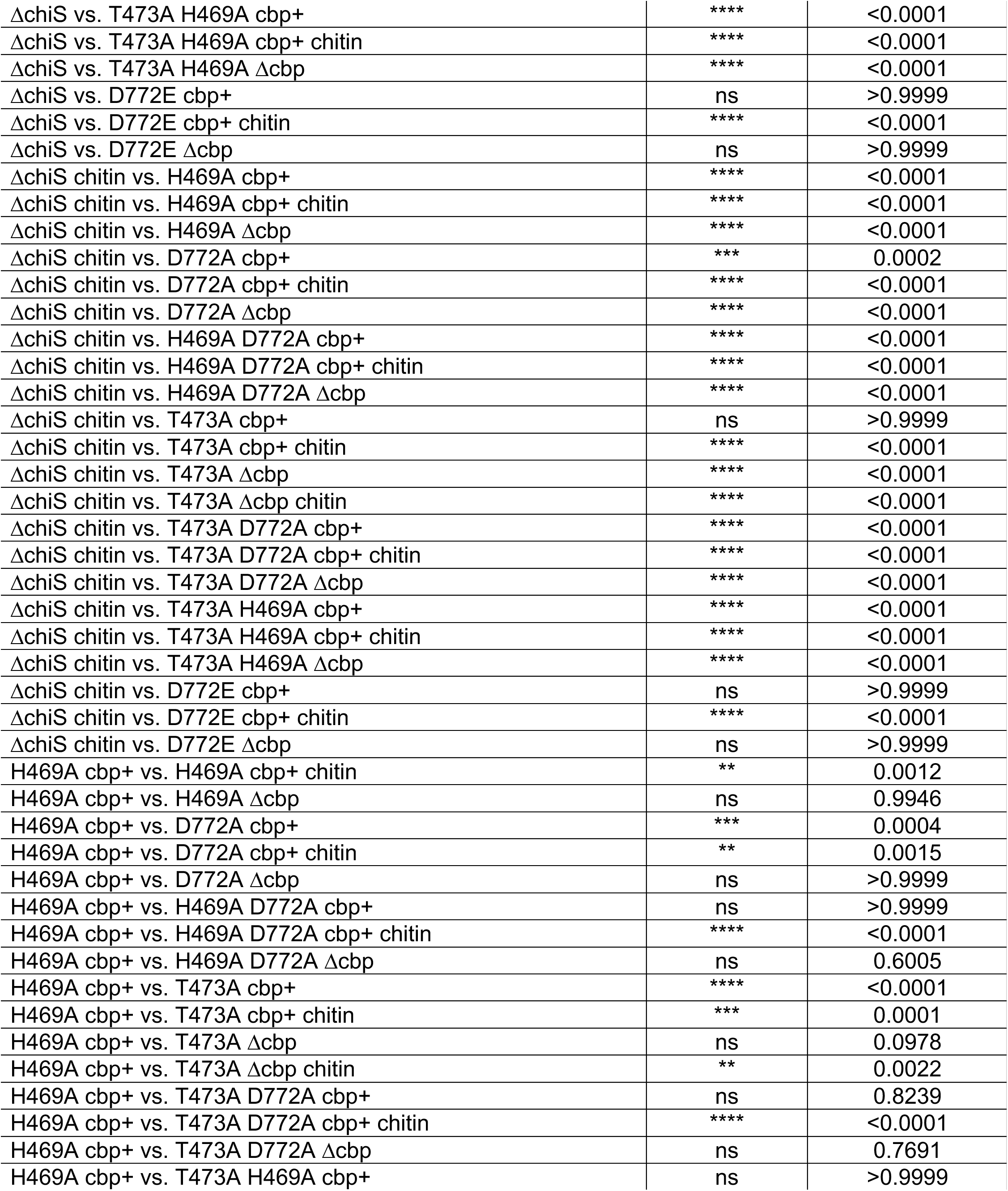
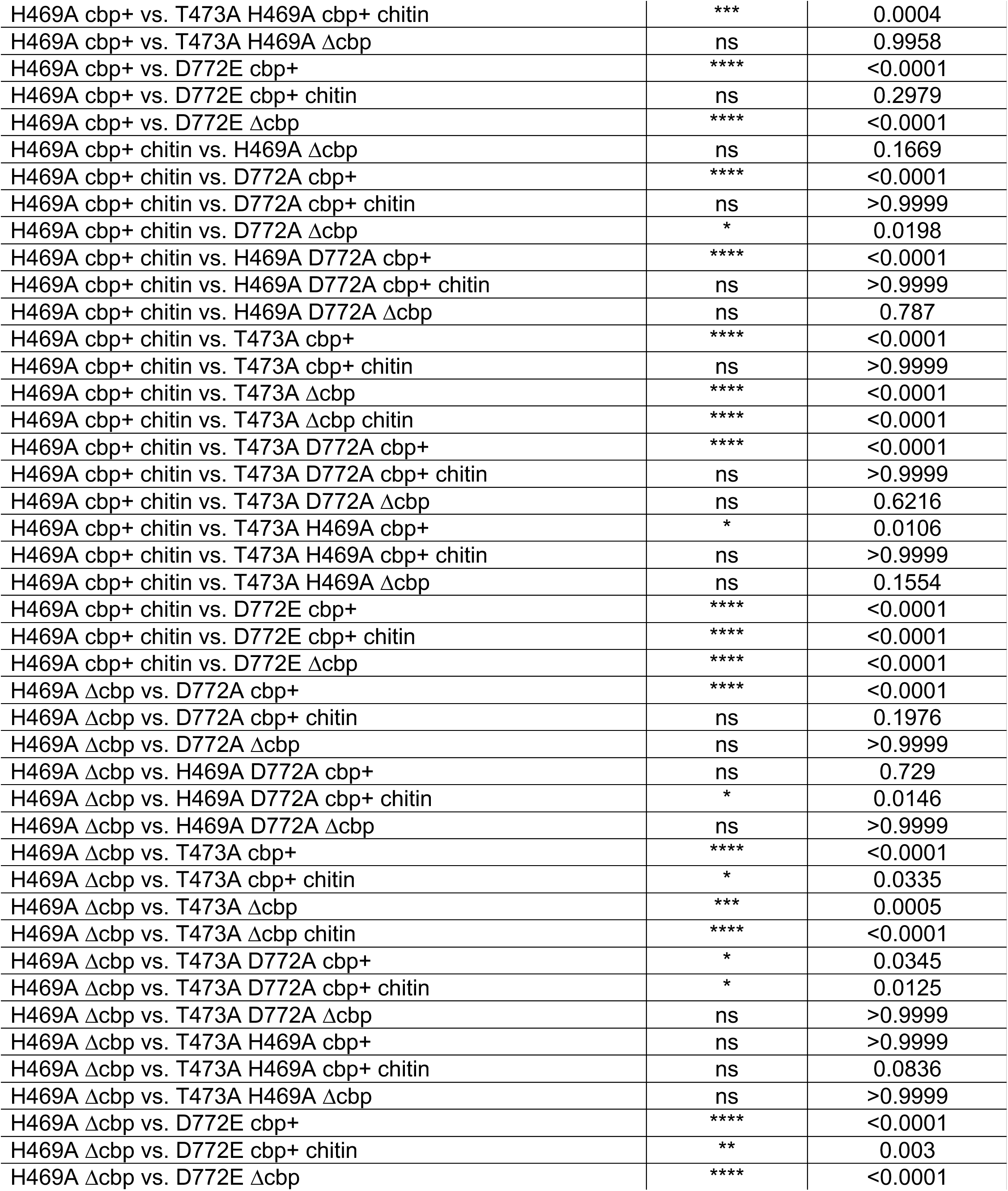
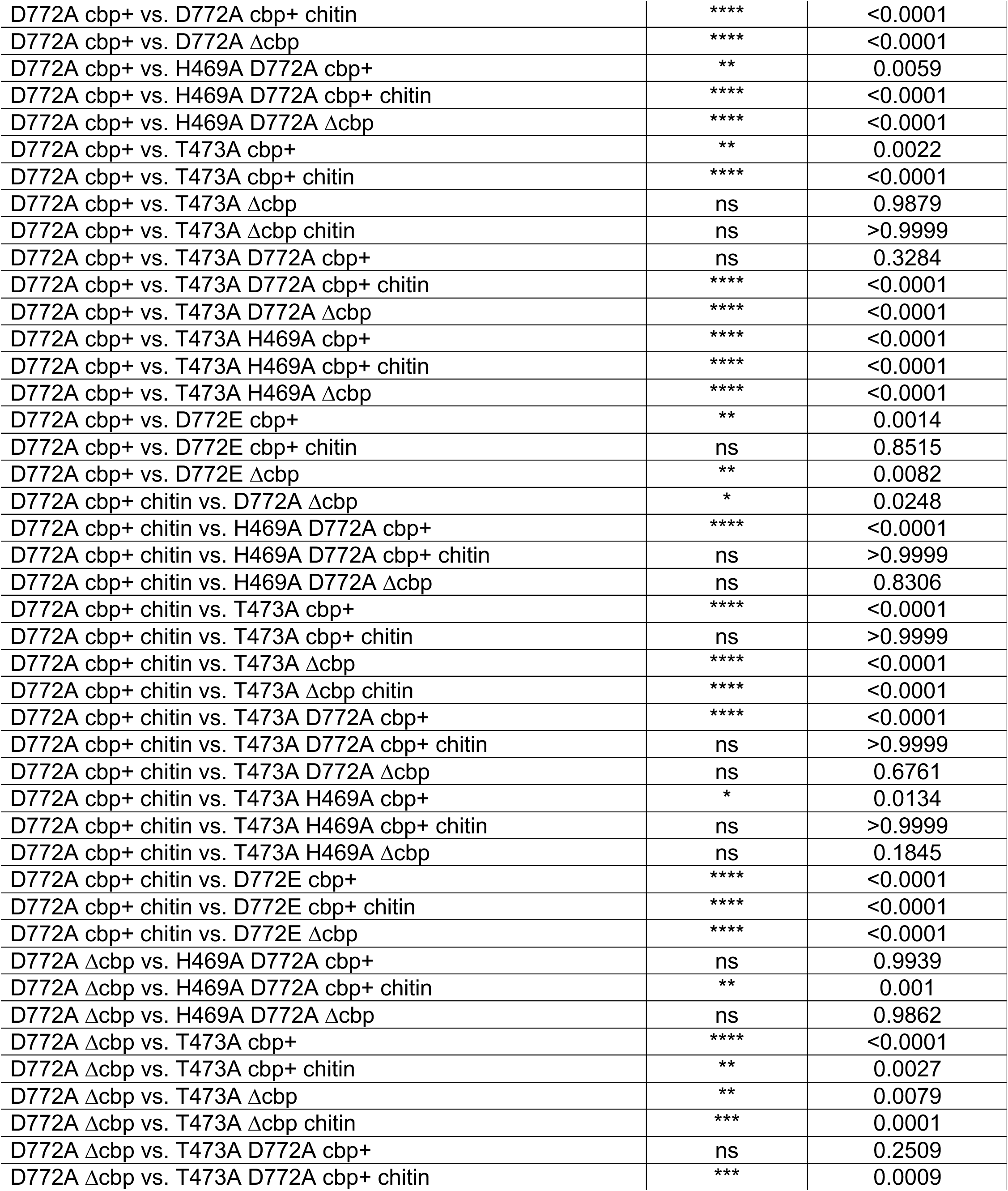
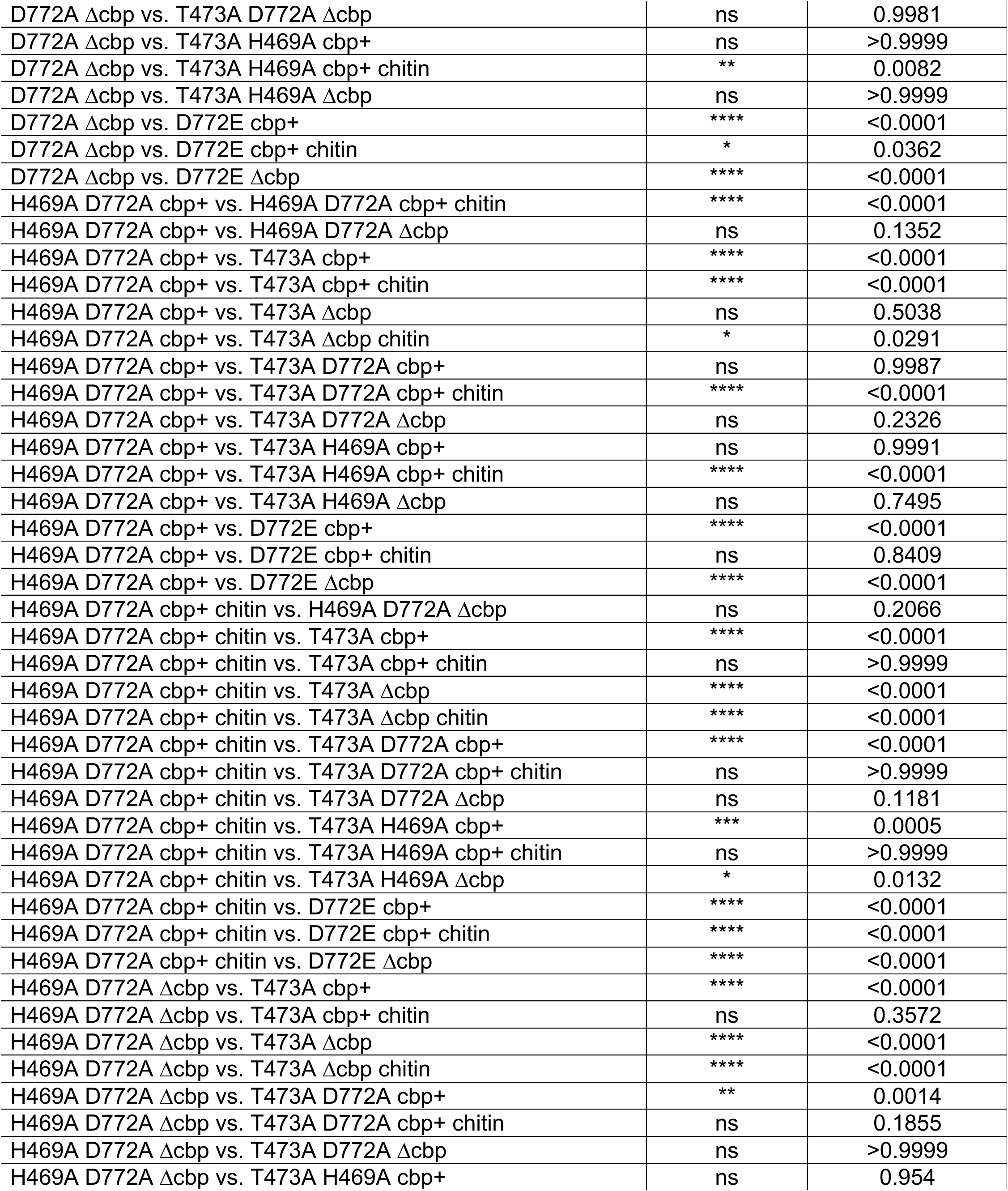
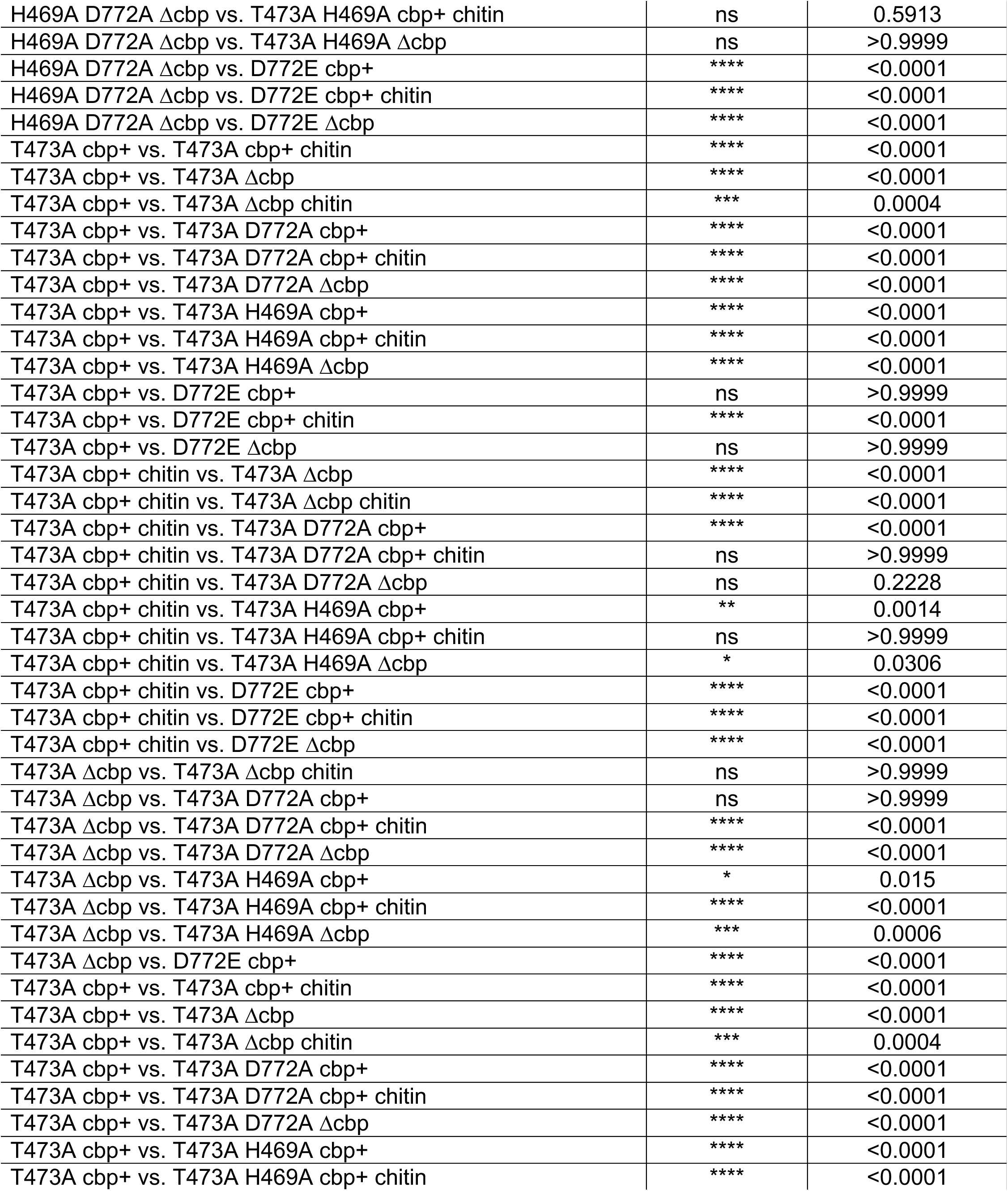
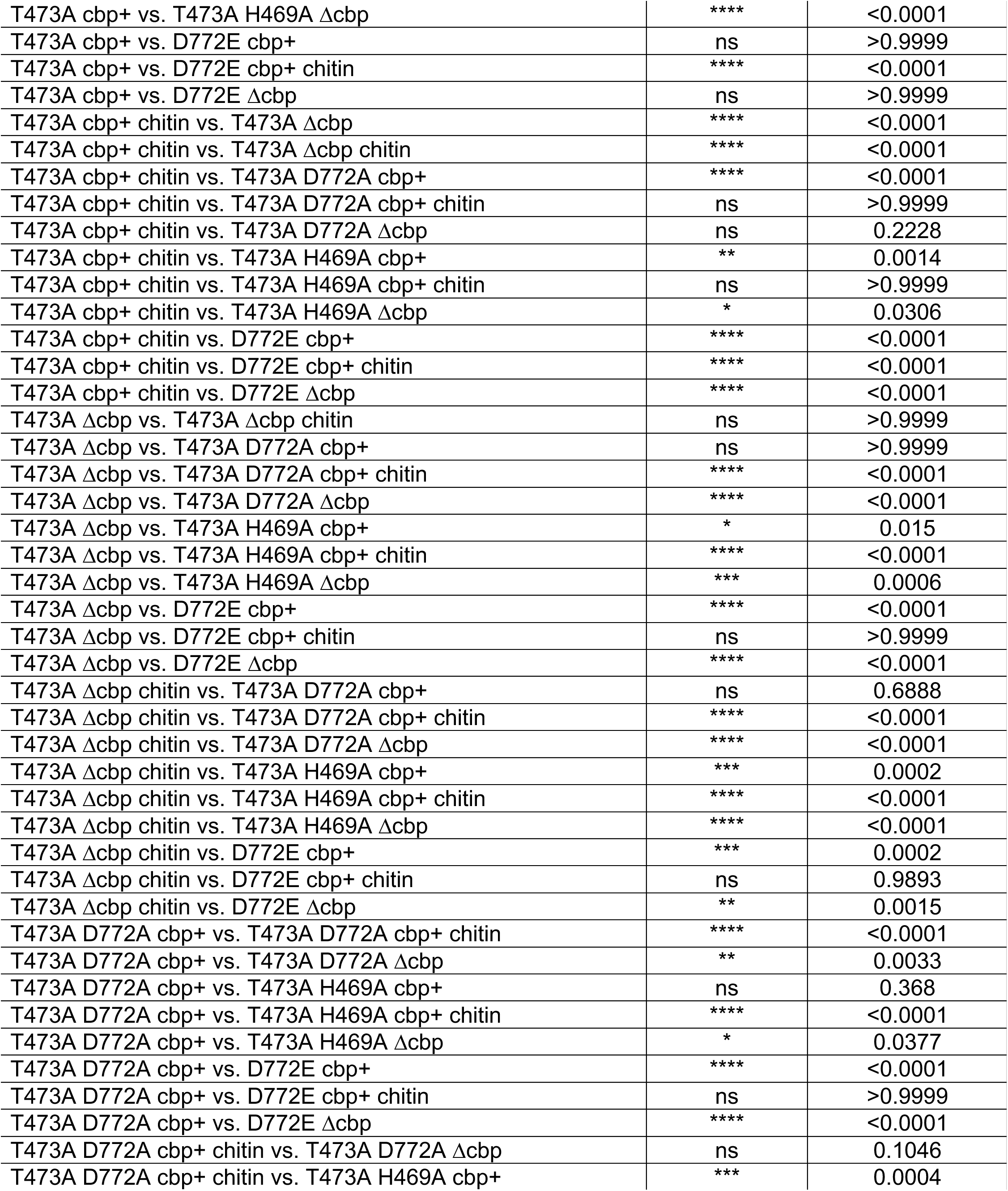
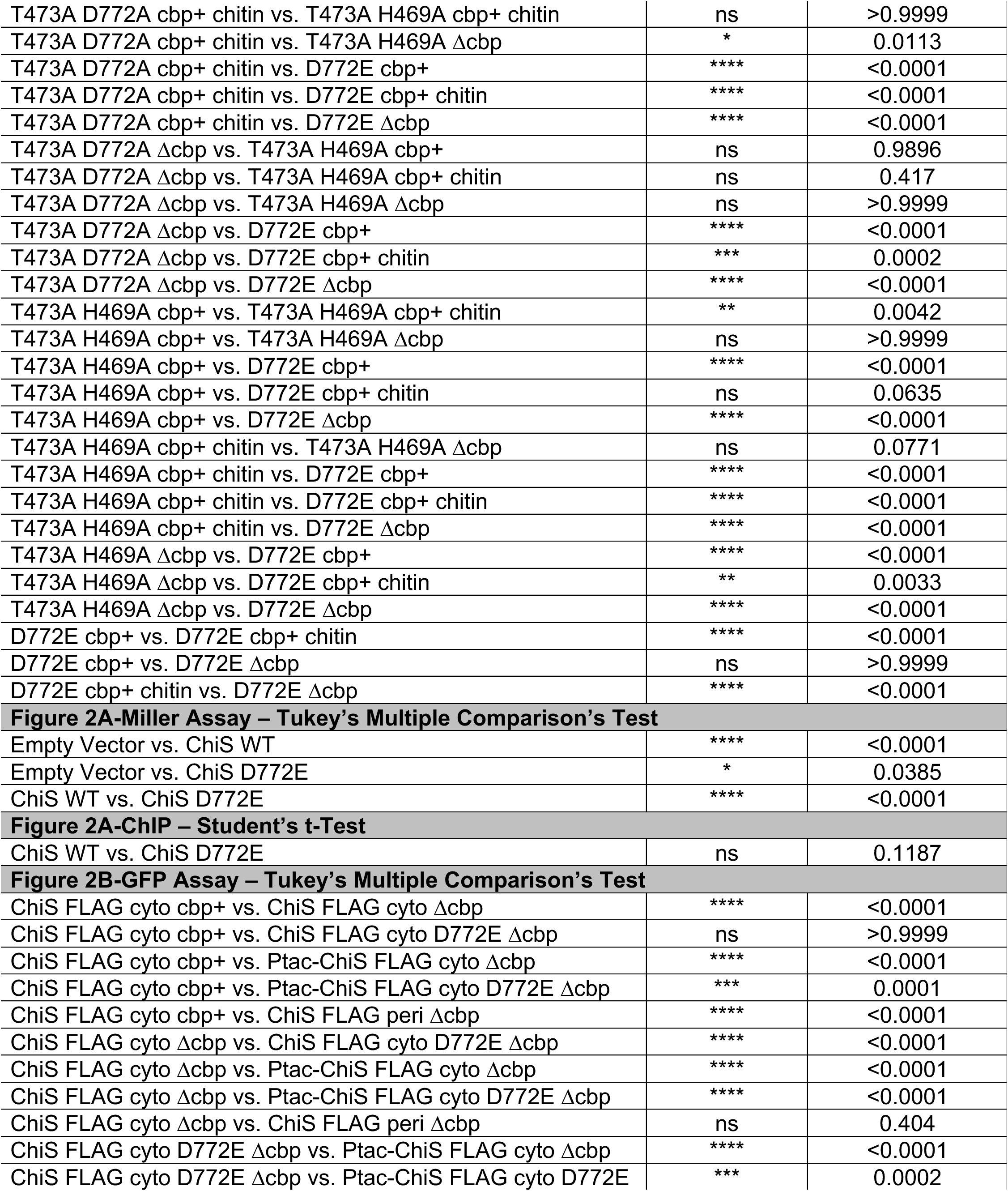
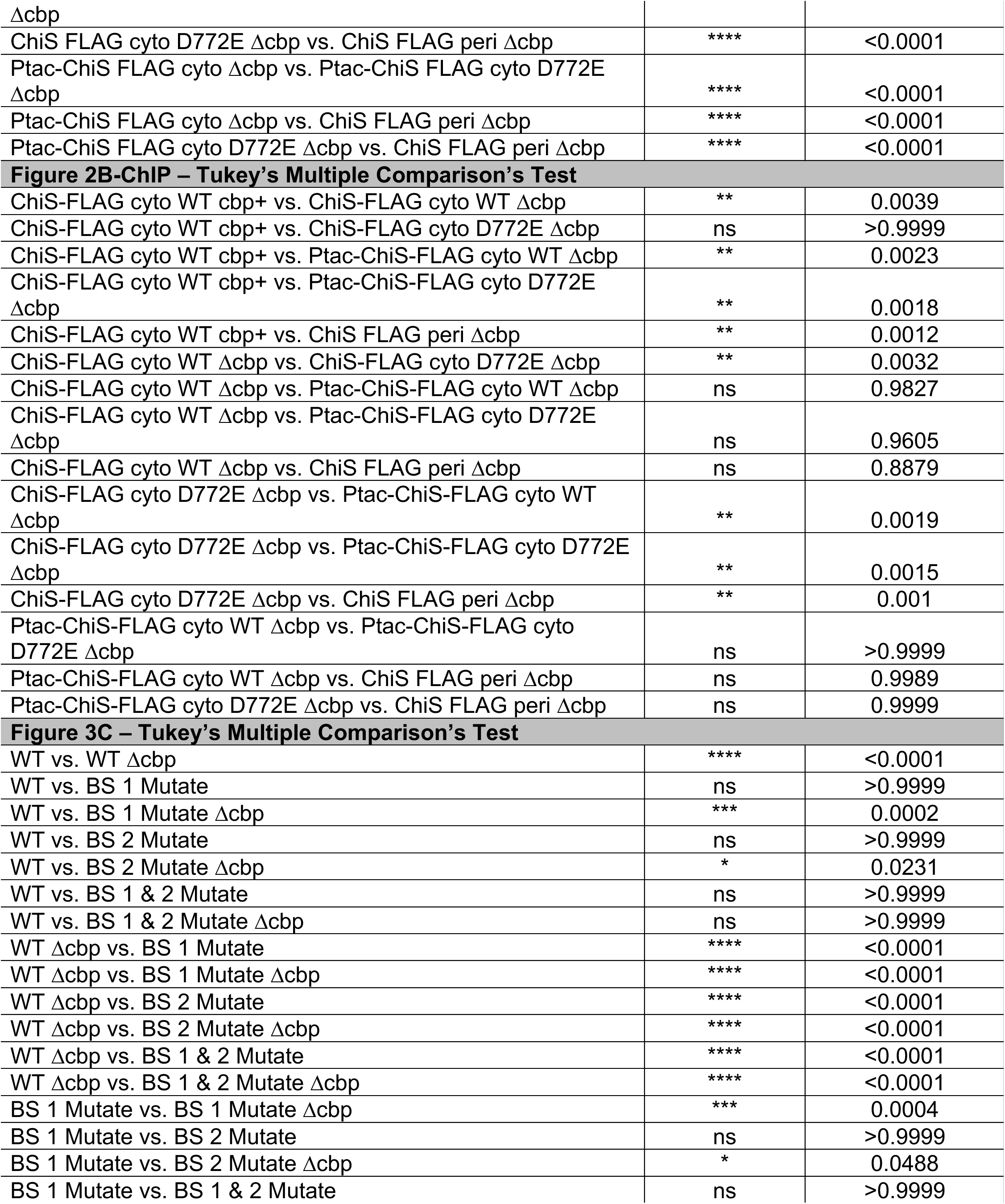
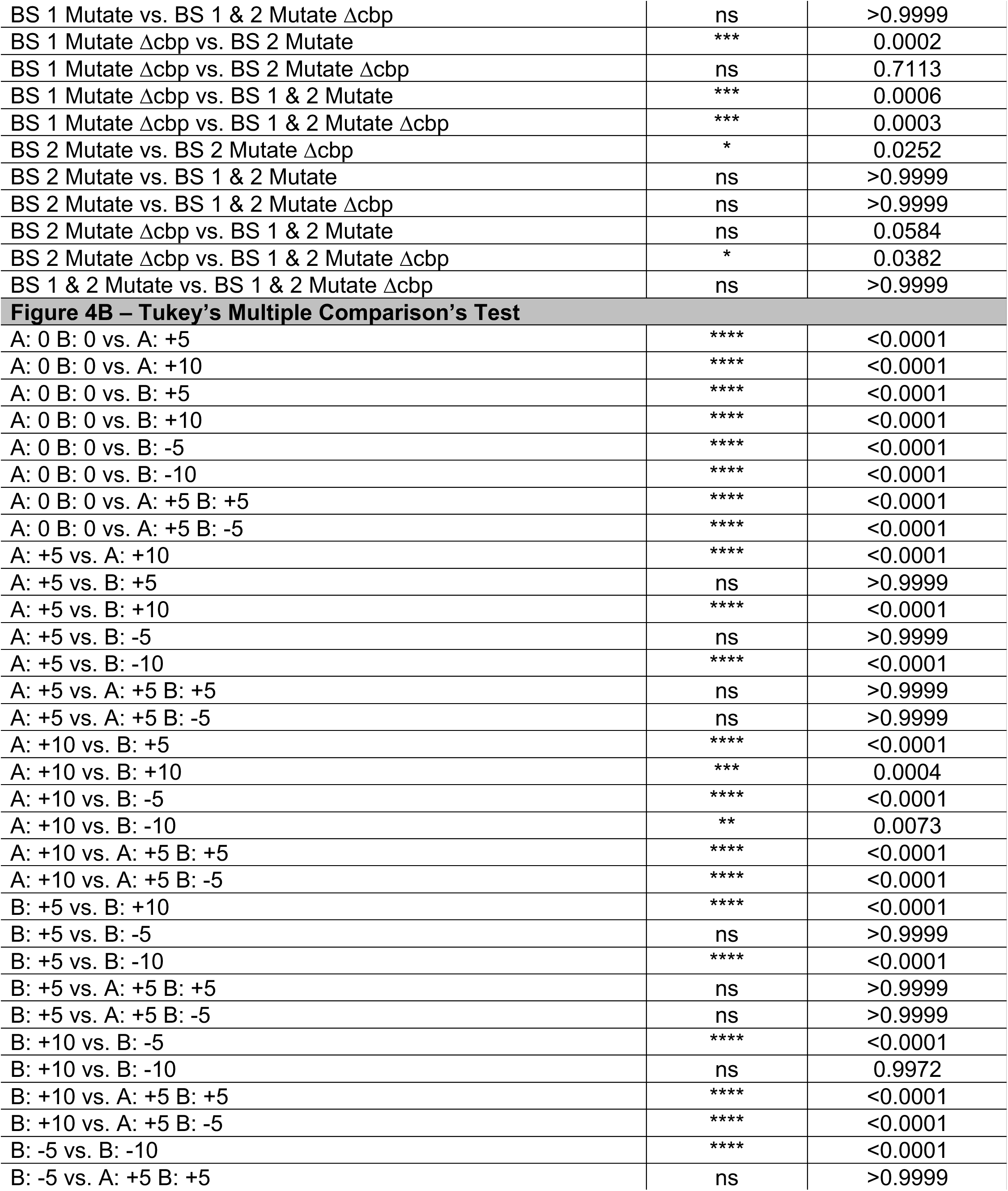
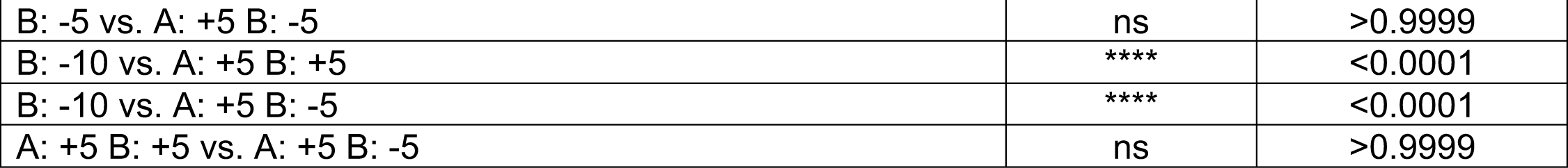
Full list of statistical comparisons made in this manuscript

**Table S2.**
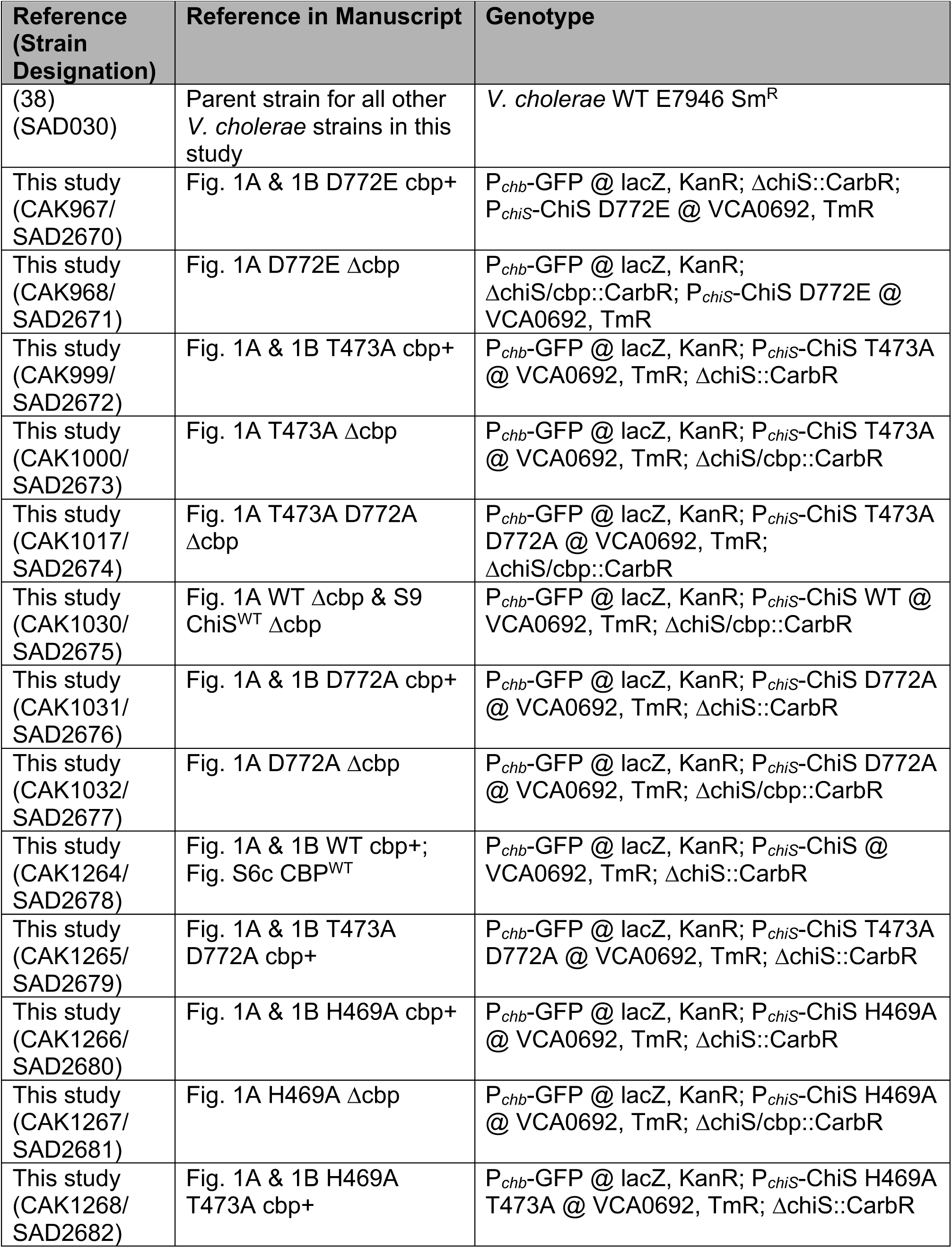
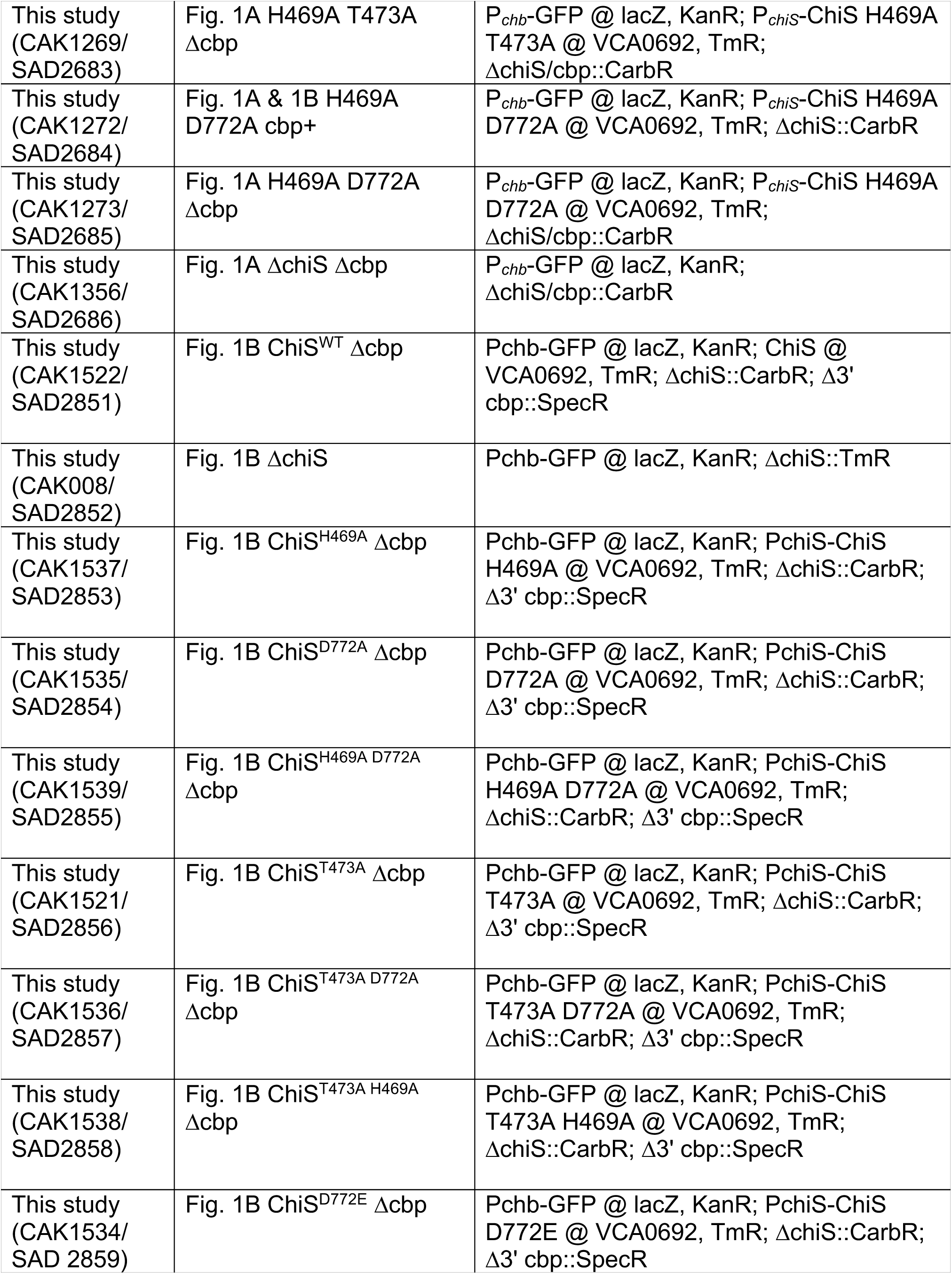
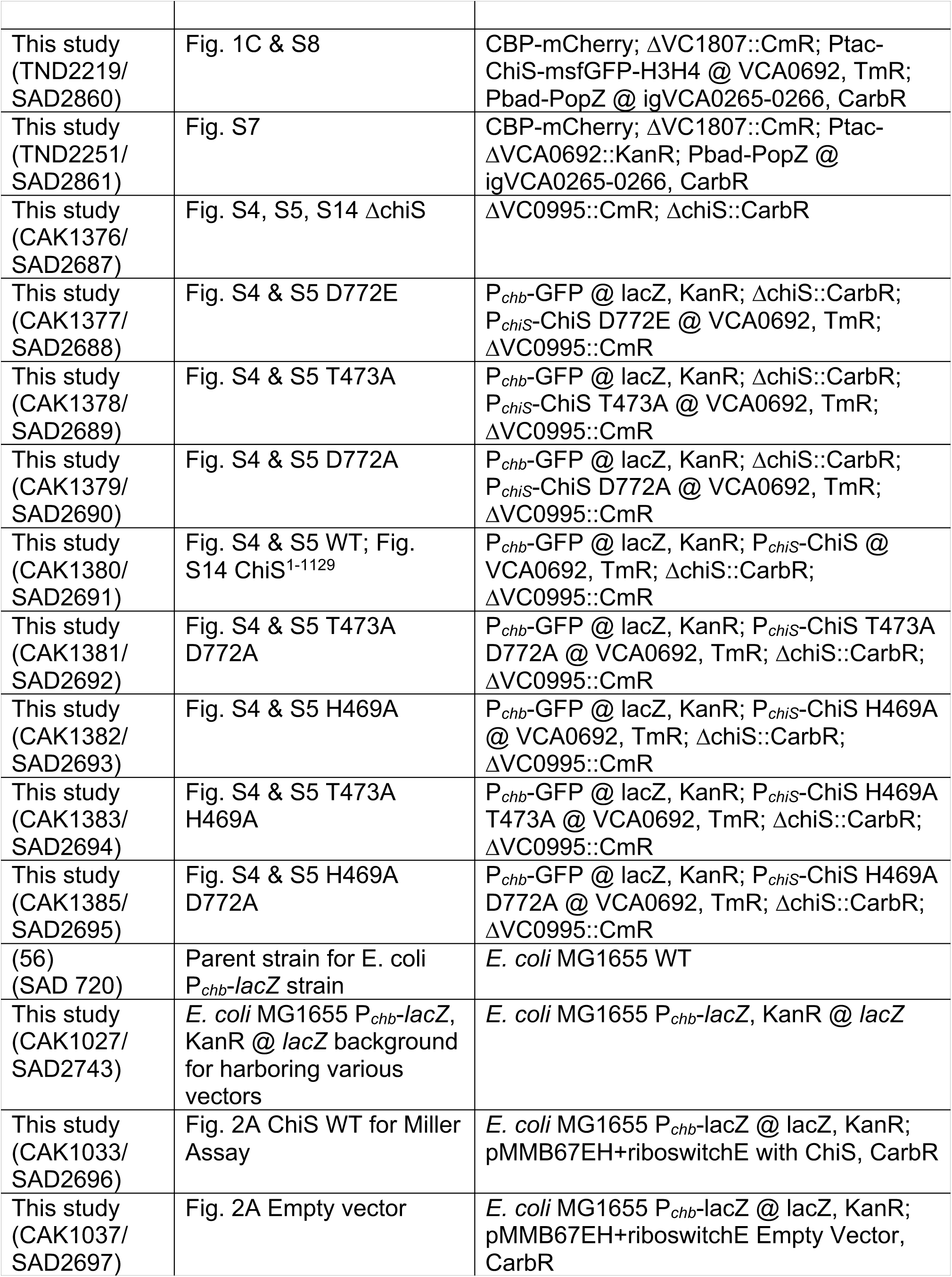
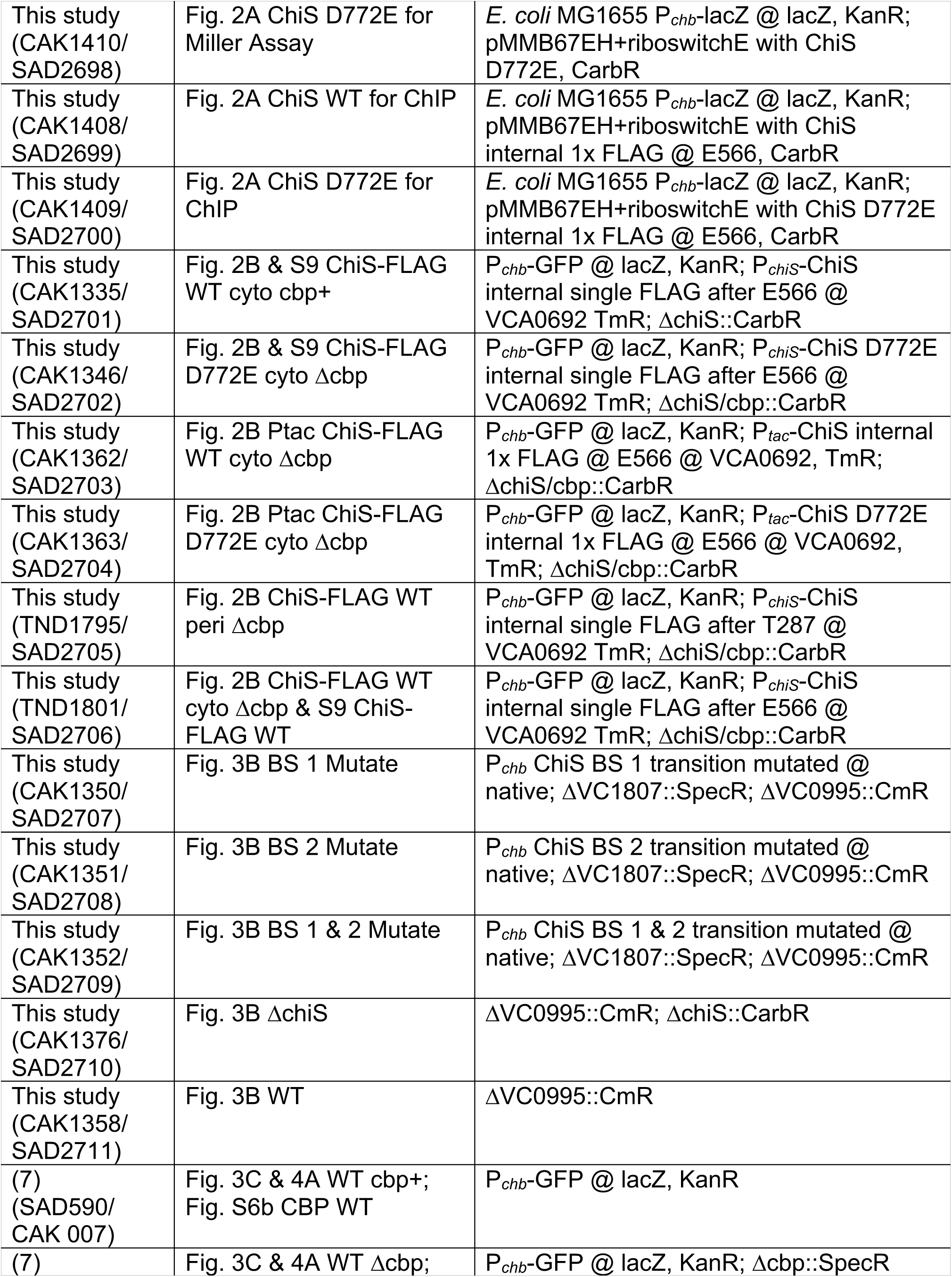
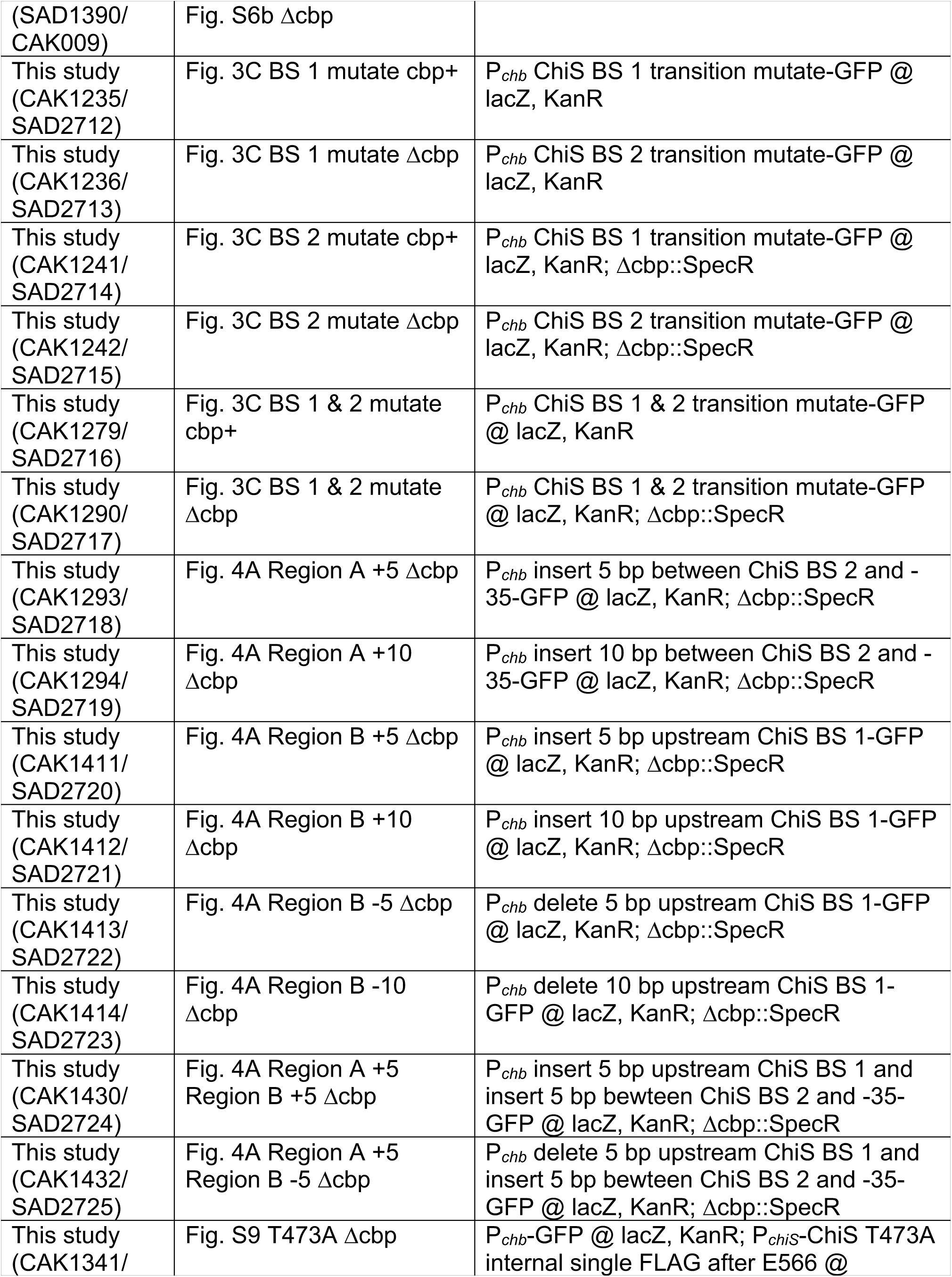
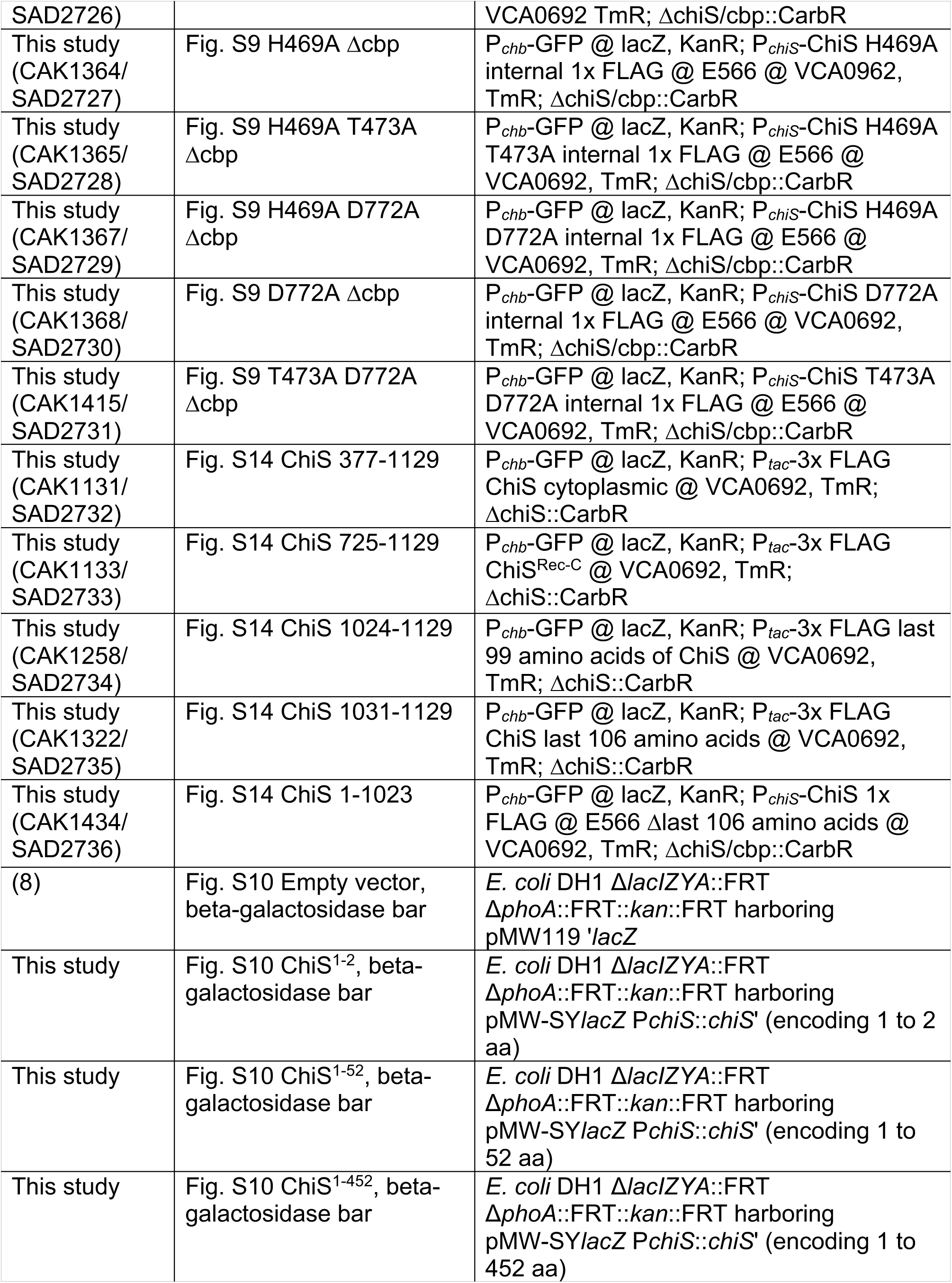
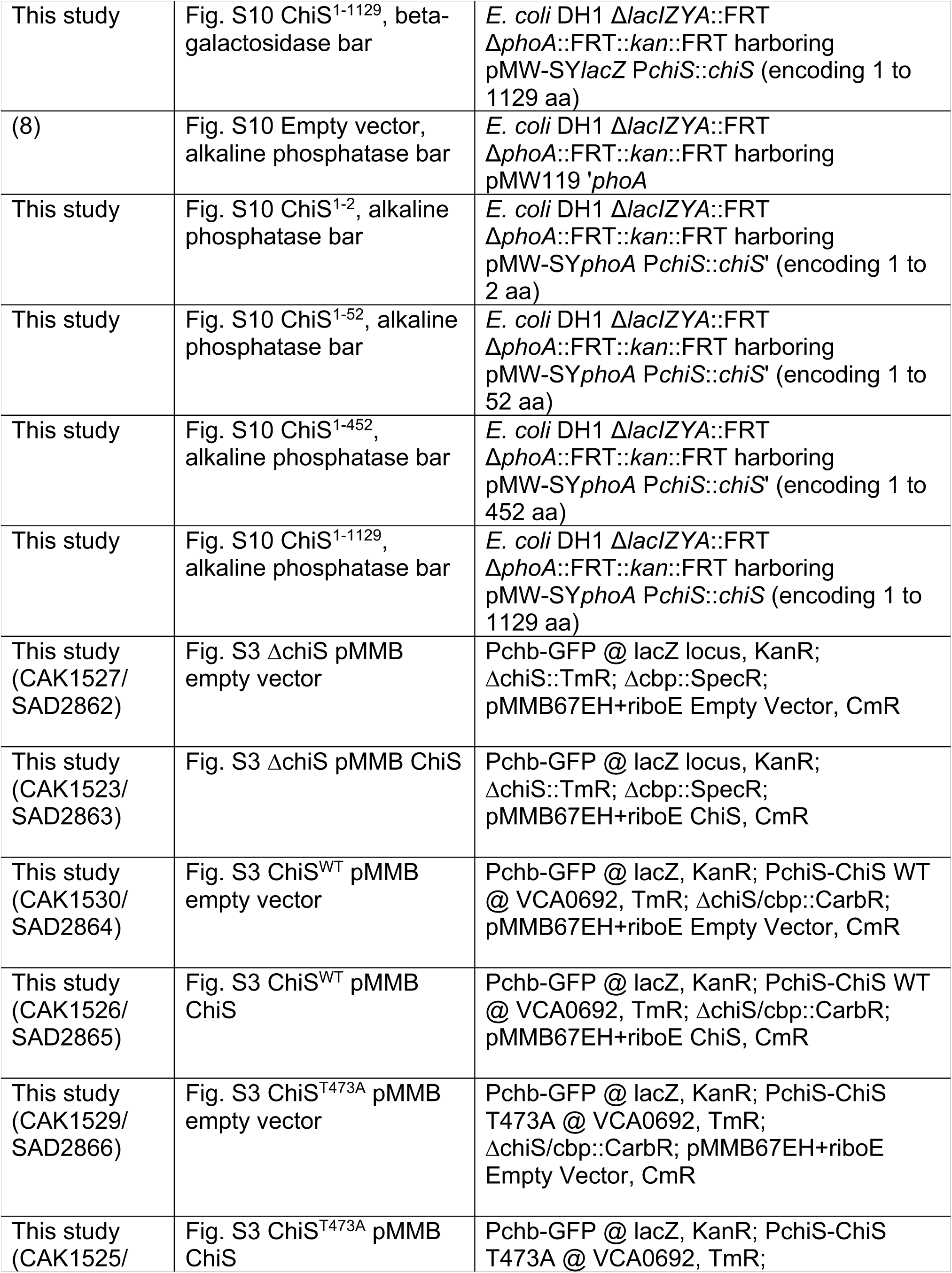
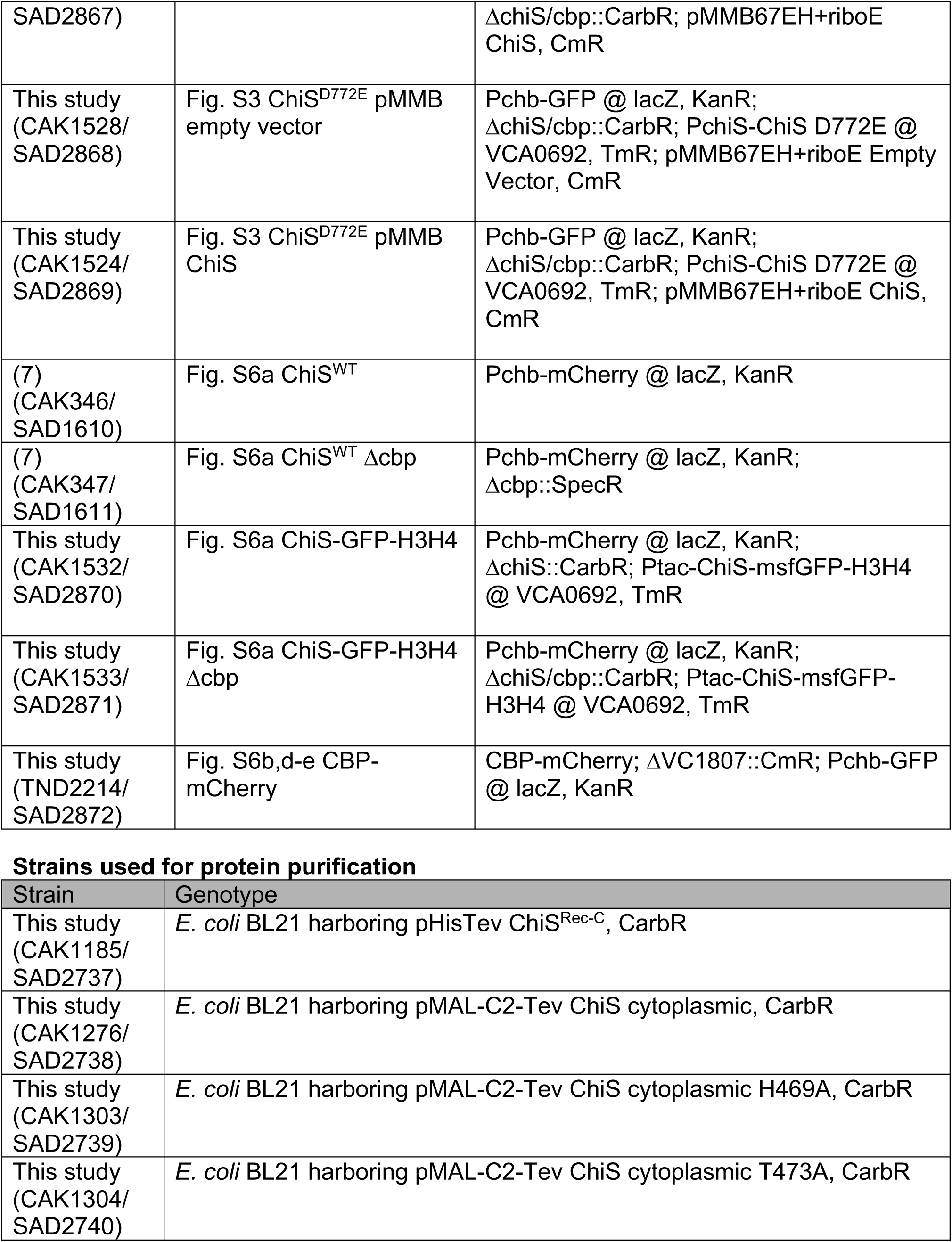
Strains used in this manuscript

**Table S3.**
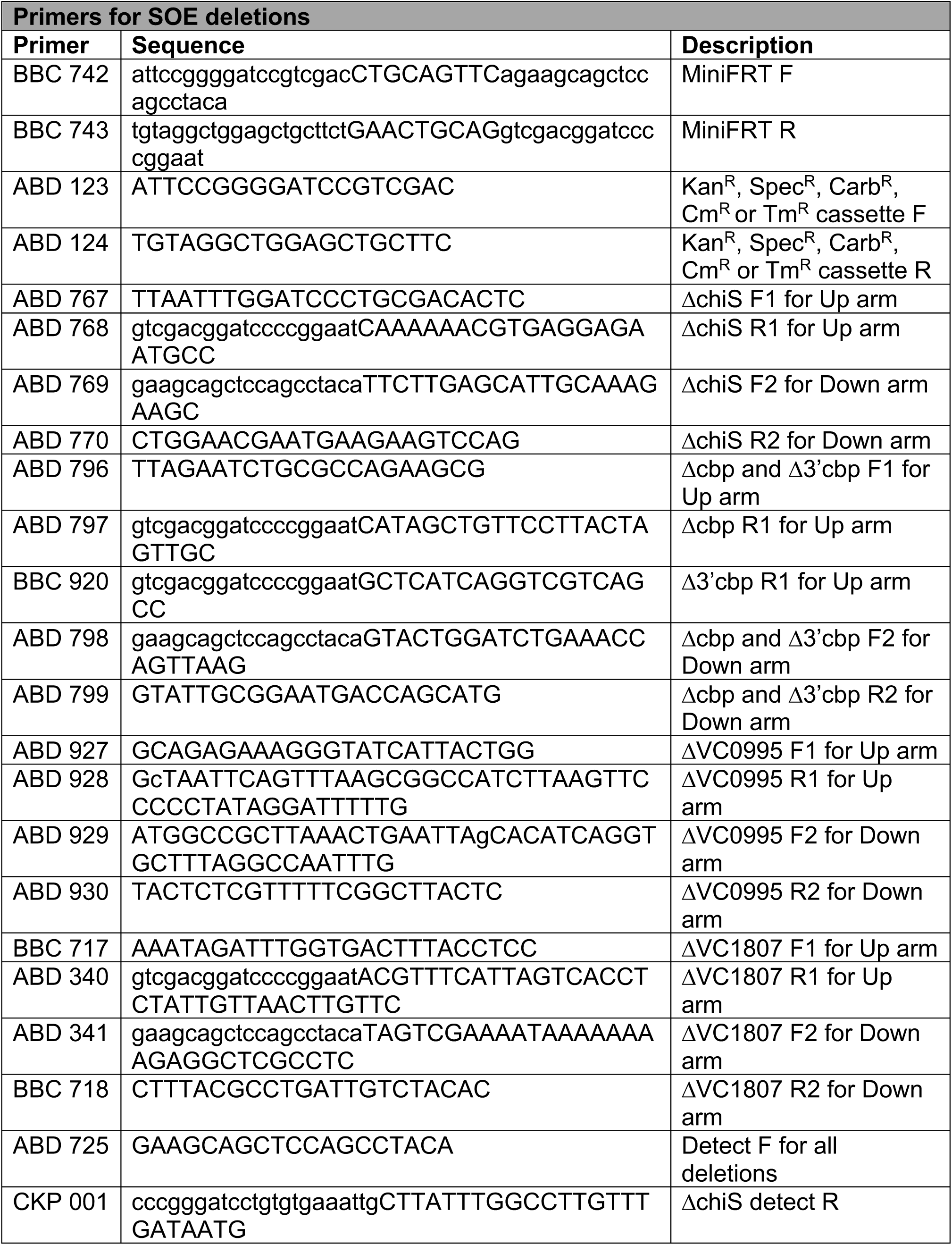
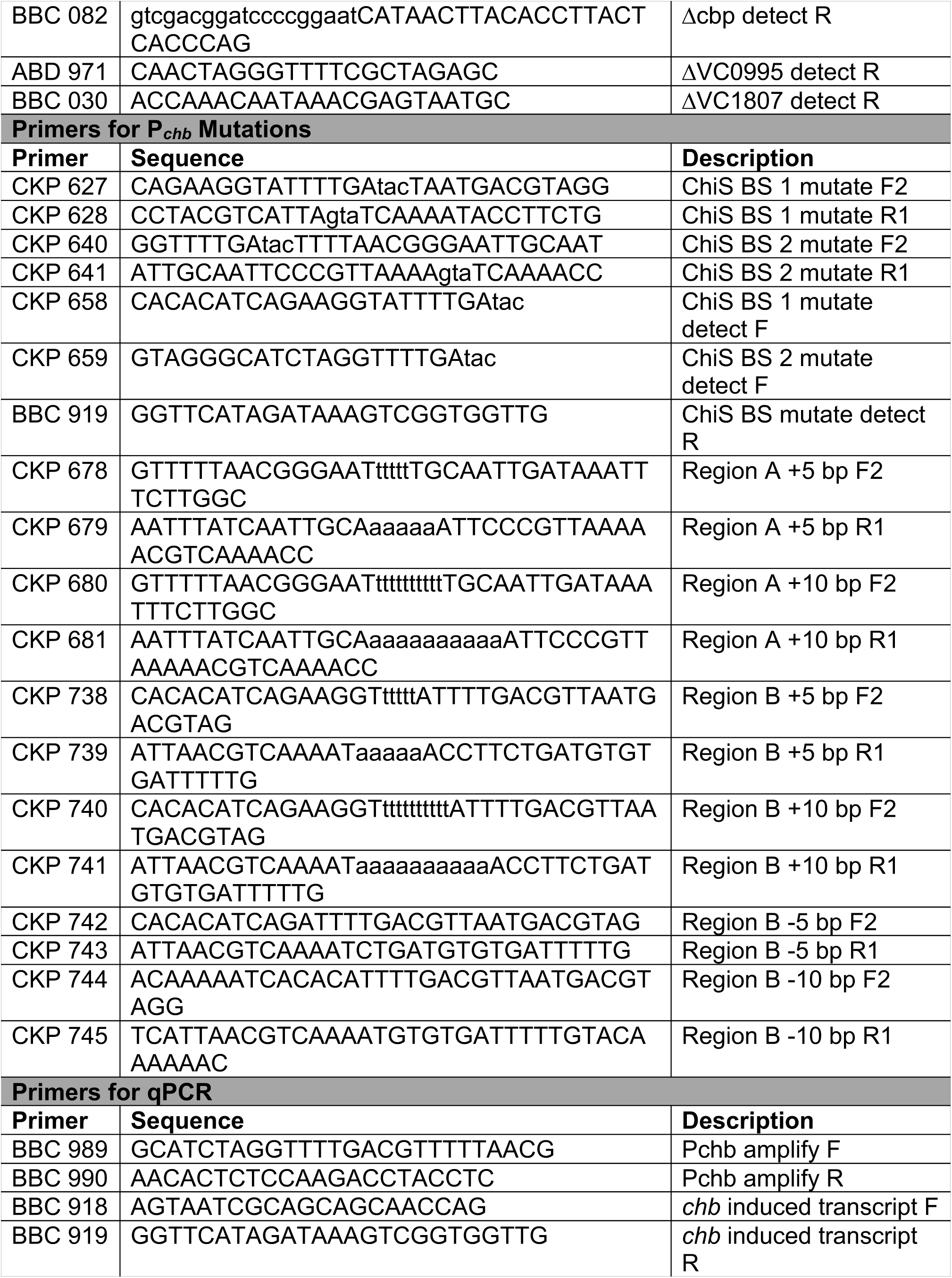
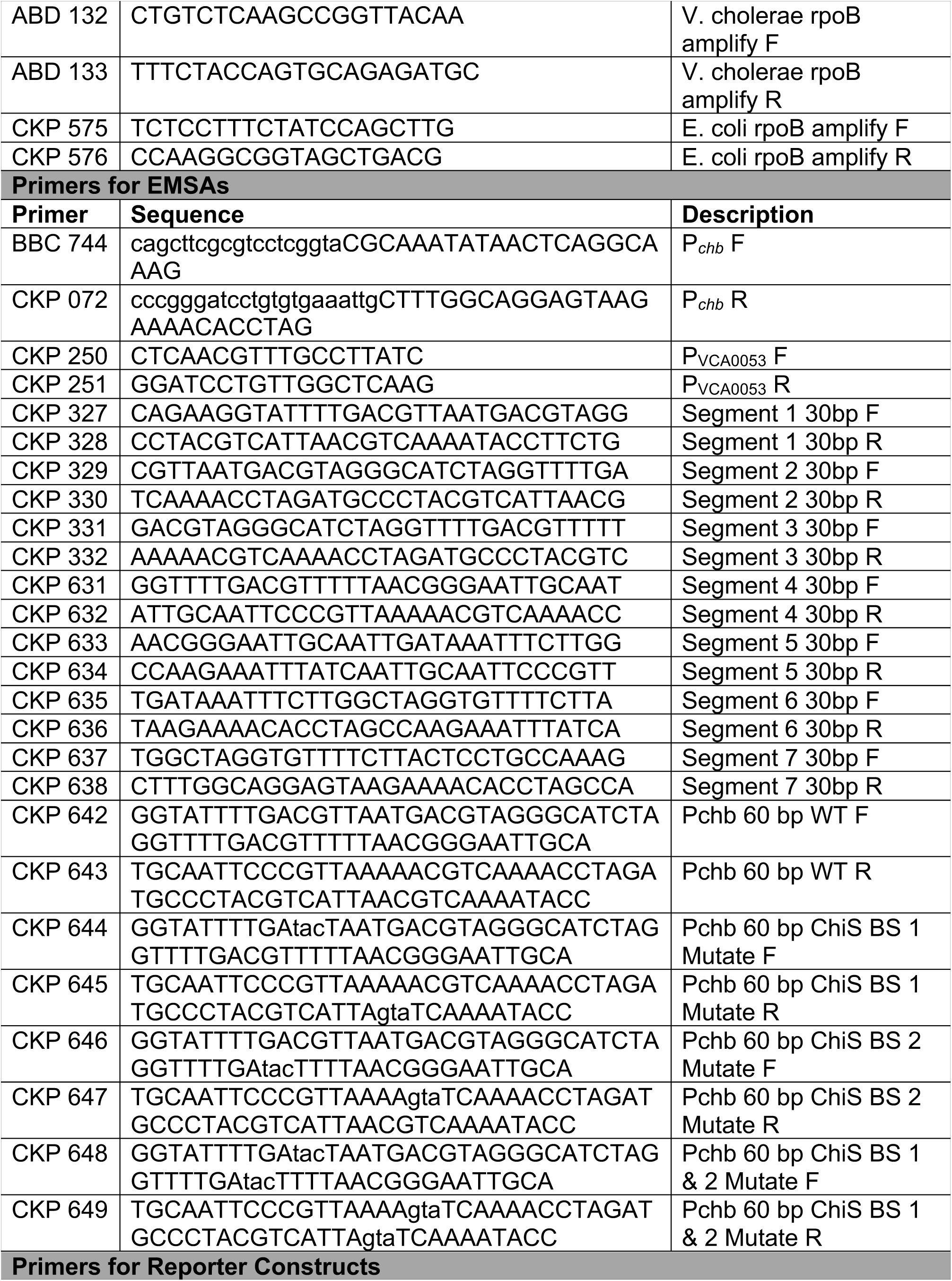
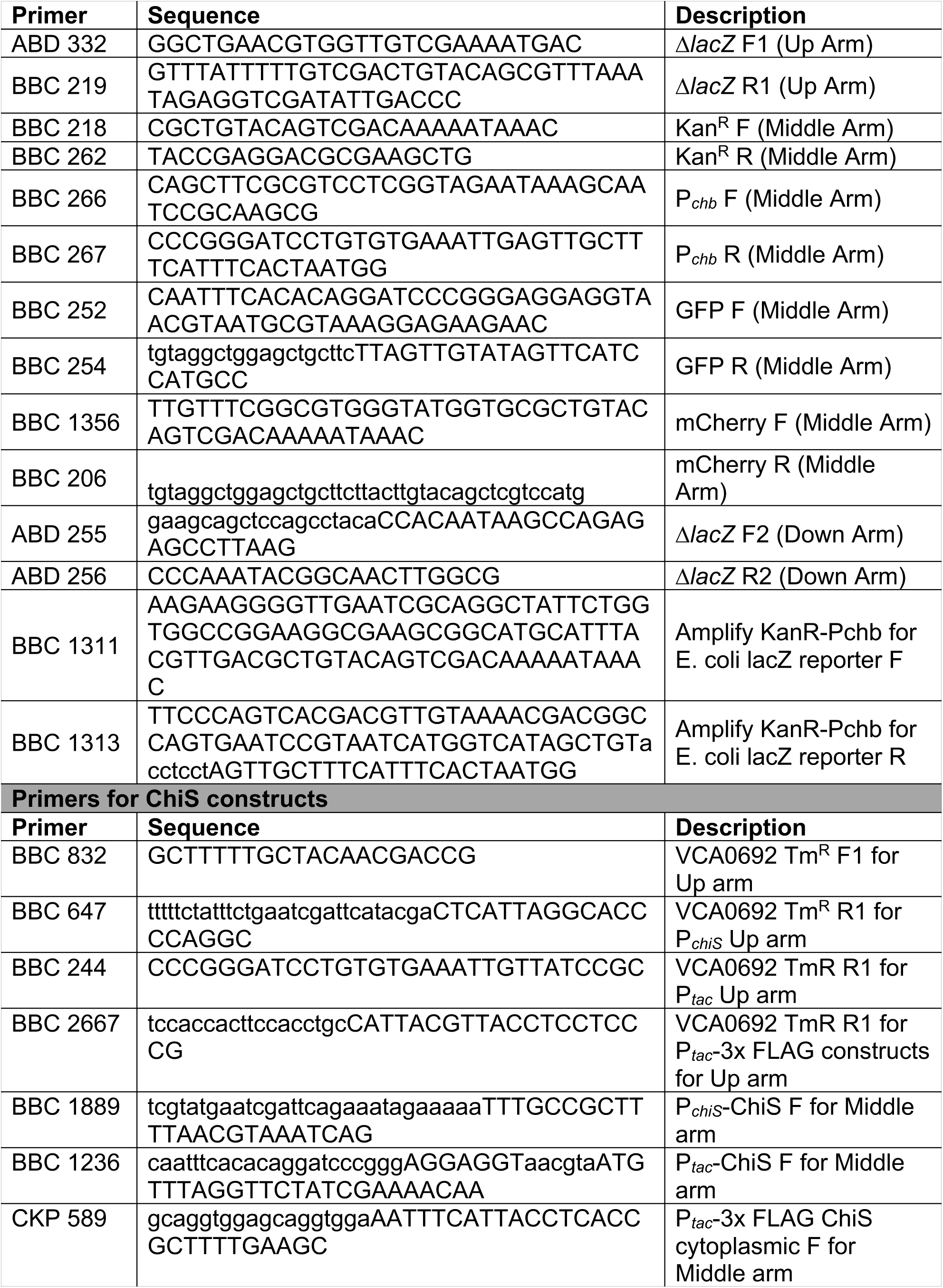
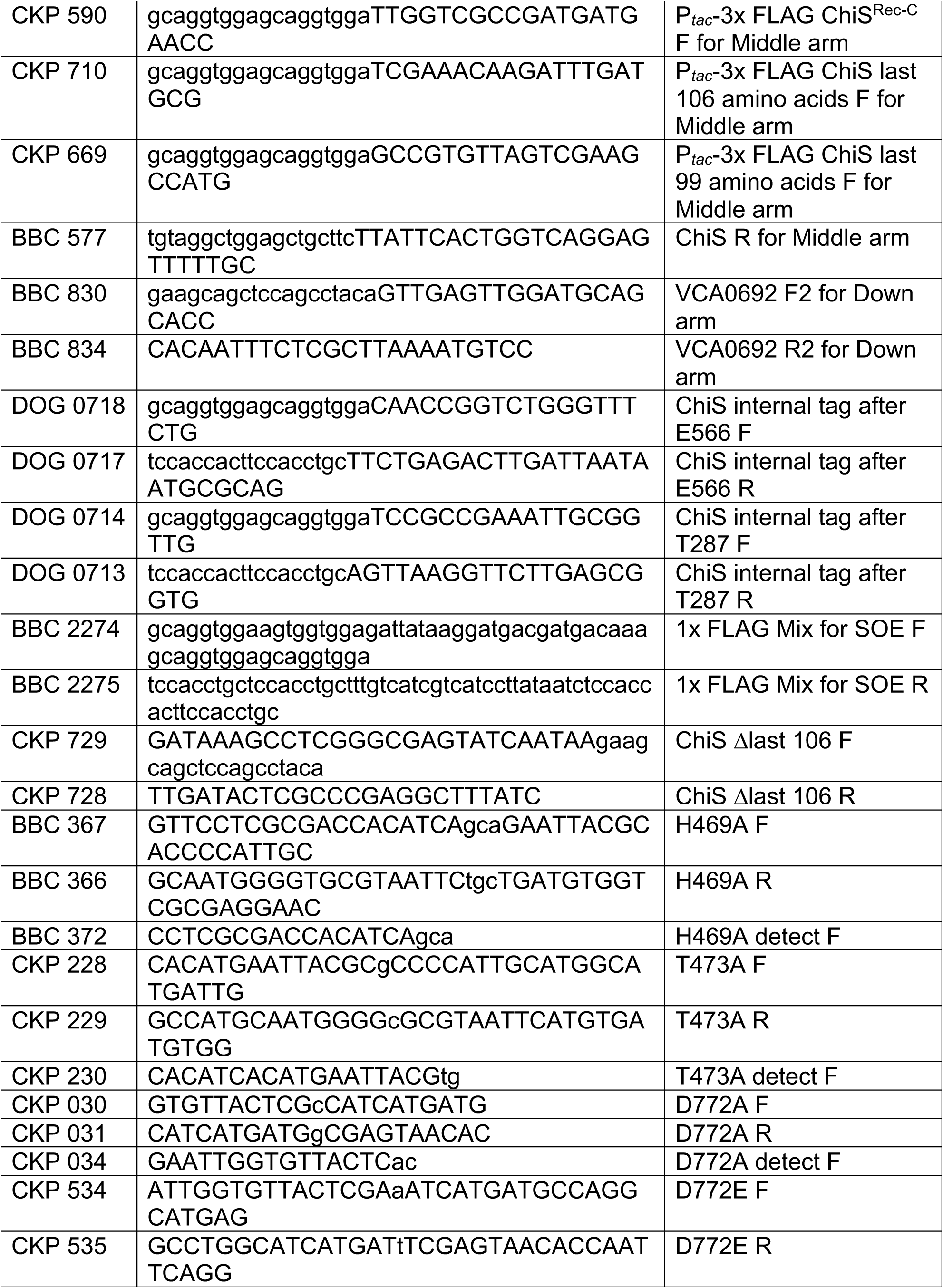
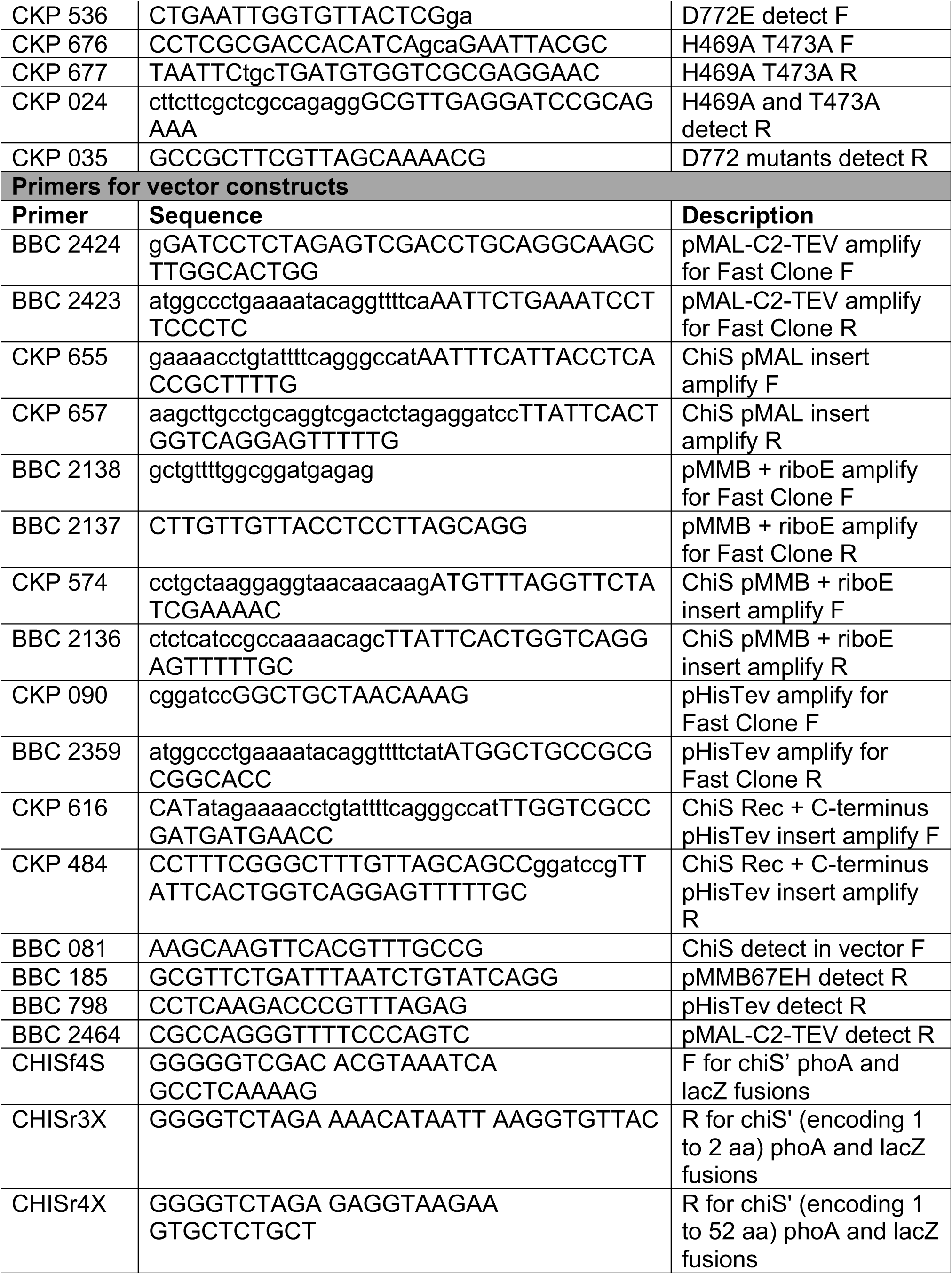
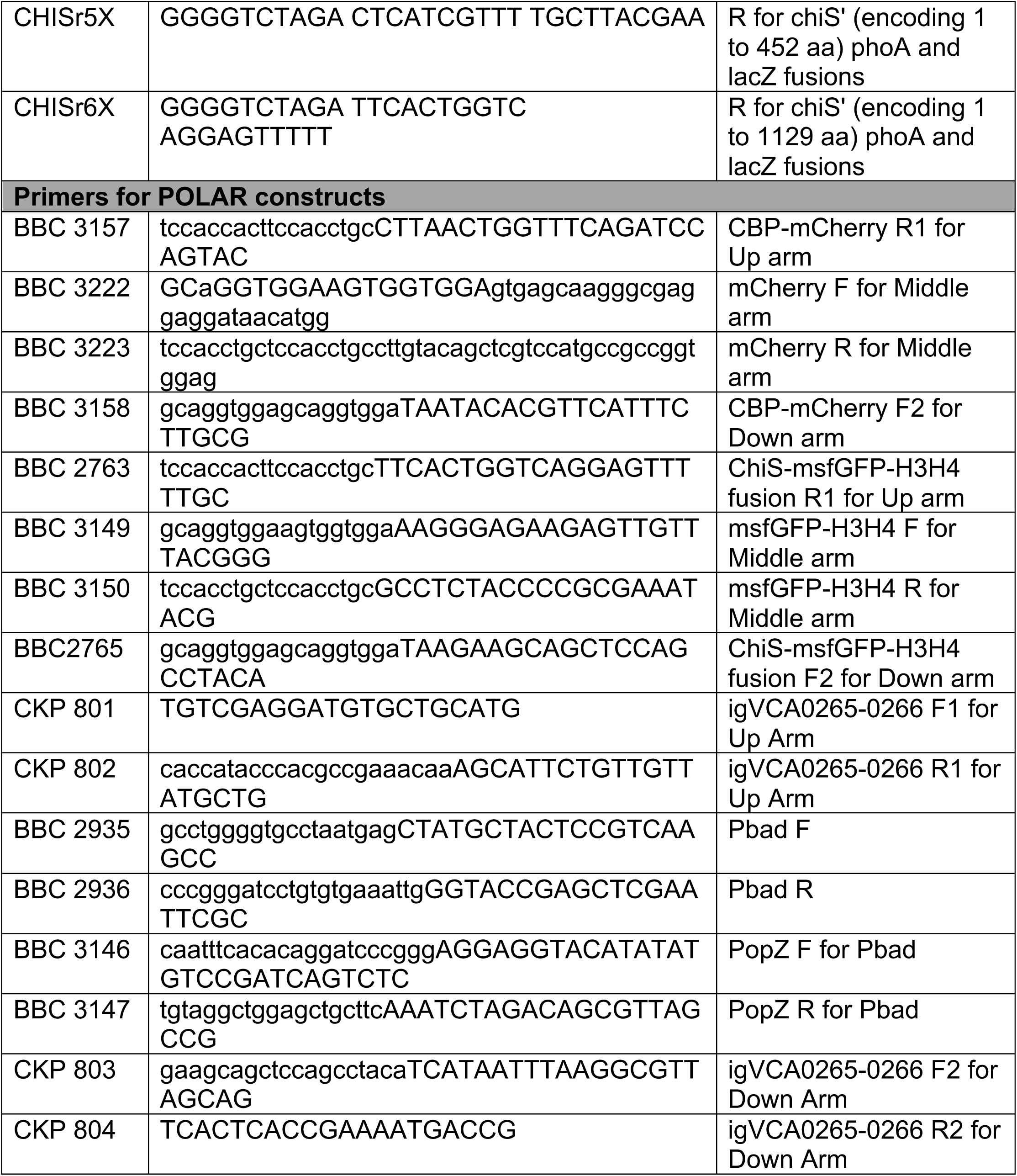
Primers used in this study.

